# Altered Purinergic Signaling and CD8^+^ T Cell Dysregulation in STAT3 GOF Syndrome

**DOI:** 10.1101/2024.12.12.626682

**Authors:** Jose S. Campos Duran, Samir Sayed, Megan C. Dalalo, Andrea A. Mauracher, Montana S. Knight, Peyton E. Conrey, Aaron B. Schultz, Ceire A. Hay, Robert B. Lindell, Christian A. Howard, Eric D. Abrams, Erica G. Schmitt, Martin A. Thelin, Sara Bluestein, Christine M. Seroogy, Tamara C. Pozos, Akaluck Thatayatikom, Ingrid Lundgren, Amelie Gauthier, Scott W. Canna, Helen C. Su, Michael D. Keller, Ottavia M. Delmonte, Lisa R. Forbes Satter, Steven M. Holland, Jenna R. E. Bergerson, Jennifer W. Leiding, Neil Romberg, Alexandra F. Freeman, Alejandro V. Villarino, Mark S. Anderson, Megan A. Cooper, Tiphanie P. Vogel, Sarah E. Henrickson

**Affiliations:** Division of Allergy and Immunology, Department of Pediatrics, Children’s Hospital of Philadelphia, Philadelphia, PA, USA; Perelman School of Medicine, University of Pennsylvania, Philadelphia, PA, USA; Institute for Immunology and Immune Health, University of Pennsylvania Perelman School of Medicine, Philadelphia, PA, USA; Department of Biomedical and Health Informatics, Children’s Hospital of Philadelphia, Philadelphia, PA, USA; Department of Microbiology and Immunology, Miller School of Medicine, University of Miami, Miami, FL, USA; Division of Critical Care Medicine, Department of Anesthesia and Critical Care, Children’s Hospital of Philadelphia, Philadelphia, PA, USA; Department of Pediatrics, Division of Rheumatology and Immunology, Washington University School of Medicine, St. Louis, Missouri, USA; Department of Pathology and Immunology, Washington University School of Medicine, St. Louis, Missouri, USA; Diabetes Center, University of California, San Francisco, San Francisco, CA, USA; Department of Pediatric Endocrinology, University of California San Francisco, San Francisco, CA, USA; Atlanta Allergy & Asthma, Atlanta, GA, USA; University of Wisconsin School of Medicine and Public Health, Madison, WI, USA; Clinical Immunology, Children’s Minnesota, Minneapolis, MN, USA; Pediatric Rheumatology/ Immunology, AdventHealth for Children, Orlando, FL, USA; Infectious Diseases, St. Luke’s Children’s Hospital, Boise, ID, USA; Department of Allergy and Immunology, CHU de Québec-CHUL, Laval University Hospital Center, Laval University, Quebec City, CA; Division of Rheumatology, Immune Dysregulation Program, Children’s Hospital of Philadelphia, Philadelphia, PA, USA; Laboratory of Clinical Immunology and Microbiology, National Institute of Allergy and Infectious Diseases, National Institutes of Health, Bethesda, MD, USA; Center for Cancer & Immunology Research, Children’s National Hospital, Washington, DC, USA; Division of Allergy and Immunology, Children’s National Hospital, Washington, DC, USA; GW Cancer Center, George Washington University School of Medicine, Washington, DC, USA; Department of Pediatrics, Baylor College of Medicine and William T Shearer Center for Human Immunobiology, Texas Children’s Hospital, Houston, TX, USA; Division of Allergy and Immunology, Department of Pediatrics, Johns Hopkins University, Baltimore, MD, USA; Institute for Clinical and Translational Research and Cancer and Blood Disorders Institute, Johns Hopkins All Children’s Hospital, St. Petersburg, FL, USA; Department of Medicine, University of California, San Francisco, San Francisco, CA, USA; Department of Microbiology and Immunology, University of California, San Francisco, San Francisco, CA, USA; Department of Microbiology, Perelman School of Medicine, University of Pennsylvania, Philadelphia, PA, USA

## Abstract

Signal transduction downstream of activating stimuli controls CD8^+^ T cell biology, however these external inputs can become uncoupled from transcriptional regulation in Primary Immune Regulatory Disorders (PIRD). Gain-of-function (GOF) variants in STAT3 amplify cytokine signaling and cause a severe PIRD characterized by early onset autoimmunity, lymphoproliferation, recurrent infections, and immune dysregulation. In both primary human and mouse models of STAT3 GOF, CD8^+^ T cells have been implicated as pathogenic drivers of autoimmunity. The molecular mechanisms by which STAT3 GOF variants drive this pathology remain unclear. We found that naïve CD8^+^ T cells have an increased capacity for IFN-γ and TNF-α secretion. Given this dysregulation of CD8^+^ T cell function, we evaluated changes in immunoregulatory pathways and found evidence of dysregulated purinergic signaling via high dimensional immune profiling, single-cell RNA sequencing, and functional assessment. Specifically, while expression of CD39, which transforms ATP to AMP, was increased on CD8^+^ T cells from patients with STAT3 GOF, downstream purinergic family members, CD73 and the adenosine receptor, A_2A_R, were downregulated, impairing the potential to produce or sense inhibitory adenosine. Patients with STAT3 GOF can be clinically treated with JAK inhibitors and this partially normalized naïve CD8^+^ T cell dysregulation, including aberrant cytokine production. The extent of normalization of cytokine secretion scaled with normalization of CD73 and A_2A_R. This suggests that a dysregulated purinergic signaling axis plays an important role in CD8^+^ T cell dysregulation in STAT3 GOF, which may have implications for other inflammatory disorders with amplified STAT signaling.

**One Sentence Summary:** Amplified STAT3 signaling in CD8^+^ T cells from STAT3 GOF patients alters purinergic signaling and dysregulates CD8^+^ T cell function.

## INTRODUCTION

Rare, monogenic inborn errors of immunity (IEI) provide insight into human immune function via specific alterations of key immune pathways. The approximately 500 currently defined IEIs have a broad range of effects, which include changing the relative proportion and/or function of cellular and humoral components of the immune system (*1*, *2*). In a subset of IEI, termed Primary Immune Regulatory Disorders (PIRD), genetic variants in more than 100 different genes can alter the regulation of immune cell differentiation and function. PIRD are characterized by a broad range of clinical manifestations, including autoimmunity, infections, and lymphoproliferation (*3*, *4*). PIRD that impact cytokine signaling, including STAT3 gain-of-function (STAT3 GOF), allow us to focus on the impact of those pathways on human T cell function.

STAT3 GOF is a rare PIRD initially identified 10 years ago in patients with early-onset severe multi-organ autoimmunity; there are 191 individuals and 72 genetic variants described in the most recently published global cohort (*5–9*). STAT3 GOF is caused by a range of variants across the *STAT3* gene that affect all protein domains, with variable functional impacts on STAT3 function. These include altered magnitude of phosphorylation, delayed dephosphorylation, and/or amplified baseline STAT3 signaling (*8–10*). STAT3 GOF manifests with a heterogeneous and broad spectrum of clinical features, but most patients experience early onset autoimmunity (e.g., cytopenias, enteropathy, skin disease, interstitial lung disease and arthritis), lymphoproliferation, growth failure, and increased susceptibility to infections, including viral infections (*8*, *9*, *11*). Broadly, JAK/STAT signaling impacts the transcriptional regulation of immune cell differentiation and activation by transmitting extracellular stimuli to the cell nucleus (*12–14*). STAT3 is a transcription factor activated downstream of numerous cytokines, including the gp130 family of cytokines (e.g., IL-6), IL-10 family of cytokines, IL-12 family of cytokines (e.g., IL-27), adipokines (e.g., leptin), and growth factors (*15*, *16*). STAT3 signaling regulates diverse immune cell processes, including proliferation, differentiation, apoptosis, and inflammation (*15*).

Given the early onset autoimmunity experienced by patients with STAT3 GOF and the impact of STAT3 on regulation of T_H_17 and regulatory T cell (T_REG_) differentiation (*17–19*), initial investigations into STAT3 GOF mechanisms focused on the CD4^+^ T cell compartment (*20–25*). While some patients show evidence of decreased T_REG_ and increased T_H_17 frequencies (*9*, *23*, *26*, *27*), this has been variable. Recent work in two different mouse models of STAT3 GOF (C57BL/6 STAT3^+/G421R^ and STAT3^+/K392R^) have shown normal T_REG_ development, including differentiation and phenotype, and suppressive function. Overall, this suggests that T_REGS_ may not be the main drivers of disease pathology (*24*, *25*). However, consistent with the key role for STAT3 in CD8^+^ T cell differentiation (*28*, *29*), patients with STAT3 GOF have been shown to have altered CD8^+^ T cell differentiation and mouse models of STAT3 GOF have suggested a role for CD8^+^ T cells as drivers of autoimmunity (*30*, *31*). As an example, STAT3 GOF has been identified as a monogenic cause of neonatal onset type 1 diabetes (T1D) in some STAT3 GOF patients, including patients with the p.K392R variant (*5*). STAT3 knock-in mice bearing the p.K392R variant have been generated on both a C57BL/6 and on nonobese diabetic (NOD) backgrounds (*24*, *30*). In the NOD model, STAT3^+/K392R^ mice demonstrated more rapid onset and higher prevalence of T1D (*30*). CD8^+^ T cells in the pancreatic islets in this model demonstrated altered differentiation and enhanced cytotoxic gene signatures by single cell RNA-sequencing (*30*). Moreover, restricting expression of the STAT3 GOF variant to islet-specific TCR transgenic CD8^+^ T cells followed by adoptive transfer into NOD SCID recipients was sufficient to increase the prevalence of T1D compared to those recipients receiving WT STAT3 islet-specific TCR transgenic CD8^+^ T cells (*30*). In a separate study, pathogenic effector CD8^+^ T cells contributed to autoimmunity and mortality in homozygous mice with the STAT3 p.K658N variant (*31*). Moreover, depletion of CD8^+^ T cells in that model diminished both autoimmune and inflammatory pathology, as well as mortality (*31*). Mouse models of STAT3 GOF thus highlight dysregulation of effector CD8^+^ T cell differentiation and function (*24*, *25*, *30*, *31*).

Given the evidence of altered CD8^+^ T cell differentiation in STAT3 GOF, a chronic inflammatory monogenic disorder (*9*, *32*), we considered the potential role of dysfunctional immunoregulatory mechanisms. In addition to the fundamental nature of antigen, costimulatory, and cytokine signals, CD8^+^ T cell activation and function are also modulated by regulatory mechanisms, including purinergic signaling (*33–35*). While adenosine 5’-triphosphate (ATP) is a major source of energy in cells, once released from damaged or activated cells, extracellular ATP (eATP), acts as a danger-associated molecular pattern (DAMP) by binding P2 purinergic receptors (e.g., P2X7R) to trigger signaling cascades that lead to immune cell activation and the release of pro-inflammatory cytokines (*36*). To prevent eATP-induced pathology, ectoenzyme CD39 (*ENTPD1*) hydrolyzes eATP into ADP and ADP into AMP (*37*, *38*). Subsequently, CD73 (*NT5E*) breaks down AMP into immunosuppressive adenosine. This inhibits signaling cascades and can dampen overactive CD8^+^ T cell effector function, including cytokine production (e.g., IFN-γ), cytotoxicity, and proliferation, in part via binding to adenosine receptors (e.g., A_2A_R) (*39–41*).

Here, we investigated whether purinergic immunoregulatory signaling was intact in STAT3 GOF and how this impacted CD8 T^+^ cell differentiation and function. We found that human and mouse STAT3 GOF CD8^+^ T cells show evidence of dysregulation in their immune phenotype and transcriptional profiles, including aberrant pro-inflammatory cytokine production (IFN-γ and TNF-α). We also observed evidence of dysregulated purinergic signaling in CD8^+^ T cells from patients with STAT3 GOF. Specifically, while there was increased CD39 on CD8^+^ T cells from patients with STAT3 GOF, downstream expression of both CD73 and the adenosine receptor, A_2A_R, were decreased, suggesting that this immunoregulatory pathway is impaired. We demonstrate that STAT3 signaling in patients with STAT3 GOF and mouse models of STAT3 GOF, as well as in healthy control (HC) T cells cultured with STAT3-activating cytokines, regulates purinergic pathway ectoenzyme expression on both human and mouse CD8^+^ T cells. To test the impact of inhibiting amplified STAT3 signaling on CD8^+^ T cell function, we evaluated patients with STAT3 GOF who were receiving JAK inhibition (JAKi) as targeted therapy. This yielded correction of amplified inflammatory cytokine production by CD8^+^ T cells, as well as partial normalization of CD73 and normalization of A_2A_R expression. Together, these data suggest that dysregulation of the purinergic axis contributes to CD8^+^ T cell dysfunction in STAT3 GOF. This axis may suggest novel future strategies for targeted therapy in this rare disease, and perhaps in more common inflammatory disorders as well.

## RESULTS

### The CD8^+^ T cell compartment is dysregulated in patients with STAT3 gain-of-function

To better define the effect of amplified STAT3 signaling *in vivo*, we evaluated phenotypic dysregulation of immune cells in patients with STAT3 GOF (**Fig. 1A, Table S1**). Peripheral blood mononuclear cells (PBMCs) from a cohort of 8 patients with STAT3 GOF at baseline (untreated) and 21 age-matched HC were profiled using 43-parameter mass cytometry (**Supp. Fig. 1A**). Within our cohort and consistent with prior publications, the proportion of bulk T cells, B cells, monocytes and dendritic cells did not significantly differ between patients with STAT3 GOF and HC (**Fig. 1B**) (*25*, *42–45*). However, given that STAT3 is a critical regulator of CD8^+^ T cell differentiation and activation in cancer and viral infections (*28*), and the evidence of altered CD8^+^ T cell phenotype in mouse models of STAT3 GOF, we defined the immune phenotype of CD8^+^ T cells from patients with STAT3 GOF (*24*, *25*, *28*, *30*, *31*). Within the conventional αβ T cell compartment, we found an increased frequency of CD8^+^ T cells in patients with STAT3 GOF compared to HC, leading to a decreased CD4:CD8 ratio (**Fig. 1C**). Moreover, we detected a significant decrease in naïve CD8^+^ T cells (CD45RA^+^CD27^+^) and an increased frequency of effector memory (T_EM_, CD45RA^-^CD27^+^) and effector memory cells re-expressing CD45RA (T_EMRA_, CD45RA^+^CD27**^-^**) CD8^+^ T cells (**Fig. 1D**). Beyond altered differentiation, we also assessed the activation state of CD8^+^ T cells and found downregulation of memory markers, including TCF- 1 and IL-7Rα (CD127), and an increased frequency of expression of a subset of activation markers and effector molecules including Granzyme B and Ki-67, the latter consistent with a proliferative population of CD8^+^ T cells (**Fig. 1E)**. There were no differences in the frequency of HLA-DR^+^CD38^+^ CD8^+^ T cells, which can be elevated in hyperinflammatory disorders, and in the setting of recent antigen exposure (**Fig. 1E**). However, there was an increased frequency of cells co-expressing PD-1 and CD39, which can be seen in highly activated and/or dysfunctional CD8^+^ T cells (**Fig. 1E**) (*46–50*). Of note, we did not detect any differences in the frequency of innate-like T cells, including non-conventional NKT cells (CD3^+^CD56^+^), gamma-delta T cells (CD3^+^TCRψ8^+^), and MAIT (CD3^+^CD26^+^CD161^+^) cells between patients and HC (**Supp. Fig. 1B**). However, we confirmed decreases in NK cells (**Fig. 1B**), with trends towards decreased frequencies of CD127^Low^FoxP3^+^ regulatory T cells (T_REG_) (**Supp. Fig. 1C**), and CD27^+^ memory B cells in patients with STAT3 GOF (**Supp. Fig. 1D**) (*25*, *42–45*). We also found an increased relative proportion of early (CD56^Bright^CD16^-^) NK cells (**Supp. Fig. 1E**), decreased non-classical (HLA-DR^+^CD16^+^CD14^-^) monocytes, and an increase in classical (HLA-DR^+^CD16^-^CD14^+^) monocytes, consistent with previous findings (**Supp. Fig. 1F**) (*51*). Given the altered differentiation and dysregulated activation state of CD8^+^ T cells in patients with STAT3 GOF, we next sought to define their transcriptional state.

**Figure 1:**
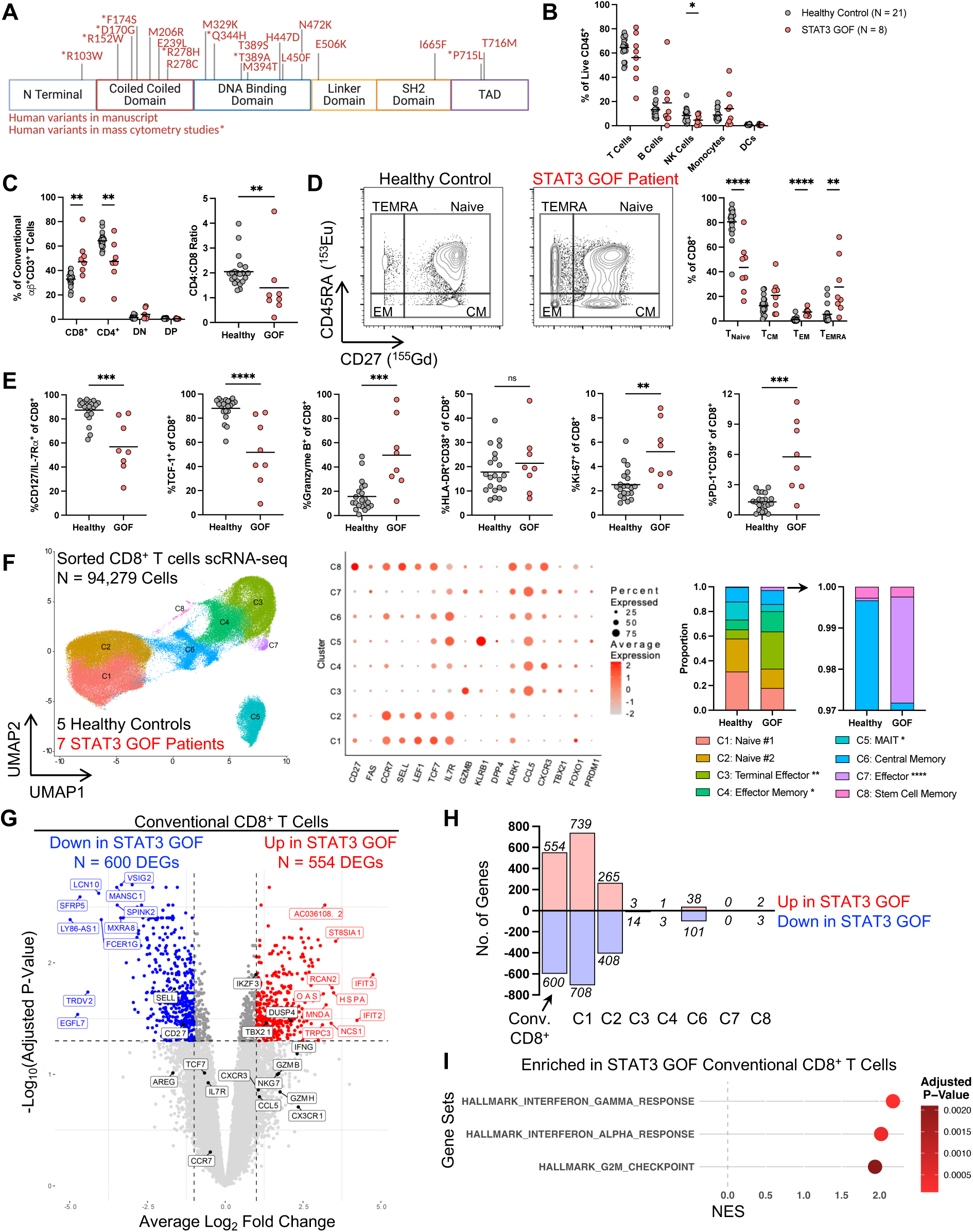
Patients with STAT3 GOF demonstrate a hyperactivated CD8^+^ T cell phenotype and transcriptional profile. **(A)** STAT3 protein domains with GOF variants in this manuscript indicated. **(B)** Frequencies of immune cell lineages in 8 untreated patients with STAT3 GOF and 21 age-matched HC. **(C)** Frequencies of CD3^+^αβ^+^ T cell subsets after exclusion of NKT (CD3^+^CD56^+^), Gamma-Delta (CD3^+^TCRψ8^+^), and MAIT (CD3^+^CD26^+^CD161^+^) cells; quantification of CD4:CD8 ratio. **(D)** Differentiation status of CD8^+^ T cells. Naïve = CD45RA^+^CD27^+^, Central Memory (CM): CD45RA^-^CD27^+^, Effector Memory (EM): CD45RA^-^ CD27^-^, Effector Memory Re-Expressing CD45RA (EMRA): CD45RA^+^CD27^-^. **(E)** Activation and effector molecule expression on CD8^+^ T cells. **(F)** scRNA-seq UMAP dimensionality reduction, cluster marker expression, and cluster proportion/annotation of sorted CD8^+^ T cells from 7 patients with STAT3 GOF and 5 age-matched HC. **(G)** Volcano plot demonstrating top 10 most upregulated (red, labeled), downregulated (blue, labeled) and selected genes (black, labeled) in conventional STAT3 GOF CD8^+^ T cells. **(H)** Number of differentially expressed genes by cluster. **(I)** GSEA of pathways enriched in conventional CD8^+^ T cells. Data are pooled from 3 independent experiments. For **B-E**: \**P* ≤ 0.05, \*\**P* ≤ 0.01, \*\*\**P* ≤ 0.001, \*\*\*\**P* ≤ 0.0001 by Mann-Whitney Test. For **F**: \**P* ≤ 0.05, \*\**P* ≤ 0.01, \*\*\*\**P* ≤ 0.0001 by *propeller* (moderated t-test). Not listed or ns was not statistically significant.

### STAT3 GOF CD8^+^ T cells exhibit the transcriptional imprint of inflammatory cytokine signaling

To better understand the impact of amplified STAT3 signaling on CD8^+^ T cells at a transcriptional level, we performed single-cell RNA sequencing (scRNA-seq) on sorted CD8^+^ T cells from 7 patients with STAT3 GOF (6 not receiving any immunomodulatory therapy and one receiving JAKi) and 5 age-matched HC. After quality control, 94,279 cells were further analyzed (*52–54*). After performing integration and clustering using Seurat and clustree packages, we identified 8 clusters of CD8^+^ T cells (**Fig. 1F)** (*55*). Final cluster annotations were defined using SingleR (*56*) with the Monaco Immune Cell Data reference dataset (*57*), and then assessed manually using marker expression (**Fig. 1F**) (*56*, *57*). This strategy demonstrated trends towards reduction in both naïve CD8^+^ T cell clusters (i.e., clusters 1 and 2, p-values = 0.07 and 0.12 respectively) in patients with STAT3 GOF (*58*). Naïve clusters were defined based on increased expression of *CCR7*, *SELL*, *LEF1*, *TCF7*, *IL7R* and absence of expression of effector and activation genes including *CCL5*, *CXCR3,* and *TBX21* (**Fig. 1F**). There were significant expansions of clusters of terminal effector (cluster 3: defined based on increased *GZMB, KLRB1, CCL5, TBX21, PRDM1* and absent expression of genes present in clusters 1 and 2; p-value = 0.0015), effector memory (cluster 4: increased expression of *CCR7*, *TCF7*, *IL7R*, *GZMB, CCL5, CXCR3, TBX21;* p-value = 0.019), and effector (cluster 7: increased expression of *FAS*, *GZMB*, *KLRB1*, *KLRK1*, *CCL5*, *CXCR3*, *TBX21*, *PRDM1*; p-value = 3.7 ×10^-6^) CD8^+^ T cells in STAT3 GOF (**Fig. 1F**). The remaining clusters were annotated as central memory (cluster 6: increased expression of *CD27*, *CCR7*, *TCF7*, *IL7R*, *CCL5,* and decreased expression of *TBX21*, *CXCR3* and *PRDM1*) and stem cell memory (cluster 8: presence of cluster 1 and 2 genes along with increased expression of *FAS, CXCR3* and effector genes *KLRK1*, *CCL5*, *TBX21,* and *FOXO1*) and did not differ between HC and STAT3 GOF. The last cluster, MAIT cells (cluster 5: *KLRB1*, *IL7R* and *DPP4*) was decreased in STAT3 GOF (p-value = 0.013) (**Fig. 1F**). Using a multi-modal approach with this large cohort we defined altered CD8^+^ T cell differentiation in STAT3 GOF, with an increase in the proportions of terminal effector, effector memory and effector T cells.

Defining the mechanism by which STAT3 GOF impacts CD8^+^ T cell differentiation and activation state required further evaluation of transcriptional networks within these T cell states, so we next evaluated differential gene expression (DGE) and performed gene set enrichment analysis (GSEA) (*59*). Differential gene expression within conventional CD8^+^ T cells (e.g., excluding cluster 5, MAIT cells) revealed increased expression of 554 genes and decreased expression of 600 genes. (**Fig. 1G-H, Table S2**). Genes upregulated in STAT3 GOF CD8^+^ T cells included genes associated with antiviral immunity (e.g., *IFIT2*, *IFIT3*, *OASL*) and genes associated with cytotoxic and effector function (*TBX21*, *CCL5, CXCR3,* and *NKG7*), while downregulated genes included *CD27, SELL*, *TCF7, IL7R* (**Fig. 1G**). To better understand how these gene expression changes altered the biological state and function of CD8^+^ T cells, we performed GSEA using the Hallmark Human MSigDB collection (*59*, *60*). This analysis demonstrated significant enrichment of three biological processes in patients with STAT3 GOF: HALLMARK_INTERFERON_GAMMA_RESPONSE (NES = 2.17, FWER p-value < 0.0001), HALLMARK_INTERFERON_ALPHA_RESPONSE (NES = 2.02, FWER p-value < 0.0001) and HALLMARK_G2M_CHECKPOINT (NES = 1.95, FWER p-value = 0.001), providing transcriptional evidence of interferon exposure and impact on cell cycle progression (**Fig. 1I, Supp. Fig. 2A**-C**, Table S3**). To connect transcriptional alterations to the environment that immune cells experience, we performed plasma proteomic analysis of patients with STAT3 GOF compared to HC (*61*). This analysis demonstrated synergistic evidence of systemic inflammation, including increased IL-1β, IL-6, IL-10, IL-18, IFN-γ, CXCL9, and CXCL10 (with the last two being downstream targets of IFN-γ) in STAT3 GOF (**Supp. Fig. 3A- B**) (*62–64*). Overall, patients with STAT3 GOF experience increased systemic inflammatory cytokine exposure and amplified STAT3 signaling, with CD8^+^ T cells demonstrating a greater transcriptional imprint of inflammatory cytokine exposure. Given the impacts of altered STAT3 on CD8^+^ T cell differentiation, and early onset autoimmunity in patients, we next quantified the impact of STAT3 GOF on effector function.

### STAT3 GOF CD8^+^ T cells demonstrate aberrant cytokine production profiles

Having observed alterations both in the immune phenotype and transcriptional profile, we next investigated the impact of STAT3 GOF on CD8^+^ T cell function, particularly on cytokine production. Compared to HC, there was no difference in IL-2 production by total or naïve (CD45RA^+^CD27^+^) CD8^+^ T cells from patients with STAT3 GOF (**Fig. 2A, Supp. Fig. 4A**). However, there was a trend towards a deficit in production of IL-2 by non-naïve (non- CD45RA^+^CD27^+^) CD8^+^ T cells (p-value = 0.063; **Fig. 2A**). In contrast, TNF-α and IFN-γ production were increased from total CD8^+^ T cells in patients with STAT3 GOF (**Fig. 2B-C**). Unlike IL-2, CD8^+^ T cell subset analysis revealed that naïve (CD45RA^+^CD27^+^) CD8^+^ T cells produced increased TNF-α and IFN-γ when stimulated (**Fig. 2B-C**, middle panels). However, non-naïve STAT3 GOF CD8^+^ T cells produced levels of TNF-α and IFN-γ in response to stimulation that were comparable to HC (**Fig. 2B-C**, right panels). Given that early onset autoimmunity is frequently found in patients with STAT3 GOF, with evidence of pathogenic CD8^+^ T cells contributing to lymphoproliferation and autoimmunity in mouse models of STAT3 GOF (*24*, *30*, *31*), we next sought to define potential underlying mechanisms of dysregulated CD8^+^ T cell function in STAT3 GOF.

**Figure 2:**
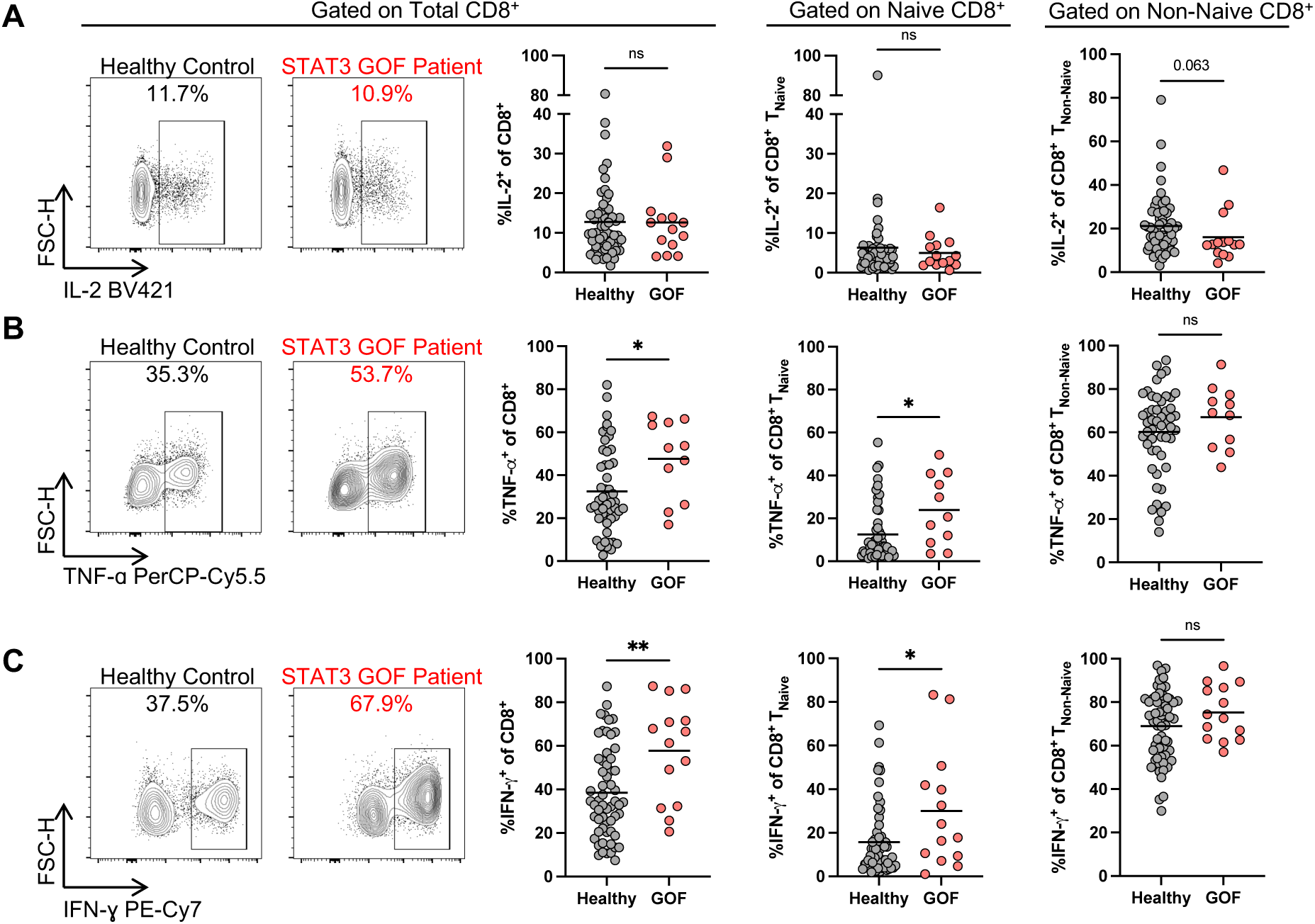
STAT3 GOF differentially affects effector function in a subset-specific manner in CD8^+^ T cells. Quantification of **(A)** IL-2, **(B)** TNF-α, and **(C)** IFN-γ cytokine production within total CD8^+^, naïve (CD45RA^+^CD27^+^) and non-naive (antigen-experienced or non- CD45RA^+^CD27^+^) CD8^+^ T cells following 4hr stimulation with PMA/Ionomycin in the presence of BFA/Monensin. Data are pooled from 7 to 8 independent experiments with n = 51 to 59 HC and n =11 to14 untreated patients with STAT3 GOF. \**P* ≤ 0.05, \*\**P* ≤ 0.01 by Mann-Whitney Test.

### Increased expression of CD39 on STAT3 GOF CD8^+^ T cells

We began by considering whether chronic inflammation may be impairing immune regulatory pathways and thus contributing to dysregulated CD8^+^ T cell function. While there are multiple regulatory systems and networks in the immune system which could be engaged to counteract chronic inflammation in STAT3 GOF, we noted that compared to HC, CD8^+^ T cells from patients with STAT3 GOF displayed elevated levels of the ectoenzyme, CD39 (**Fig. 3A**). Expression of CD39 was also increased on other immune cells including CD4^+^ T cells (including T_REGS_) and NK cells (**Supp. Fig. 5A**-D). In populations of immune cells constitutively expressing CD39, including B cells, monocytes, and dendritic cells, expression was comparable between STAT3 GOF and HC (**Supp. Fig. 5E**-G). Functionally, CD39, in conjunction with CD73, promotes immune suppression through the hydrolysis of extracellular ATP (eATP) into the immunosuppressive molecule, adenosine (**Fig. 3B**). In addition, the purinergic axis has been shown to promote forms of CD8^+^ T cell dysfunction including T cell exhaustion (T_EX_), a state of impaired function found alongside increased expression of inhibitory receptors and altered transcriptional networks and epigenetic poise, in the setting of chronic exposure to antigen and inflammation (e.g., cancer and chronic viral infection) (*47*, *50*, *65–70*). In that context, CD39 has been used as a marker to identify the most terminally differentiated exhausted CD8^+^ T cells (*46*). To evaluate whether T_EX_ is present in STAT3 GOF, we assessed TOX expression, a key transcriptional and epigenetic driver of T_EX_, in patients with STAT3 GOF compared to HC, using spectral flow cytometry. We recapitulated the differences in CD8^+^ T cell differentiation (**Supp. Fig. 6A**), and within these individuals, we found no significant differences in TOX protein expression in non-naïve CD8^+^ T cells from patients with STAT3 GOF vs. HC (**Supp. Fig. 6B**). When we evaluated the expression of a gene set with increased expression in T_EX_ (*71*) in our scRNA-seq data, we did not find a significant increase in expression in non-naïve CD8^+^ T cells from patients with STAT3 GOF (**Supp. Fig. 6C**). Overall, these data are not consistent with increase in T_EX_ in STAT3 GOF. With regards to other forms of CD8^+^ T cell dysregulation, previous evaluation of CD8^+^ T cells in STAT3 GOF has identified increased frequencies of CD57^+^ effector CD8^+^ T cells; these cells share similarities with the effector NKG2D^+^CD8^+^ T cells that contribute to autoimmune and inflammatory pathology in a mouse model of STAT3 GOF (*31*). In our cohort, we did not identify increased expression of NKG2D on CD8^+^ T cells (**Supp. Fig. 6D**) (*31*). However, we found an increased frequency of CD57^+^CD8^+^ T cells, but minimal co-expression of CD57 with CD39, suggesting that both CD57^+^ and CD39^+^ populations are present as non-overlapping CD8^+^ T cell states in STAT3 GOF **(Supp. Fig. 6E)**. Therefore, we sought to better understand the mechanisms underlying increased CD39 expression in CD8^+^ T cells in patients with STAT3 GOF.

**Figure 3:**
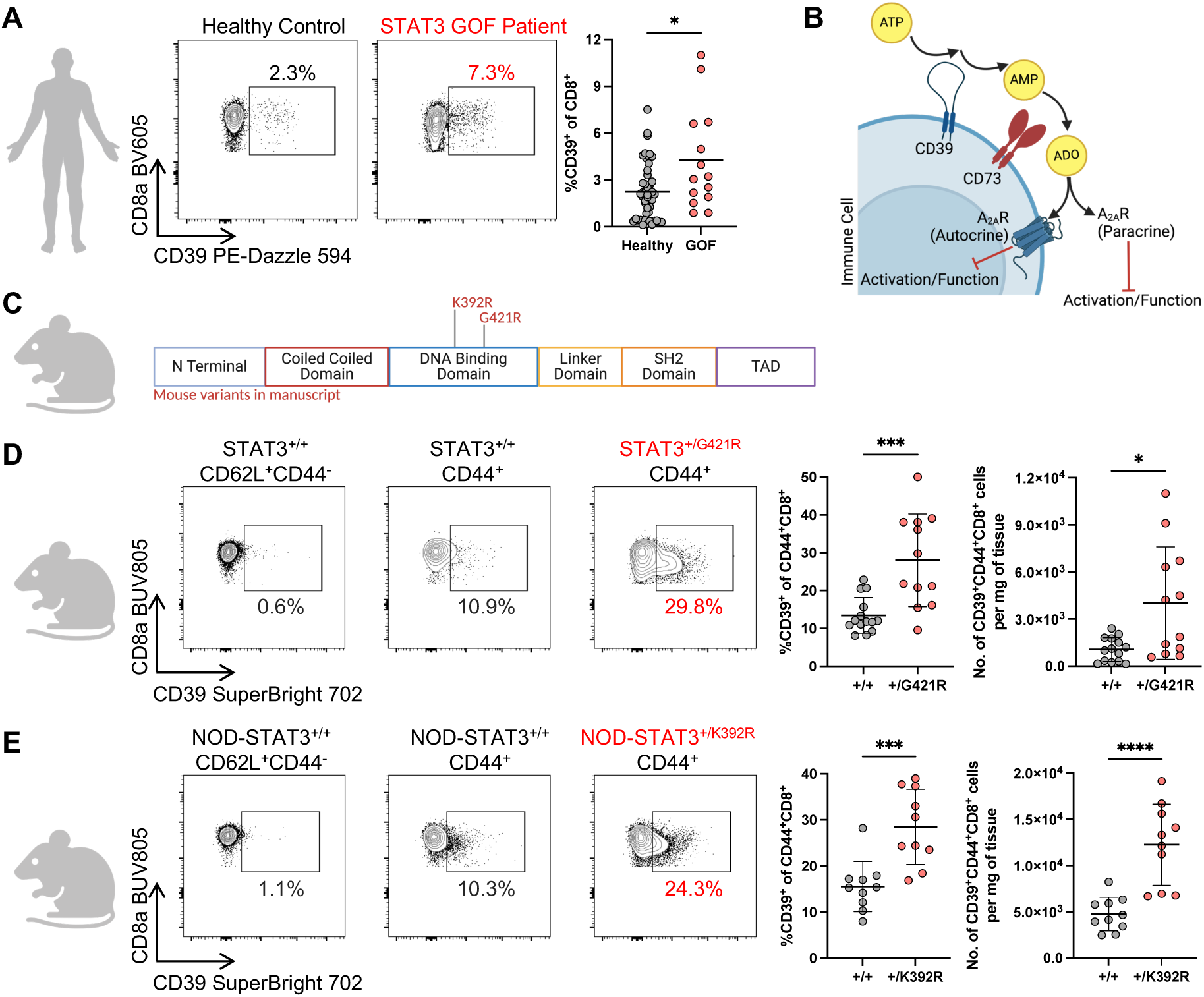
Increases in the regulatory ectoenzyme CD39 are conserved across humans and mouse models of STAT3 GOF. **(A)** Spectral flow cytometry gating and quantification of CD39 frequency within CD8^+^ T cells in HC (n = 47) and patients with STAT3 GOF (n = 14). **(B)** Schematic of purinergic signaling regulation of immune cell activation and function. **(C)** Schematic of STAT3 protein with human STAT3 GOF variants investigated in mouse models. **(D)** Representative flow cytometry plots of CD39 frequency on CD8^+^CD44^+^ cells, frequencies and absolute counts of CD39^+^CD44^+^CD8^+^ T cells in spleens from STAT3^+/+^ (n = 14) and STAT3^+/G421R^ (n = 12) mice. **(E)** Representative flow cytometry plots of CD39 frequency on CD8^+^CD44^+^ cells, frequencies and absolute counts of CD39^+^CD44^+^CD8^+^ T cells in spleens from NOD-STAT3^+/+^ and NOD-STAT3^+/K392R^ mice (n = 10 each). For **A**: data are pooled from 6 independent experiments; for **D**: data are pooled from 3 independent experiments; for **E**: data are pooled from 2 independent experiments. \**P* ≤ 0.05, \*\*\**P* ≤ 0.001, \*\*\*\**P* ≤ 0.0001 by Mann-Whitney Test.

Having identified an increased frequency of CD39^+^ CD8^+^ T cells in patients with STAT3 GOF, we sought to understand the role of STAT3 signaling in regulating purinergic signaling, starting with control of CD39 expression in humans and mice. There is evidence that the CD39 (*Entpd1)* locus is directly bound by STAT3 in mouse CD4^+^ T cells (*72*). We also found that CD39 (*Entpd1*) is directly bound by STAT3 in mouse CD4^+^ T cells, based on reanalysis of published chromatin immunoprecipitation-sequencing (ChIP-seq) data (**Supp. Fig. 7A)** (*73–75*). Specifically, these data show that IL-27-driven STAT3 engages multiple sites upstream of the *Entpd1* locus (**Supp. Fig. 7A**). These findings are consistent with work demonstrating that IL-6 mediated-STAT3 activation promotes CD39 expression in CD4^+^ T_H_17 cells (*72*), and is consistent with increased CD39 expression in CD4^+^ T cells (**Supp. Fig. 5B**), including T_REGS_ (**Supp. Fig. 5C**) in patients with STAT3 GOF. Prior data and our work thus support that *Entpd1* is a direct target of STAT3 in mouse CD4^+^ T cells, and while we have shown that CD8^+^ T cells from patients with STAT3 GOF have increased expression of CD39 (**Fig. 3A**), the impact of amplified STAT3 expression on CD39 expression on CD8^+^ T cells remained to be determined.

With evidence that *Entpd1* is a target of STAT3, we next tested whether the increased CD39 expression in STAT3 GOF patients was also evident in CD8^+^ T cells from STAT3 GOF mouse models (*25*, *30*, *76*). Consistent with prior work, we found that STAT3^+/G421R^ and NOD- STAT3^+/K392R^ mice (**Fig. 3C**) develop splenomegaly (**Supp. Fig. 8A, 9A**), and evidence of immune dysregulation as indicated by altered CD8^+^ T cell differentiation (**Supp. Fig. 8B**-C**, 9B-C, E**) (*25*, *30*). Detailed immunophenotyping of CD8^+^ T cells in both models also revealed altered expression of activation markers, including PD-1 and Ki-67 (**Supp. Fig. 8D, 9D**). Effector cytokines, including IFN-γ and TNF-α, were increased in both mouse models of STAT3 GOF, consistent with our evaluation of patients with STAT3 GOF (**Supp. Fig. 8E**-G**, 9F-H**). However, unlike patients with STAT3 GOF, in whom we observed increased naïve CD8^+^ T cell cytokine production, this increase was predominantly from CD8^+^CD44^+^ cells (**Supp. Fig. 8F, 9G**). Within CD44^+^CD8^+^ T cells, both STAT3^+/G421R^ and NOD-STAT3^+/K392R^ mice, respectively, showed approximately a 2- fold increased frequency of CD39^+^ cells and an increase in the absolute count of CD39^+^CD44^+^CD8^+^ T cells (**Fig. 3D-E**). Additionally, we re-analyzed published scRNA-seq data (*30*) evaluating CD45^+^ lymphocytes isolated from the pancreatic islets of 8-10 week-old NOD- STAT3^+/+^ and NOD-STAT3^+/K392R^ mice prior to diabetes onset (**Supp. Fig. 6A**). Analysis of identified CD8^+^ T cell clusters in the scRNA-seq data revealed a significant, though small amplitude, upregulation of *Entpd1* (CD39) expression in NOD-STAT3^+/K392R^ mice (Log_2_F.C. = 0.15, adjusted p-value = 2.7×10^-8^) (**Supp. Fig. 10A**), consistent with our flow cytometry (**Fig. 3E**). Therefore, in both C57BL/6 and NOD backgrounds, STAT3 GOF mouse models demonstrate an increase in CD39 expression on CD8^+^ T cells. Given that we observed consistent upregulation of CD39 on CD8^+^ T cells from patients with STAT3 GOF and in two mouse models of STAT3 GOF, we next evaluated the signals regulating CD39 expression on CD8^+^ T cells.

### STAT3 signaling, in the setting of TCR stimulation, directly induces CD39 expression on CD8^+^ T cells from HC

Since T cell activation is governed by a variety of signals, including T cell receptor (TCR) engagement, we next assessed the kinetics of CD39 upregulation upon αCD3/αCD28 activation in PBMCs from HC. At baseline (without activation), CD8^+^ T cells from HC express low levels of CD39, but CD39 expression is upregulated in a time-dependent manner over a 96-hour period (**Fig. 4A**). T cell activation is also regulated by the cytokine environment (*77*, *78*). To test the cell intrinsic impacts of amplified STAT3 on CD39 expression, we developed an acute *in vitro* model in which isolated CD8^+^ T cells from HC were left unstimulated or stimulated with αCD3/αCD28 alone, STAT3-activating cytokines alone, or a combination of αCD3/αCD28 plus cytokines for 96 hours (**Fig. 4B**). STAT3 is activated downstream of a variety of cytokines, including IL-6, which we and others have identified as being increased systemically in STAT3 GOF (**Supp. Fig. 3A)** (*32*, *79*). To test whether increasing STAT3 signaling induces higher levels of CD39, we evaluated additional cytokines which induce high levels of phospho-STAT3 (Tyrosine 705 residue, pSTAT3^Y705^) and, to varying degrees, phospho-STAT1 (Tyrosine 701 residue, pSTAT1^Y701^) including IL-10, IL-21, and IL-27 (**Supp. Fig. 11A**-B). *In vitro* culture of CD8^+^ T cells from HC with these cytokines alone was not sufficient to induce CD39 expression (**Supp. Fig. 11C)**. However, activation of T cells with αCD3/αCD28 alone led to robust induction of CD39, which was further augmented when STAT3-activating cytokines (e.g., IL-6, IL-10, IL-21, or IL-27) were added to the cultures (**Fig. 4B**). To further test the link between STAT3 signaling and upregulation of CD39, we evaluated the impact of these individual cytokine T cell cultures on levels of pSTAT3. First, we assessed levels of pSTAT3 within CD8^+^ T cells that did not upregulate CD39 (CD8^+^CD39^Low^) and those that did upregulate CD39 (CD8^+^CD39^High^). Irrespective of stimulation conditions (i.e., αCD3/αCD28 vs. αCD3/αCD28 + cytokine), pSTAT3 levels were higher in CD8^+^CD39^High^ cells relative to CD8^+^CD39^Low^ cells (**Fig. 4C**).

**Figure 4:**
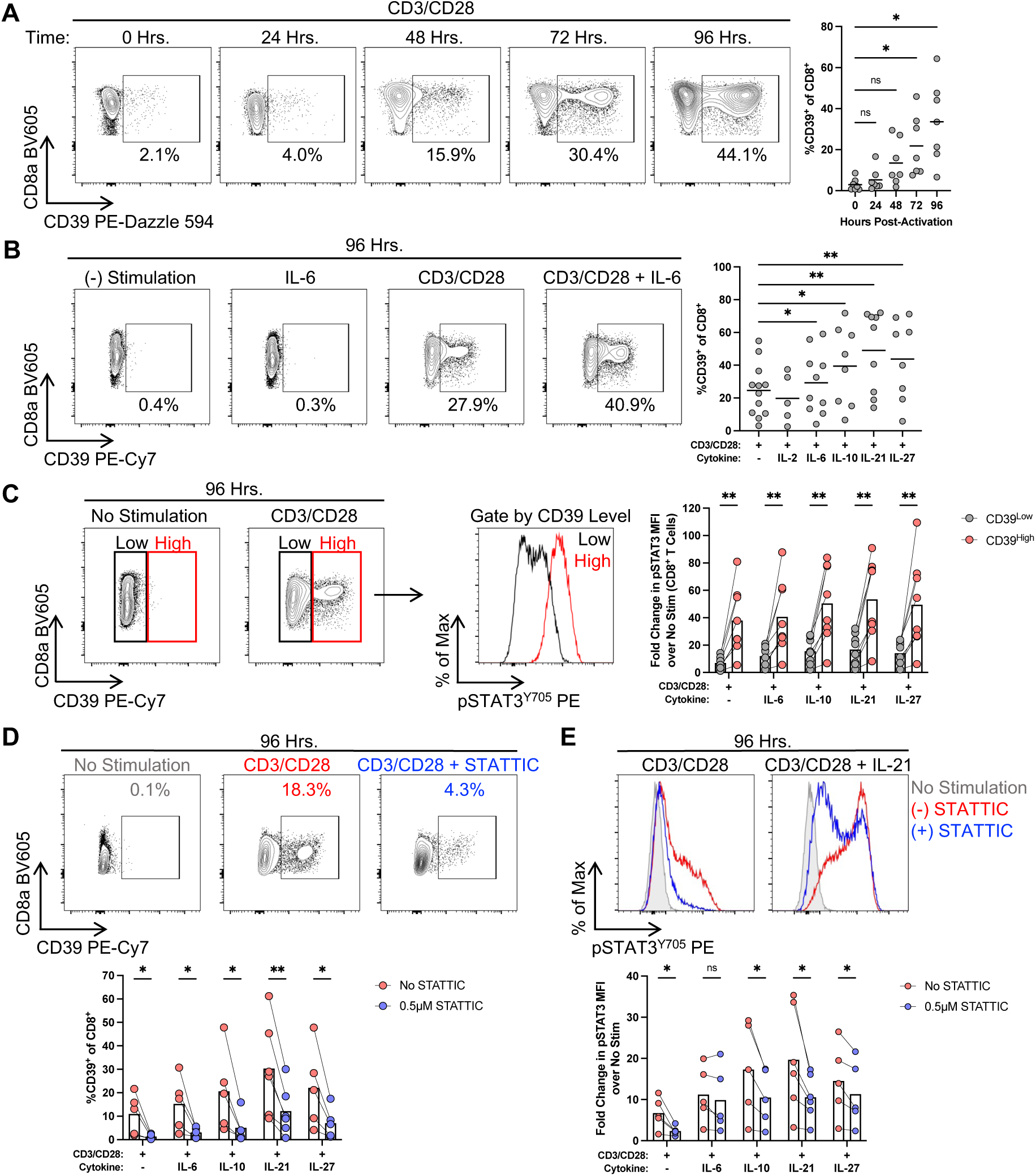
T cell receptor engagement and STAT3-activating cytokines synergize to increase CD39 expression in healthy donor CD8^+^ T cells. **(A)** Representative flow cytometry plots of CD39 induction in PBMCs (gated on CD8^+^ T cells) over a 96-hour period following αCD3/αCD28 activation, and quantification of CD39-positivity (n = 7). **(B)** Representative plots of CD39- positivity in isolated CD8^+^ T cells left unstimulated, stimulated with IL-6 alone or αCD3/αCD28 ± IL-6; quantification of CD39-positivity across various αCD3/αCD28 ± cytokine conditions (n > 5). **(C)** Stratification of CD39^Low^ and CD39^High^ CD8^+^ T cells and pSTAT3^Y705^ intensity; quantification of fold change in pSTAT3^Y705^ levels across CD39 levels in αCD3/αCD28 ± cytokine conditions (n = 8). **(D)** Representative plots and quantification of CD39^+^CD8^+^ T cells in the presence of the STAT3 inhibitor, STATTIC, after 96 hours of activation (n > 5). **(E)** Representative histograms of pSTAT3^Y705^ levels in HC CD8^+^ T cells in the presence of STATTIC (n > 5). For **A**: data are pooled from 2 independent experiments. For **B-E**: data are pooled from > 5 independent experiments; each experiment performed in a single donor with each data point representing the average of 2-3 technical replicates. \**P* ≤ 0.05 \*\**P* ≤ 0.01 by Multiple Paired Mann-Whitney Tests or RM one-way ANOVA with Dunnett’s multiple comparison test; adjusted p-values are indicated for multiple comparisons.

Having assessed differences in pSTAT3 levels in CD39 expressing and non-expressing cells, we tested whether this increase in CD39 expression is dependent on STAT3 signaling. To do this we used a STAT3 inhibitor, STATTIC (*80*), to inhibit phosphorylation of STAT3 in our *in vitro* CD8^+^ T cell cultures. This drug primarily inhibits STAT3, though at high doses it can also impair STAT1 (**Supp. Fig. 11D**-E). Therefore, we identified a dose of STATTIC (0.5µM) that reduced pSTAT3 with minimal impact on pSTAT1 (**Supp. Fig. 11D**-E). In αCD3/αCD28-activated culture conditions in the presence or absence of STAT3-activating cytokines, incubation with STATTIC decreased the frequency of CD39^+^CD8^+^ T cells (**Fig. 4D**) and decreased pSTAT3 levels (**Fig. 4E**). These data suggest that STAT3 directly contributed to the regulation of CD39 expression in a TCR-dependent manner in HC CD8^+^ T cells.

### Activation induces greater CD39 expression in STAT3 GOF CD8^+^ T cells

Having established that STAT3 regulates CD39 expression *in vitro*, we next assessed CD39 upregulation of CD8^+^ T cells from patients with STAT3 GOF upon αCD3/αCD28 activation in the absence of exogenous cytokines. After 4 days of *in vitro* activation with αCD3/αCD28, cells from patients with STAT3 GOF had an increased frequency of CD39^+^CD8^+^ T cells and a 2.1-fold increase in relative expression of CD39 in STAT3 GOF compared to age-matched HCs (relative fold change in CD39 MFI: HC = 7.8, STAT3 GOF = 16.1) (**Fig. 5A**). To further evaluate the connection between the amplitude of STAT3 signaling and the level of CD39 expression, we compared our patients with STAT3 GOF and HC to a cohort of patients with STAT3 dominant negative (DN) Hyper IgE syndrome. STAT3 DN (formerly known as Job’s syndrome), is a monogenic immune deficiency caused by heterozygous STAT3 variants, which yields decreased STAT3 signaling (*44*, *81–84*). We found that, at baseline, CD39 expression in CD8^+^ T cells from patients with STAT3 DN was comparable to HC and reduced when compared to STAT3 GOF patients (**Supp. Fig. 12A**). However, upon activation, CD8^+^ T cells from patients with STAT3 DN had a decreased extent of CD39 induction compared to HC (**Supp. Fig. 12B**). By comparing the impact of amplifying and diminishing STAT3 signaling, in STAT3 GOF and STAT3 DN respectively, we find evidence to support the hypothesis that modulating the amplitude of STAT3 signaling regulates CD39 expression *ex vivo* upon activation of CD8^+^ T cells.

**Figure 5:**
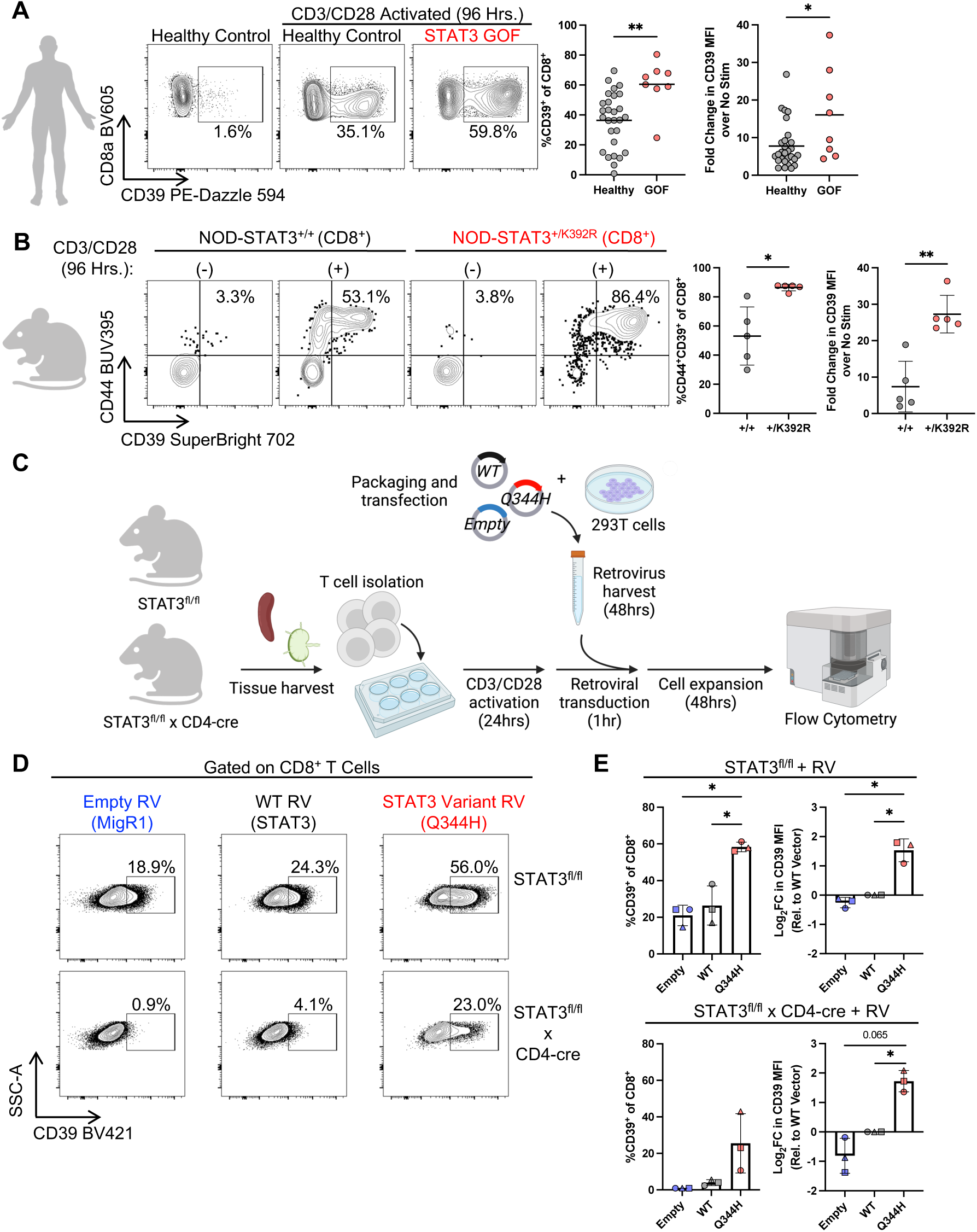
CD8^+^ T cells bearing a STAT3 GOF variant induce higher levels of CD39 upon T cell receptor engagement in humans and mice. **(A)** Analysis of CD39 induction (frequency and relative MFI) in HC (n = 28) and untreated patients with STAT3 GOF (n = 8) after 4 days of activation with αCD3/αCD28. **(B)** CD39 induction in isolated naive (CD62L^+^CD44^-^) CD8^+^ T cells from NOD-STAT3^+/+^ and NOD-STAT3^+/K392R^ spleens (n = 5 each) after 96 hours of activation with αCD3/αCD28. **(C)** Introduction of STAT3 GOF variants into STAT3^fl/fl^ (control) or STAT3^fl/fl^ x CD4-cre (test) murine T cells using a retroviral transduction system with Empty (MigR1), WT STAT3, or a STAT3 GOF variant (Q344H). **(D)** Representative CD39 flow cytometry plot in CD8^+^T cells in control and test mice across retroviral transduction conditions. **(E)** Quantification (frequency and relative expression) of CD39^+^CD8^+^ T cells from STAT3^fl/fl^ (top) or STAT3^fl/fl^ x CD4-cre (bottom) mice (n = 3). For **A**: data are pooled from 3 independent experiments; for **E**: data are pooled from 3 independent experiments as indicated by symbol shape. Where asterisks are used: \**P* ≤ 0.05, \*\**P* ≤ 0.01 by Mann-Whitney Test or Kruskal-Wallis test with Dunn’s multiple comparison test; for comparisons of more than 2 groups, adjusted p-values are indicated.

To test whether higher CD39 induction in STAT3 GOF was due to amplified STAT3 signaling *in vivo*, we returned to a mouse model of STAT3 GOF. First, we isolated naïve (CD62L^+^CD44^-^) CD8^+^T cells from the spleens of NOD-STAT3^+/K392R^ mice and tested their ability to upregulate CD39. Naïve CD8^+^ T cells isolated from the spleens of NOD-STAT3^+/K392R^ mice and activated with αCD3/αCD28 generated higher frequencies of CD39^+^CD8^+^ T cells compared to NOD-STAT3^+/+^ littermate controls (**Fig. 5B**). Second, we used a system in which STAT3 signaling was endogenously absent in mouse T cells to directly compare the impact of STAT3 GOF variants on CD39 levels. *Stat3* flox/flox (STAT3^fl/fl^) mice were crossed to CD4-Cre mice to yield STAT3^fl/fl^ x CD4-cre which specifically ablates STAT3 expression in all T cells (*85*). To assess the ability of STAT3 mutants to mobilize target genes, we transduced T cells from WT (*Stat3*^fl/fl^) or STAT3 deficient (*Stat3*^fl/fl^ x CD4-cre) mice with either WT or STAT3 GOF patient variants and measured CD39 by flow cytometry. Importantly T cells from WT STAT3 mice approximate the scenario in humans, where WT and mutant alleles co-exist, while STAT3 deficient T cells present a ‘clean’ background where the only available STAT3 is the one that we introduce. These studies showed that in control STAT3^fl/fl^ mice, containing endogenous WT STAT3, introduction of the p.Q344H variant was sufficient to drive higher levels of CD39 compared to introducing a WT STAT3 (**Fig. 5D-E**, top row). In T cells from STAT3-deficient mice, we found that WT STAT3 induces higher levels of CD39, and the STAT3 GOF variant (p.Q344H) drove even higher levels of CD39 than WT STAT3 (**Fig. 5D-E**, bottom row). Together, these data suggest that activating variants that increase STAT3 signaling augment CD39 expression upon TCR engagement without the addition of exogenous cytokines. This leaves us with a paradoxical finding in STAT3 GOF of increased frequencies of CD39^+^CD8^+^ T cells, which could be anticipated to constrain effector function, in parallel with CD8^+^ T cells that have been shown to have an amplified potential to secrete inflammatory cytokines. This prompted further interrogation of the purinergic signaling axis in in CD8^+^ T cells in STAT3 GOF.

### STAT3 GOF patients have alterations in key purinergic pathway members in CD8^+^ T cells

Downstream of CD39, CD73 (*NT5E)* and A_2A_R (*ADORA2A*) are key partners involved in T cell purinergic signaling. CD73 is an ectoenzyme that transforms AMP to adenosine, while A_2A_R is a key adenosine receptor on T cells. scRNA-seq from sorted CD8^+^ T cells revealed that expression of *NT5E* and *ADORA2A* were both decreased on STAT3 GOF total CD8^+^ T cells compared to HC (**Fig. 6A**). We hypothesized this could prevent CD8^+^ T cells from producing and sensing adenosine from ATP, even if increased AMP was produced in the setting of increased expression of CD39. We therefore evaluated protein levels of CD73 and A_2A_R across major immune cell subsets, including CD8^+^ T cells in patients with STAT3 GOF and HC (**Fig. 6B-C, Supp. Fig. 13A**-B). We confirmed that CD73 protein levels were decreased on total CD8^+^ T cells from patients with STAT3 GOF (**Fig. 6B**), and in both naive and non-naïve CD8^+^ T cells separately (**Supp. Fig. 14A**). Patients with severe STAT3 GOF are often treated with immunomodulatory therapy that directly inhibits the upstream components of the signaling pathway (e.g., JAKi ± α-IL-6R, tocilizumab) (*86*). Therefore we could evaluate the impact of *in vivo* modulation of STAT signaling with samples from patients with STAT3 GOF treated with JAKi ± α-IL6R (*9*, *86*). Here, we found that CD73 levels on total CD8^+^ T cells from patients with STAT3 GOF who were receiving JAKi were comparable to HC (**Fig. 6B**, **Supp. Fig. 14A**). Further along the purinergic signaling pathway, A_2A_R expression was also decreased on CD8^+^ T cells in untreated patients with STAT3 GOF (**Fig. 6C**). This observed decrease in A_2A_R was normalized on CD8^+^ T cells from patients with STAT3 GOF treated with JAKi (**Fig. 6C**), and in both naive and non-naïve CD8^+^ T cell subsets (**Supp. Fig. 14B**). Interestingly, we found that CD39 levels did not decrease with JAKi (**Supp. Fig. 14C**). Thus, at baseline patients with STAT3 GOF demonstrate increased potential for naïve CD8^+^ T cells to secrete inflammatory cytokines (**Fig. 2B-C**) along with reduced levels of the terminal components of the immunoregulatory purinergic signaling cascade (**Fig. 6B-C**), the latter of which was found to be partially corrected with JAKi ± α-IL6R.

**Figure 6:**
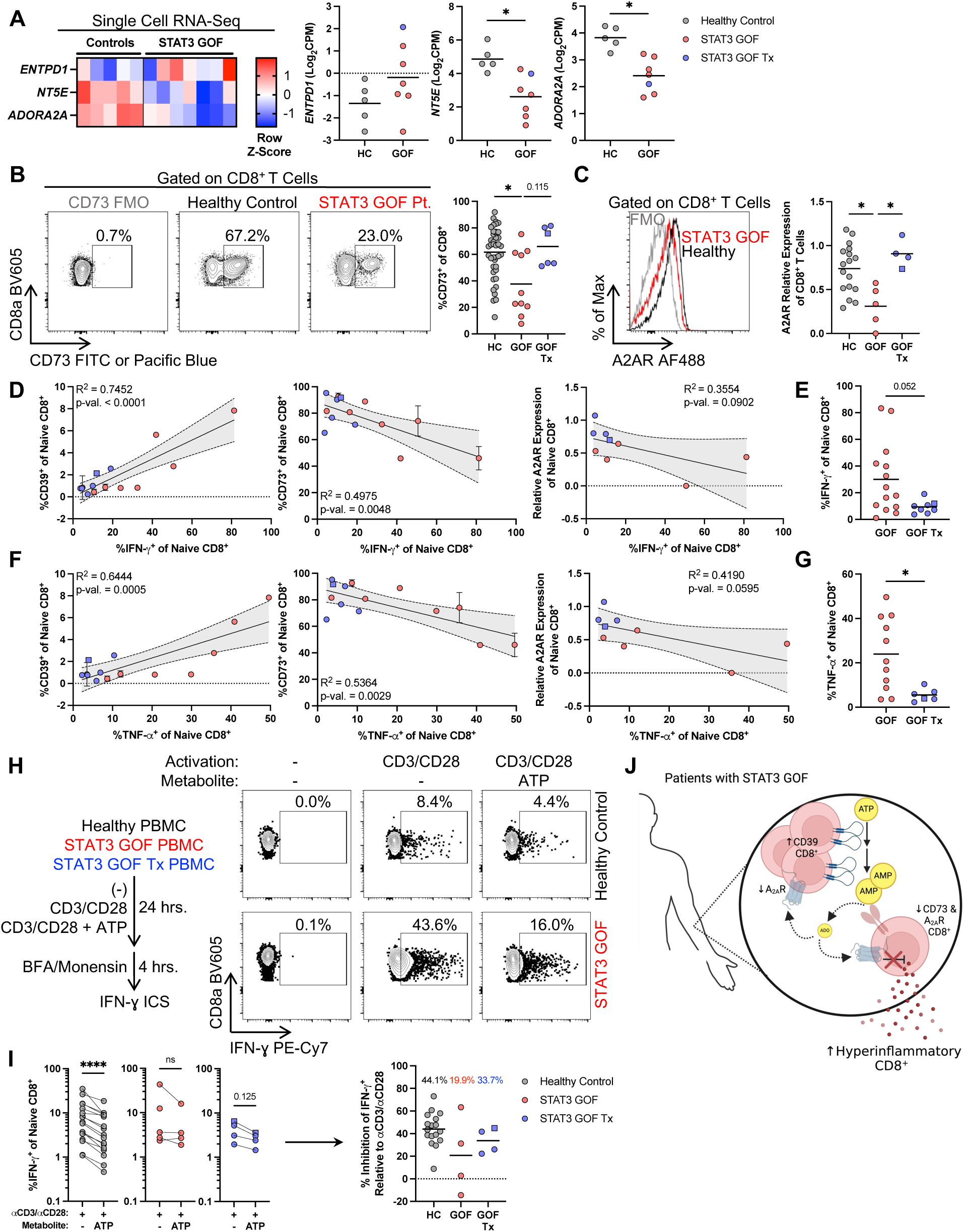
Altered purinergic signaling molecules which correlate with aberrant functionality, with some normalization with targeted therapy. **(A)** CD39 (*ENTPD1*), CD73 (*NT5E*), and A_2A_R (*ADORA2A*) row Z-score and expression in CD8^+^ T cells from scRNA-seq studies. **(B)** Representative flow cytometry plot and CD73 frequency on CD8^+^ T cells from HC (n = 42), untreated (n = 10), and treated (n = 6) patients with STAT3 GOF. **(C)** Representative histogram of A_2A_R expression on CD8^+^ T cells in HC (n = 18), untreated (n = 5), and treated (n = 4) STAT3 GOF patients. **(D)** Correlation of IFN-γ production by CD39, CD73, and A_2A_R by naïve CD8^+^ T cells. **(E)** Impact of treatment on IFN-γ cytokine production (n = 14 untreated STAT3 GOF; n = 8 treated STAT3 GOF patients). **(F)** Correlation of TNF-α production by CD39, CD73, and A_2A_R by naïve CD8^+^ T cells. **(G)** Impact of treatment on TNF-α cytokine production (n = 11 untreated STAT3 GOF; n = 6 treated STAT3 GOF patients). **(H)** Representative flow cytometry plots of naïve CD8^+^ T cell IFN-γ intracellular cytokine detection following *in vitro* αCD3/αCD28 activation in the presence or absence of ATP. (**I**) Paired analysis and quantification of percent inhibition in naïve CD8^+^ T cell IFN-γ production in αCD3/αCD28 activated cells in the presence of ATP. **(J)** Schematic of purinergic signaling deficits in STAT3 GOF patients and impact on CD8**^+^** T cell function. Blue square represents patient treated with Rapamycin rather than JAKi (which is shown as blue circles). For **I**: data are pooled from 2 independent experiments with 3 repeat HC and 1 repeat patient with STAT3 GOF; repeated samples across the two experiments were averaged into one data point per participant. A total of n = 17 HC (gray), n = 4 untreated (red), and n = 4 treated (blue) patients with STAT3 GOF were analyzed. \**P* ≤ 0.05, \*\*\*\**P* ≤ 0.0001 by Kruskal-Wallis test with Dunn’s multiple comparison test, Wilcoxon matched-pairs signed rank test, or Simple Linear Regression. Shaded area indicates 95% confidence interval. For comparisons >2 groups, adjusted p-values are indicated and ns or not listed were not statistically significant.

To test for a more direct connection between the observed decrease in the terminal components of the purinergic signaling cascade and the altered function previously quantified in naïve CD8^+^ T cells in STAT3 GOF (**Fig. 2B-C**), we correlated CD8^+^ T cell cytokine production levels with expression of CD39, CD73, and A_2A_R for patients with STAT3 GOF for whom we had both paired purinergic molecule phenotypic and functional data. Here, we found that the frequency of IFN-γ^+^ and CD39^+^ naïve CD8^+^ T cells positively correlated (R^2^ = 0.75, p-value < 0.0001) while the frequency of IFN-γ^+^ and CD73^+^ (R^2^ = 0.50, p-value = 0.0048) and A_2A_R (R^2^ = 0.36, p-value = 0.09) are negatively correlated (**Fig. 6D**). Additionally, patients with STAT3 GOF who were receiving JAKi demonstrated a trending decrease in IFN-γ production (p-value = 0.052) (**Fig. 6E**). The frequency of TNF-α^+^ naïve CD8^+^ T cells was also positively correlated (R^2^ = 0.64, p-value < 0.0005) with increased CD39 frequencies and inversely correlated with CD73 frequencies (R^2^ = 0.54, p-value < 0.0029) and A_2A_R expression (R^2^ = 0.42, p-value = 0.06) (**Fig. 6F)**. Here, again, we find that patients on JAKi demonstrate a significant decrease in TNF-α compared to untreated patients with STAT3 GOF (**Fig. 6G**). We also found that within total CD8^+^ T cells, the inverse relationship between CD73 and A_2A_R levels and IFN-γ (**Supp. Fig. 14D**) and TNF-α (**Supp. Fig. 14E**) cytokine production was statistically significant. Within non-naïve CD8^+^ T cells we observed an inverse correlation between IFN-γ^+^ and CD73^+^ cells (p-value = 0.01), a trending inverse correlation between IFN-γ^+^ cells and A_2A_R expression (p-value = 0.08) (**Supp. Fig. 14F**), and an inverse correlation between TNF-α^+^ and A_2A_R expression (p-value = 0.04) (**Supp. Fig. 14G**). Overall, downregulation of terminal components of the purinergic signaling cascade are associated with increased levels of pro-inflammatory cytokine production across CD8^+^ T cell subsets.

Having observed decreased frequencies of CD73^+^CD8^+^ T cells from patients with STAT3 GOF, we wanted to test whether STAT3 positively or negatively regulated CD73 expression *in vitro* in HC. Using our *in vitro* activation model of αCD3/αCD28 ± cytokines, we found that acute activation (96 hours) with αCD3/αCD28 reduced CD73 levels on HC CD8^+^ T cells (**Supp. Fig. 15A**). However, CD73 levels did increase, by both frequency and relative expression, after this short-term culture with αCD3/αCD28 ± IL-21 or IL-27, although not all, STAT3-activating cytokines (**Supp. Fig. 15A**). These acute culture conditions likely do not reflect the conditions in STAT3 GOF patients, who experience chronic STAT3 signaling *in vivo* from the initial differentiation of T cells from hematopoietic stem cells through their peripheral recirculation. In fact, recent work in mice has shown that CD73 is downregulated upon initial activation but is upregulated upon the resolution phase of an immune response in the setting of an infection, and during memory formation (*87*). Therefore, we tested whether more prolonged exposure of CD8^+^ T cells to STAT3-activating cytokines would increase CD73 levels after initial activation. Here, we cultured HC CD8^+^ T cells for 10 days, where for the first 4 days, cells were activated in the presence of αCD3/αCD28 alone, αCD3/αCD28 + IL-10, αCD3/αCD28 + IL-21, αCD3/αCD28 + IL-27, or αCD3/αCD28 + IFN-α (with both IL-27 and IFN-α also being strong inducers of STAT1) (**Supp. Fig. 11B**-C**, Supp. Fig. 15B**-C). On day 4 of activation, the media was exchanged, and cells were cultured with just cytokine alone (or control media) for an additional 6 days. Co-culture with IL-10 or IL-21 resulted in increased frequencies (**Supp. Fig. 15D**-E) and expression (**Supp. Fig. 15F**) of CD73^+^CD8^+^ T cells, compared to co-culture with IL-27 or IFN-α, where the activation induced reduction in CD73 expression was maintained (**Supp. Fig. 15D**-E). Moreover, co-culture with IL-27 or IFN-α, led to decreased frequencies of CD39^+^CD73^+^CD8^+^ cells (**Supp. Fig. 15E**). These data suggest that prolonged exposure to an inflammatory milieu that includes STAT1 activation in the context of STAT3 activation may inhibit the re-upregulation of CD73. This would be consistent with our *in vitro* IL-27 data (**Supp. Fig. 15A**), interferon signature from our scRNA-seq analysis (**Fig. 1I**), and inflammatory state observed in patients with STAT3 GOF (**Supp. Fig. 3A**) (*87*).

Lastly, since we observed decreased CD73 and A_2A_R levels on CD8^+^ T cells from patients with STAT3 GOF, we hypothesized that the immunoregulatory, anti-inflammatory, role of purinergic signaling may be dysfunctional in STAT3 GOF. To address this hypothesis, we tested the integrity of adenosine signaling in CD8^+^ T cells by assessing the impact of exogenous ATP (which can be metabolized to adenosine) or adenosine on cytokine production. PBMCs from HC, patients with STAT3 GOF who are not receiving JAKi, and patients with STAT3 GOF receiving JAKi were activated with αCD3/αCD28 for 24 hours in the presence or absence of ATP or 2-Chloroadenosine (ADO; agonist for A_2A_R), followed by staining for IFN-γ intracellular cytokine detection (**Fig. 6H**). We had previously observed hypersecretion of IFN-γ from naïve (CD45RA^+^CD27^+^) CD8^+^ T cells from patients with STAT3 GOF (**Fig. 2B**), so we focused our analyses of the impact of purinergic signaling on this CD8^+^ T cell subset.

We hypothesized that if purinergic signaling was impaired, it may contribute to the hypersecretion of inflammatory cytokines from naïve CD8^+^ T cells in STAT3 GOF. Relative to αCD3/αCD28 alone, PBMCs from HCs demonstrated a statistically significant reduced frequency of IFN-γ^+^ naïve CD8^+^ T cells when activated with αCD3/αCD28 + ATP, but this inhibition was not seen in naïve CD8^+^ T cells from STAT3 GOF (**Fig. 6I**). Naïve CD8^+^ T cells from untreated patients with STAT3 GOF demonstrated a trend towards less inhibition (average inhibition = 19.9%) relative to HC (average inhibition = 44.1%) in the presence of exogenous ATP. Naïve CD8^+^ T cells from patients receiving JAKi demonstrated a slight increase in inhibition relative to untreated patients with STAT3 GOF (average inhibition = 33.7%) (**Fig. 6I**). Similar effects were seen on IFN-γ MFI (**Supp. Fig. 16A**). Relative to αCD3/αCD28, only HC demonstrated a statistically significant inhibition in IFN-γ frequency and MFI when activated with αCD3/αCD28 + ADO (**Supp. Fig. 16B**-C). The overall IFN-γ inhibition in the presence of ADO was comparable across all three groups (**Supp. Fig. 16B**-C). Total (**Supp. Fig. 16D**) and non-naïve (**Supp. Fig. 16E**) CD8^+^ T cells demonstrated that only HCs were able to be inhibited in the presence of ATP or adenosine, but not in STAT3 GOF. Altogether, these data suggest that the CD39/adenosine immunoregulatory pathway is dysfunctional in patients with STAT3 GOF (**Fig. 6J**). Despite elevated levels of CD39 in STAT3 GOF patients, driven by STAT3 signal amplification, reduced expression of CD73 and A_2A_R shows evidence of contributing to the dysregulated, hyper-inflammatory nature of naïve STAT3 GOF CD8^+^ T cells. This axis and hypersecretion of inflammatory cytokines is partially normalized by JAKi treatment of patients with STAT3 GOF.

## DISCUSSION

STAT3 GOF is a rare, early onset and complex Primary Immune Regulatory Disorder (*8*, *9*) that causes autoimmunity, lymphoproliferation and recurrent infections and is characterized by CD8^+^ T cell dysregulation. This study demonstrates amplified inflammatory cytokine production by human naïve CD8^+^ T cells in STAT3 GOF and implicates dysregulation of purinergic signaling, with increased CD39 expression and decreased CD73 and A_2A_R expression on CD8^+^ T cells, which may directly contribute to the autoimmune and inflammatory components of this disorder. *In vivo* blockade of JAK/STAT signaling was assessed by evaluating CD8^+^ T cells from patients with STAT3 GOF on JAKi and demonstrated partial normalization of both purinergic signaling components and inflammatory cytokine production in naïve T cells. This is consistent with partial response to JAKi in some STAT3 GOF patients (*86*, *88*) and may imply a role for epigenetic changes in CD8^+^ T cells that remains to be investigated in future work.

Previous work has implicated hypoxia and aryl-hydrocarbon signaling in promoting CD39 expression (*70*, *89*) and there has been evidence of CD39 as a direct target of STAT3 in CD4^+^ T cells (*18*, *72*). Here, we show that STAT3 is a critical regulator of CD39 expression in human CD8^+^ T cells. Using *in vitro* model systems, we show that CD8^+^ T cells expressing the highest levels of CD39 also contain the highest pSTAT3 signaling. Additionally, we demonstrate that various STAT3-activating cytokines in the context of αCD3/αCD28 activation can individually induce higher levels of CD39 expression. However, both *in vitro* and *in vivo*, there could synergistic impacts of multi-cytokine exposure as has previously been demonstrated with joint IL-12/IL-27 stimulation *in vitro* leading to more robust CD39 expression, and contributing to immune dysregulation (*90*). Our *in vitro* STAT3 inhibition assays, comparison to patients with STAT3 DN variants, and retroviral work further support the role of STAT3 in regulating CD39 expression.

Mouse models of STAT3 GOF have demonstrated CD8^+^ T cells that are highly activated and cytotoxic, and contribute to pathogenesis of the autoimmunity (*30*, *31*). Similar to these mouse models, we observed evidence of inflammatory processes using single-cell RNA sequencing and plasma proteomic profiling in patients with STAT3 GOF (*25*, *30*). Functional assessment revealed that naïve CD8^+^ T cells from patients who are not on targeted therapy produced higher levels of IFN-γ and TNF-α compared to HC. While CD39 has served as a marker of the most dysfunctional CD8^+^ T cells in cancer and chronic viral infections, recent work also suggests that these cells can be endowed with suppressive capacity (*46*, *47*). The increased CD39 expression identified in our cohort of patients with STAT3 GOF in isolation could indicate greater negative tone via purinergic pathways on bystander cells. However, we found that downstream purinergic signaling molecules, CD73 and A_2A_R, were downregulated on CD8^+^ T cells from patients with STAT3 GOF transcriptionally and by protein levels using flow cytometry. Overall, these findings within CD8^+^ T cells suggest that reduced presence of downstream purinergic signaling mediators (CD73 and A_2A_R) may prevent proper negative regulation of CD8^+^ T cells in settings of chronic inflammation and could contribute to the hypersecretion of inflammatory effector cytokines (e.g., IFN-γ and TNF-α) in STAT3 GOF. Given the evidence in STAT3 GOF mouse models that CD8^+^ T cells are a key pathogenic cell type mediating autoimmunity, it is possible that this dysregulated pathway contributes to autoimmune pathophysiology in both mouse models and patients, although this remains to be studied further in the future. Previous studies have shown that persistent activation of T cells results in decreased expression of CD73 (*91*, *92*). We show that prolonged maintenance of cells in an inflammatory milieu that favors STAT1 activation, with concurrent STAT3 activation (e.g., IL-27 and IFN-α), decreases CD73 expression. This could contribute to persistently decreased CD73 expression in patients with STAT3 GOF. Additionally, it is unknown which STAT3 GOF variants affect heterodimerization with other STAT proteins (such as STAT1) (*74*) and whether altered heterodimerization could have an impact on the inflammatory milieu observed in patients with STAT3 GOF, which is an area for future research.

We tested the hypothesis that the purinergic axis lacks its intended immunoregulatory effect on CD8 T cells in this monogenic chronic inflammatory disease. Specifically, we assessed the ability of CD8^+^ T cells to be inhibited by the hydrolysis of eATP, or by adenosine itself. Indeed, although there was heterogeneity in the ability of STAT3 GOF CD8^+^ T cells to be inhibited *in vitro*, we observed trends that suggest patients with STAT3 GOF have decreased sensitivity to adenosine-mediated inhibition through the hydrolysis of ATP. While these studies focused on A_2A_R, expression of additional adenosine receptors with varying degrees of affinity (e.g., A1, A2B, A3) were not assessed in this study. This could contribute to the impact of adenosine supplementation on cytokine production. Assessment of these additional adenosine receptors warrants future investigation in this disorder. Here, we focused on purinergic molecule expression on CD8^+^ T cells. However, evaluation of CD73 and A_2A_R expression across non-T cell lineages demonstrated that, apart from monocytes which had a slight increase in CD73 expression, CD73 and A_2A_R expression was comparable between HC and patients. The impact of these alterations on monocytes differentiation and function was not assessed in these studies. Future studies should elucidate whether this inhibition occurs directly on CD8^+^ T cells or via collaboration amongst additional immune cells, such as T cells with monocytes (which express the highest level of A_2A_R) or dendritic cells and could particularly impact activation of CD8^+^ T cells, especially during these acute time points.

Lastly, we tested the impact of standard of care therapy for severe STAT3 GOF (e.g., JAKi) on CD8^+^ T cells from patients with STAT3 GOF (*86*). Here, we found that dysregulated cytokine production from naïve CD8^+^ T cells from patients with STAT3 GOF receiving JAKi was decreased compared to untreated patients with STAT3 GOF. Permanent changes to the epigenetic status of cells from untreated and treated patients with STAT3 GOF were not investigated here but would be useful in mechanistically dissecting the impact of pathway perturbation in these patients. However, our available phenotypic and functional data set leverages samples from the largest published patient cohort of STAT3 GOF, and we were able to assess correlations between expression of CD39/CD73/A_2A_R and cytokine production (*10*, *31*). Individuals with the least CD73 and A_2A_R demonstrated the highest level of IFN-γ and TNF-α production—significantly beyond the amounts secreted by HC and may be consistent with the demonstrated pathogenicity of CD8^+^ T cells in mouse models of STAT3 GOF.

Overall, this study implicates dysfunctional purinergic signaling in STAT3 GOF as a contributor to dysregulation of CD8^+^ T cell functionality, contributing to both immune dysregulation and autoimmunity. The understanding of how purinergic signaling affects CD8^+^ T cell functionality in the pathogenesis of autoimmunity may have broader implications for more common causes of chronic inflammation that increase STAT3 signaling.

### Study Limitations

One limitation of this study is the number of STAT3 GOF patients, which is limited both because this disease is rare, and thus obtaining access to STAT3 GOF patients is challenging, but in addition, obtaining samples from STAT3 GOF patients prior to treatment with immune modulatory therapy in this severe illness is also challenging. While we focused on the CD8^+^ T cell compartment in STAT3 GOF syndrome, most of our analyses in experiments that include primary patient samples were performed on PBMCs. This is the case because obtaining enough blood volume to purify sufficient STAT3 GOF CD8^+^ T cells for assays is difficult, especially as many of these patients are young children and many are lymphopenic. Moreover, we could only make correlations between cytokine production and purinergic pathway members for individuals for whom we have paired CD39/CD73/A_2A_R and cytokine data, which is a subset of our cohort. In addition, when working *in vitro*, the chronic exposure to amplified STAT3 signals *in vitro* in primary cells is limited compared to cells from patients with STAT3 GOF whose cells experience lifelong alterations in STAT3 signaling through differentiation and activation. While we focused on patients, and modulated the purinergic axis *in vitro*, future studies could examine the impact of modulating the CD39/CD73/adenosine axis pathway *in vivo* in mouse models to understand the dynamics of expression of these molecules as mice age and as disease worsens.

## MATERIALS AND METHODS

### Recruitment of healthy controls and patients with STAT3 GOF and STAT3 DN

In accordance with IRB approved protocols, participant or guardian consent (and assent, if appropriate) were obtained before study enrollment, as below. Participants with STAT3 GOF were recruited with institutional IRB approval (Washington University IRB #201107235; NIH, NIAID NCT 03394053 and NCT00404560; CHOP IRB #18-15920; Children’s National IRB #Pro00009689; CHOP IRB #15-01226; NIAID NIH IRB #05-I-0213; TCH IRBs: #21453 and #30487). Patients with STAT3 DN were recruited under NIH Clinicaltrials.gov number NCT00006150. HC were recruited under CHOP IRB #18-15920. Samples were shared via materials transfer agreements. Age, sex, STAT3 variant and medications for STAT3 GOF and DN are shown in **Table S1,** as is age and sex distribution for healthy controls.

### Mass cytometry

Antibodies for mass cytometry (**Table S4**) were conjugated in-house (using unlabeled, purified antibodies and conjugating them to isotope-loaded polymers with MAXPAR kits (Fluidigm)) or purchased as pre-conjugated metal-tagged antibodies from Fluidigm. After titration was performed, antibodies were diluted using antibody stabilization buffer (Candor Bioscience). Maleimido-mono-amine-DOTA (Macrocyclics) was dissolved in MAXPAR L-Buffer (Fluidigm) and mixed with 139La (L/D-139) for a 50 mM concentration. Cryopreserved PBMCs were thawed into warm complete RPMI (cRPMI: RPMI with 10% fetal bovine serum (FBS), 1% Penicillin/Streptomycin, 1% L-Glutamine and 1% HEPES) and plated at 1×10^6^ live PBMC/subject/well into 96-well U-bottom tissue culture plates. Cell suspensions were pelleted and incubated with 1.5 to 2.0 μM L/D-139 for 5 minutes at room temperature (RT) for Live/Dead discrimination. Cells were then washed (520xg, 5 min, RT) in staining buffer (1% FBS in phosphate-buffered saline (PBS)) and resuspended in 50μL surface antibody cocktail for 30 minutes at RT. They were then washed twice in staining buffer, fixed and permeabilized in 50μL Fixation/Permeabilization working solution (Foxp3 Transcription Factor Staining Buffer Set, eBioscience) and washed with Permeabilization Buffer (650xg, 5 min, RT). Samples were stained intracellularly (50μL intracellular antibody cocktail in Perm Buffer, 60 min, RT). Cells were washed three times (650xg, 5 min, RT) with Perm Buffer before fixation in 1.6% paraformaldehyde (PFA) (Electron Microscopy Sciences) in PBS overnight with 125nM Iridium (Fluidigm) at 4°C. Cells were washed twice in PBS (no Ca^2+^ or Mg^2+^) and then washed twice in deionized H_2_O prior to acquisition. Samples were acquired on a CyTOF2 (Fluidigm) with bead-based normalization. During data export, bead events were removed. FCS files were analyzed using FlowJo v10 (TreeStar).

### Antibody staining for flow cytometry

Approximately 0.25 to 0.5×10^6^ PBMCs per patient per stain were plated in a 96-well U-bottom plate (see **Table S5** for spectral flow cytometry antibody information). PBMCs were washed once with PBS and stained with either Live/Dead Aqua or Live/Dead Blue and Fc Block mix (50μL, 15 min, 4°C) prepared in PBS. PBMCs were washed again with PBS, and spun down (800xg, 4 min, RT). The pellet was resuspended in 50μL of surface stain antibody mix in FACS buffer (PBS containing 1% FBS, 2mM EDTA) plus 10% BD Brilliant Stain Buffer and then incubated at 4°C for 30 minutes. PBMCs were washed with FACS buffer, spun down, centrifuged and the pellet was resuspended in 125μL of FACS buffer for acquisition. For intracellular cytokine and transcription factor staining, the surface stain mix was washed off and samples were fixed and permeabilized by incubating in 50μL of Fix/Perm buffer (20 min, RT in the dark) and washed in Perm Buffer. PBMCs were stained using 50μL of intracellular antibody mix (diluted in Perm Buffer) for 1 hour at 4°C. After the incubation, samples were washed with Perm Buffer and fixed in 125μL of 1.6% PFA in PBS at 4°C overnight. The following day, samples were acquired.

### Intracellular cytokine detection

Freshly thawed PBMCs from patients and HC (0.5×10^6^) were seeded into 96-well U-bottom plates in 100μL of cRPMI. Cells were left unstimulated in the presence of 100μL BD GolgiPlug (containing Brefeldin A) and BD GolgiStop (containing Monensin) or were stimulated with 100μL PMA (100ng/mL) and Ionomycin (1μg/mL) with GolgiPlug and GolgiStop for 4 hours. To measure intracellular cytokines, surface and intracellular staining was performed as detailed above.

### Olink plasma proteomics

For HC and patients with STAT3 GOF, heparinized plasma (100μL/participant) was sent to Olink Proteomics (Uppsala, Sweden). Identification and measurement of proteins was performed with the Olink Explore 1536 or Olink Explore 384 Inflammation proximity extension assay platforms. Concentrations of proteins are provided as Log2-scaled Normalized Protein eXpression (NPX) units post-normalization and quality control (*61*, *93*).

### Cytokine-mediated pSTAT1 and pSTAT3 phospho-flow assay

Cryopreserved PBMCs were thawed as described above, counted, and stained with Fc Block, Live/Dead Aqua and surface stain antibodies combined in FACS Buffer containing 10% BD Brilliant Stain Buffer for 30 minutes at RT in the dark (50μL/1×10^6^ cells). Cells were washed and placed in plain RPMI and allowed to rest for 1hr at 37°C, 5% CO_2_. Cells were harvested, counted and resuspended at 5×10^6^ cells/mL in plain RPMI. Then 50μL (0.25×10^6^ PBMC) were plated in a 96-well U-bottom plate and further rested for 30 min at 37°C, 5% CO_2_. Cytokine solutions were prepared in plain RPMI and kept on ice. 50μL of plain RPMI or indicated cytokine solution was added to the cells. Cells were incubated for 20 minutes at 37°C, 5% CO_2_. Cells were immediately fixed with 100μL of 4% PFA (diluted in PBS) for 15 minutes at room temperature (2% PFA final in-well concentration). The plate was centrifuged, and the cells were resuspended in 150μL of ice-cold methanol and placed at -20°C overnight for permeabilization. The following morning, cells were centrifuged and washed 2X with 150μL of Perm Buffer followed by staining for phospho- proteins in Perm Buffer for 1hr at room temperature (50μL/well). After the incubation, samples were washed with Perm Buffer, pelleted, and resuspended in 125μL of 1.6% PFA (in PBS) for fixation. Samples were acquired the next day.

### CD39 induction and phospho-flow cytometry

96-well flat bottom plates were left uncoated or were coated with 1μg/mL anti-CD3 solution in sterile PBS (200μL) overnight at 4°C. Cryopreserved CD8^+^ T cells were thawed as described above and plated at 0.15×10^6^ live cells/well (50μL) in cRPMI after aspirating the coating solution and gently washing wells with 200μL PBS and aspirating the PBS wash. 1μg/mL (final concentration) soluble anti-CD28 (50μL) or media was added per well then either 50μL media or cytokine media (to achieve the final indicated concentrations) were added. Finally, 50μL media or STATTIC (0.5μM final concentration per well) was added for a final in-well volume of 200μL/well. Cells were incubated for 4 days at 37°C, 5% CO_2_ before surface staining and intracellular staining as described above. For phospho-flow cytometry, after the end of the incubation, 150μL of supernatants was removed, and cells were washed in PBS and then transferred to a 96-well U-bottom plate and pelleted. Cells were resuspended in a mixture of Fc Block, Live/Dead discrimination dye, and surface stain antibodies in FACS Buffer containing 10% BD Brilliant Stain Buffer for 30 minutes at RT (50μL/well). Surface stain was removed by washing with 150μL of FACS buffer and pelleting cells. Cell pellets were fixed by resuspending in 100μL of 2% PFA diluted in PBS for 15 minutes at RT. Cells were then pelleted and resuspended in 150μL of ice-cold methanol and permeabilized at -20°C overnight. Completion of intracellular phospho-protein detection was performed as described above.

### Cell sorting

Cryopreserved PBMCs were thawed as described above and 0.5 to 1×10^6^ cells were transferred to a new tube for washing with PBS. After PBS wash and centrifugation, cell pellets were incubated in 50μL of Live/Dead Aqua, CD14 V500, CD16 V500, CD19 V500, CD3 APC-R700, CD4 AF488 and CD8a BV605 antibodies in FACS Buffer with 10% Brilliant Stain Buffer for 30 minutes at RT. Cells were then washed with FACS Buffer and pellets were resuspended in 500μL FACS Buffer and filtered through a nylon mesh cell strainer (35µm) to remove clumps, debris, and prevent clogging. Live, single CD14^-^CD16^-^CD19^-^CD3^+^CD4^-^CD8^+^ cells were sorted into 1.5mL Eppendorf tubes containing RPMI + 10% FBS on a Cytek Aurora CS System (Cytek Biosciences, Fremont, CA). Removal of EDTA was achieved by washing sorted cells two times with RPMI + 10% FBS. Cells were counted prior to 10X processing for single-cell RNA sequencing.

### Single cell RNA sequencing (scRNA-seq)

Single cell analyses of sorted CD8^+^ T cells from PBMC samples from human participants (HC and STAT3 GOF) were performed using the 10X Chromium Next GEM Single Cell 5’ Kit v2 and Chromium Single Cell Human TCR Amplification Kit (10X Genomics, Pleasanton, CA). We worked with the Center for Applied Genomics at Children’s Hospital of Philadelphia for single-cell isolation and library preparation. An Illumina S2 flow cell (Illumina, San Diego, CA) was used for sequencing. Sequencing basecall files were demultiplexed with 10x Genomics Cell Ranger v6.1.2’s makefastq (*52*). Gene expression fastq files were mapped to the human genome (GRCh38) and TCR fastq files were mapped to the CellRanger V(D)J reference (GRCh38, ensembl 7.1.0) using 10x Genomics Cell Ranger v7.1.0’s multi (*52*). Cellranger mapped h5 files were read in to and processed in R using Seurat v4 (*53*). Samples were assessed for quality and cells with less than 500 total reads, 300 total features, or more than a 5% mitochondrial percentage were filtered out. Doublets were also detected and removed with the DoubletFinder package (*54*). Cells from each sample were integrated together using the top 3000 most variable features and Seurat v4’s IntegrateData function (*53*). The integrated Seurat object was clustered with Seurat’s FindNeighbors and FindClusters functions (*53*). Resolutions ranging from 0.05 to 0.9 were assessed with clustree (*55*). A resolution of 0.3 was optimal and resulted in 10 clusters. These ten clusters were then annotated using a mixture of automated and manual methods. First, the Monaco reference dataset from the celldex package was applied, giving an initial set of annotations (*56*, *57*). The clusters were further examined using a select set of markers. One cluster was classified as monocytes and were filtered out. Another cluster was identified as low quality CD8^+^ T cells and removed. The rest of the annotations were defined based on the initial Monaco annotation and the expression of the markers of interest. Propeller in the speckle package was used to assess independence of the cell type annotations and patient disease status (*58*). Remaining CD8^+^ T cells were examined without the MAIT cell cluster. Differential expression among the genes was conducted by first pseudobulking cell counts per individual sample using Seurat’s AggregateExpression function (*53*). EdgeR was used to process the data and run the differential expression analysis (*94*). Counts were also normalized and log2 scaled using edgeR’s cpm function to look for potential gene set enrichment using the Broad Institute and UC San Diego’s GSEA (*94*). A curated list of gene sets were examined in the STAT3 GOF patients (*59*, *60*). Single-cell related figures including the dimension plot, dotplot, and volcano plots were made with a combination of built in Seurat functions and ggplot2 (*53*, *95*).

### scRNA-seq pipeline

*scRNA-seq pipeline:*
| Step | Tool | Summary | References |
| --- | --- | --- | --- |
| Demultiplexing | • bcl2fastq<br>• cellranger makefastq | Initial basecall (bcl) files were demultiplexed with cellranger's makefastq command (wrapper of Illumina's bcl2fastq tool). | 10x Genomics Cell Ranger v6.1.2<br>(52) |
| Read Mapping | • cellranger multi | Gene expression files were mapped to the human genome GRCh38 build) and TCR files were mapped to the CellRanger V(D)J reference (GRCh38, ensemble 7.1.0). |  |
| Initial Processing + QC | • Seurat v4 | Cellranger's h5 files were read into R to be processed using Seurat v4. Samples were filtered using minimum 500 total reads per cell, minimum 300 total features per cell, and a 5% maximum mitochondrial percentage. Doublets were detected and filtered using the DoubletFinder package. | (53, 54) |
| Integration + Clustering | • Seurat v4<br>• clustree | Samples were integrated together using Seurat v4's IntegrateData function and the top 3000 most variable features. Clustering resolutions 0.05 – 2.0 were tested and examined with the clustree package. A resolution of 0.3 was chosen which defined 10 clusters. | (55) |
| Annotation | • Seurat v4<br>• Celldex 1.15.0<br>• Propeller | Clusters were annotated using a mix of manual and automated methods. The Monaco reference in the celldex package was applied to the 10 defined Seurat clusters. The automated annotations were examined further using a select set of markers (dotplot). Final cluster annotations were defined using both the initial Monaco annotation and the marker expression. One cluster was identified as contaminating monocytes and was excluded from downstream analyses due to high levels of *CD14* while another cluster was identified as low-quality cells (low nCounts, nFeatures, high mitochondrial percentage) and filtered out. The remaining 8 clusters of CD8<sup>+</sup> T cells were used for analysis including differences in proportions. | (56–58) |
| Differential Expression + GSEA | • Seurat v4<br>• edgeR<br>• ggplot2<br>• GSEA 4.3.2 | CD8<sup>+</sup> T cells were further examined without the MAIT cluster. Genes in STAT3 GOF patients and HCs were compared in the CD8<sup>+</sup> T cells using Seurat v4's FindMarkers function with the Wilcoxon method being the test used. Counts were combined per sample using Seurat's AggregateExpression function. Aggregated counts were then normalized and log2 scaled with edgeR's cpm function. Gene set enrichment was assessed using the Broad Institutes GSEA. A select list of gene sets was analyzed (including the Human MSigDB v2023.2.Hs Hallmark gene sets).<br>Figures were generated using ggplot. | (59, 60, 94, 95) |

### Re-analysis of single cell RNA sequencing from Warshauer et al., 2021

Publicly available single cell gene expression data from the Warshauer et al., 2021 paper was re-examined in a similar manner to the original publication (*30*). The alignment h5 files were imported into R and analyzed with Seurat v4 (*53*). Cells below 200 gene features per cell, 500 total reads per cell, or more than a 5% mitochondrial percentage were filtered out. Data was integrated together using Seurat v4’s IntegrateData function (*53*). T cells were identified using a mix of manual examination of expression of *Cd8a* and the automatic cluster identifications from the Monaco reference data (*56*, *57*). Differential expression between STAT3 GOF and control CD8^+^ T cells was analyzed using Seurat’s FindMarkers function as it was in the initial publication (*30*, *53*).

### Chromatin immunoprecipitation sequencing re-analysis

ChIP-seq datasets for STAT1 and STAT3 were downloaded from the NCBI Gene Expression Omnibus (GSE65621) (*74*). We have included data for CD4^+^ T cells cultured *in vitro* with or without IL-27. 50 base pair reads were aligned using bowtie and macs2 (default settings) was used to map non-redundant reads to mouse genome (mm9). Input controls were used as references for peak calling. Homer was used for peak annotation and DNA motif enrichment analysis. We used bedtools intersect for peak overlap analysis (-wa option). Gene proximal peaks were defined as occurring within introns, exons, being less than 10 kb from transcriptional start or end sites or in untranslated regions. IGV was used to render genome browser files.

### PBMC inhibition with ATP and adenosine

Live PBMCs (0.15×10^6^, 50μL) were plated per well in a flat-bottom 96-well plate pre-coated with 200μL of 1μg/mL anti-CD3 in PBS. Media or soluble anti-CD28 solution (50μL, final concentration = 1μg/mL) was added alongside 100μL media, ATP (final concentration = 50μM) or 2-choloroadenosine (ADO, final concentration = 5μM). Cells were incubated for 24 hours then supernatants were harvested and frozen, followed by addition of Brefeldin A/Monensin for an additional 4 hours. Then, surface and intracellular cytokine staining was performed as described above. Percent inhibition in the presence of CD3/CD28 + ATP or CD3/CD28 + ADO relative to CD3/CD28 only activated cells was calculated as follows for each cell population:

% Inhibition = ((%IFN-γ^+^_CD3/CD28 Only_ - %IFN-γ^+^_CD3/CD28 + ATP or ADO_)/(%IFN-γ^+^_CD3/CD28 Only_))*100%

### Mouse models of STAT3 GOF

STAT3^+/G421R^ mice and littermate controls were maintained at Washington University in St. Louis under previously established guidelines and approvals (*25*). NOD-STAT3^+/K392R^ mice and littermate controls were maintained at the University of California, San Francisco animal facility (specific pathogen-free) under previously established guidelines and approvals (*30*). Generation of *Stat3* flox/flox CD4-Cre(+) and *Stat3* flox/flox CD4-Cre(-) littermate controls was performed as described previously (*85*). Mice were housed and handled in accordance with NIH guidelines and all experiments were approved by the University of Miami Animal Care and Use Committee.

### Mouse splenocyte processing

Spleens were harvested from indicated mouse strains and shipped overnight in R10 media (RPMI + 10% FBS + 1% Penicillin/Streptomycin and 1% L-Glutamine) on ice packs. On arrival, tissues were weighed and mechanically dissociated by passing through a 70μm cell strainer with the back of a 1mL syringe. The strainer was regularly flushed with cold FACS buffer. Cells were pelleted at 4°C (1600 RPM x 5 min) and resuspended in 4mL of ACK Lysis Buffer, vortexed, and incubated for 8 min to lyse red blood cells, followed by quenching with 16mL of R10 media. Cells were counted with Nexcelom Cellometer K2 Image Cytometer Automated Cell Counter and after pelleting, cells were resuspended at 1×10^7^ live splenocytes/mL in mouse cRPMI (RPMI + 10% FBS, 1% Penicillin/Streptomycin, 1% L-Glutamine, 1% HEPES, 1% Non-Essential Amino Acids and 50μM β-ME). 100μL of cells were used per mouse per assay.

### Retroviral transduction

cDNA encoding WT (ENSMUST00000127638.8) or mutant (c.1032G>C, p.Q344H) mouse Stat3 were cloned into an MoMLV-based plasmid vector (MigR1) directly in upstream of the internal ribosome entry sequence (IRES) and GFP marker. Plasmid vectors were then transfected into 293T cells along with pCL-Eco helper plasmid (*96*). Supernatants containing virus were harvested after 48 hours. In parallel, naive CD44^Low^CD25^-^CD4^+^ or CD8^+^ T cells were sorted (purity > 95%) from pooled spleen and lymph nodes of *Stat3* flox/flox CD4-Cre(+) and *Stat3* flox/flox CD4-Cre(-) mice. Cells were activated with plate-bound anti-CD3e/CD28 (10μg/mL each) for 24 hours and then incubated with viral supernatant for 1 hour (2200 rpm, 18°C). Following transduction, cells were cultured *in vitro* for a further 48 hours with recombinant murine IL-27 (10ng/μL), anti-mouse IL-4 (10μg/mL) and anti-mouse IFN-γ (10μg/mL), then stained for flow cytometry. Cytometry data were collected on an Aurora instrument (Cytek Biosystems).

### Flow cytometry

Samples were acquired on a five-laser Cytek Aurora at the Children’s Hospital of Philadelphia Flow Cytometry Core, except for the retroviral research, which was performed at the Flow Cytometry Shared Resource of the Sylvester Comprehensive Cancer Center at the University of Miami (RRID: SCR022501). SpectroFlo® QC Beads were used for routine performance tracking. UltraComp eBeads were used for compensation. Data were analyzed using FlowJo software (version 10.9.0, Tree Star, Ashland, OR).

### Statistical Analysis

All statistical analyses were performed using GraphPad Prism (version 10.4.1 GraphPad Software, La Jolla, CA, USA). When HC and/or patient participant’s samples were run in more than one batch of an individual assay, all values obtained for that participant across the batches of that assay were averaged into one data point for each individual participant for plotting and analysis. Statistical significance among groups was calculated with Mann-Whitney test, analysis of variance (ANOVA), linear regression, or Wilcoxon matched-pairs signed rank test depending on the design of each experiment and distribution of the data. The number of samples analyzed and exact statistical test for each experiment is listed in the figure captions or in the figures; for analysis of more than 2 groups, adjusted p-values were used.

### Diagrams and schematics

All schematics were created using BioRender.com. Agreement number: MZ27JWZQUI (Created in BioRender. Campos, J. (2024) https://BioRender.com/ n45g327), Agreement number: FD27JX02BR (Created in BioRender. Campos, J. (2024) https://BioRender.com/ n88d277), Agreement number: QT27JX0SBF (Created in BioRender. Campos, J. (2024) https://BioRender.com/ p28i069), Agreement number: FO27JX145G (Created in BioRender. Campos, J. (2024) https://BioRender.com/ z66z909), Agreement number: KS27JX1HF0 (Created in BioRender. Campos, J. (2024) https://BioRender.com/ t14h362), Agreement number: HH27JX1VBV (Created in BioRender. Campos, J. (2024) https://BioRender.com/ u59z820).

## Acknowledgements

We would like to thank the patients and their families for participating. We would like to thank the members of the Children’s Hospital of Philadelphia Biorepository Resource Center Specimen Processing Unit (Richard Tustin III, Annemarie Butler, Vanessa Oliva) for help in processing PBMCs and plasma from samples, and Margaret Abaandou for assistance with patient biospecimens. Additionally, we also thank the Children’s Hospital of Philadelphia (CHOP) Flow Cytometry Core members for their assistance with flow cytometry experiments. We additionally would like to thank Dr. Christopher Hunter, Dr. Kathleen E. Sullivan, Dr. Laurence Eisenlohr, and Dr. Will Bailis for their scientific discussions, critical reading, and suggestions on this manuscript. All diagrams and schematics were created with BioRender.com.

## Funding

Howard Hughes Medical Institute Gilliam Fellowship for Advanced Studies GT15736 (JSCD) Penn Presidential Ph.D. Fellowship (JSCD)

NIH NIAID P01AI155393 (EGS, MAC)

The Jeffrey Modell Diagnostic and Research Center for Primary Immunodeficiencies at St. Louis Children’s Hospital, and the Children’s Discovery Institute of Washington University’s Center for Pediatric Immunology and St. Louis Children’s Hospital (EGS, MAC)

NIH NIAID 1K08AI182483 (EGS) NIH NIAMS P30AR073752 (EGS) NIH NIDDK K12DK133995 (MAT)

JDRF Advanced Postdoctoral Fellowship (MAT) Intramural Research Program of the NIAID (HCS) NIH with 1ZIAAI001059 (HCS)

Jeffrey Modell Foundation (LRFS)

NIH NCATS 5UG3TR003908-02 (LRFS) NIH NIAID AI146026 (NR)

NIH NIAID AI179680 (NR)

NIH NIAID P01AI155393 (MSA)

Arthritis National Research Foundation (TPV) NIH NIAID 1K08AI135091 (SEH)

Burroughs Wellcome Fund CAMS (SEH) The Hartwell Foundation (SEH)

Immune Deficiency Foundation (SEH)

Primary Immune Deficiency Treatment Consortium (SEH)

## Author contributions

Conceptualization: SEH, JSCD Methodology: JSCD, SEH

Investigation: JSCD, SS, MCD, AAM, MSK, PEC, ABS, CAH, RBL, CAH

Bioinformatics: MSK

Patient Sample Acquisition: EDA, SB, CMS, TCP, AT, IL, AG, SWC, HCS, MDK, OMD, LRFS, SMH, JREB, JWL, NR, AFF, MAC, TPV, SEH

Mouse Samples: EGS, MAT, MSA, MAC Retroviral Investigations: ABS, AVV Data Curation: JSCD, SEH

Formal Analysis: JSCD, SEH Visualization: JSCD, SEH

Writing – Original Draft Preparation: JSCD, SEH Writing – Reviewing & Editing: All Authors Funding Acquisition: SEH

Supervision: SEH

## Data and materials availability

All data will be available upon reasonable request. Request for reagents or resources should be directed to the corresponding author. Single-cell RNA-sequencing data will be deposited at the time of publication.

