## Supplementary Materials for "Altered Purinergic Signaling and CD8^+^ T Cell Dysregulation in STAT3 GOF Syndrome"

**The Supplementary Materials file includes the following:**

Fig. S1. Gating strategies and additional characterization of immune cell perturbations in STAT3 GOF in $CD4^+$ T cells, NK cells, B cells, and Monocytes.
Fig. S2. Hallmark Gene Set Enrichment Analysis (GSEA) leading-edge genes in STAT3 GOF.
Fig. S3. Olink proteomic analysis of healthy control and STAT3 GOF patients with signs of systemic inflammation.
Fig. S4. Gating strategy for intracellular cytokine detection on $CD8^+$ T cells.
Fig. S5. Mass cytometry quantification of CD39 expression across immune cell lineages.
Fig. S6. Spectral flow characterization of TOX, NKG2D, and CD57 expression in $CD8^+$ T cells.
Fig. S7. ChIP-Seq analysis of STAT3 binding at *Entpd1* locus in mouse $CD4^+$ T cells.
Fig. S8. Additional characterization of $CD8^+$ T cell compartment in $STAT3^{+/G421R}$ model.
Fig. S9. Additional characterization of $CD8^+$ T cell compartment in NOD-$STAT3^{+/K392R}$ model.
Fig. S10. scRNA-Seq re-analysis of murine $CD8^+$ T cells for CD39 expression in NOD-$STAT3^{+/K392R}$ model.
Fig. S11. Characterization of STAT1 and STAT3 signaling in $CD8^+$ T cells by various cytokines.
Fig. S12. CD39 induction in patients bearing STAT3 DN variants.
Fig. S13. CD73 and A2AR across immune cell lineages by spectral flow cytometry in STAT3 GOF.
Fig. S14. Additional CD73 and $A_{2A}R$ expression and cytokine correlations by $CD8^+$ T cell subsets.
Fig. S15. Role of STAT3 in regulating CD73 expression in $CD8^+$ T cells.
Fig. S16. Additional quantification of cytokine production inhibition in total, naïve, and non-naïve $CD8^+$ T cells after activation in the presence of ATP or adenosine.

Table S1. Healthy control, STAT3 GOF, and STAT3 DN participant metadata.
Table S2. List of DEGs in conventional $CD8^+$ T cells from human scRNA-seq analysis.
Table S3. GSEA result summary in conventional $CD8^+$ T cells.
Table S4. Mass cytometry antibodies.
Table S5. Flow cytometry antibodies.
Table S6. Additional key reagents/resources.

### Supplemental Figures:

#### Supp. Fig. 1

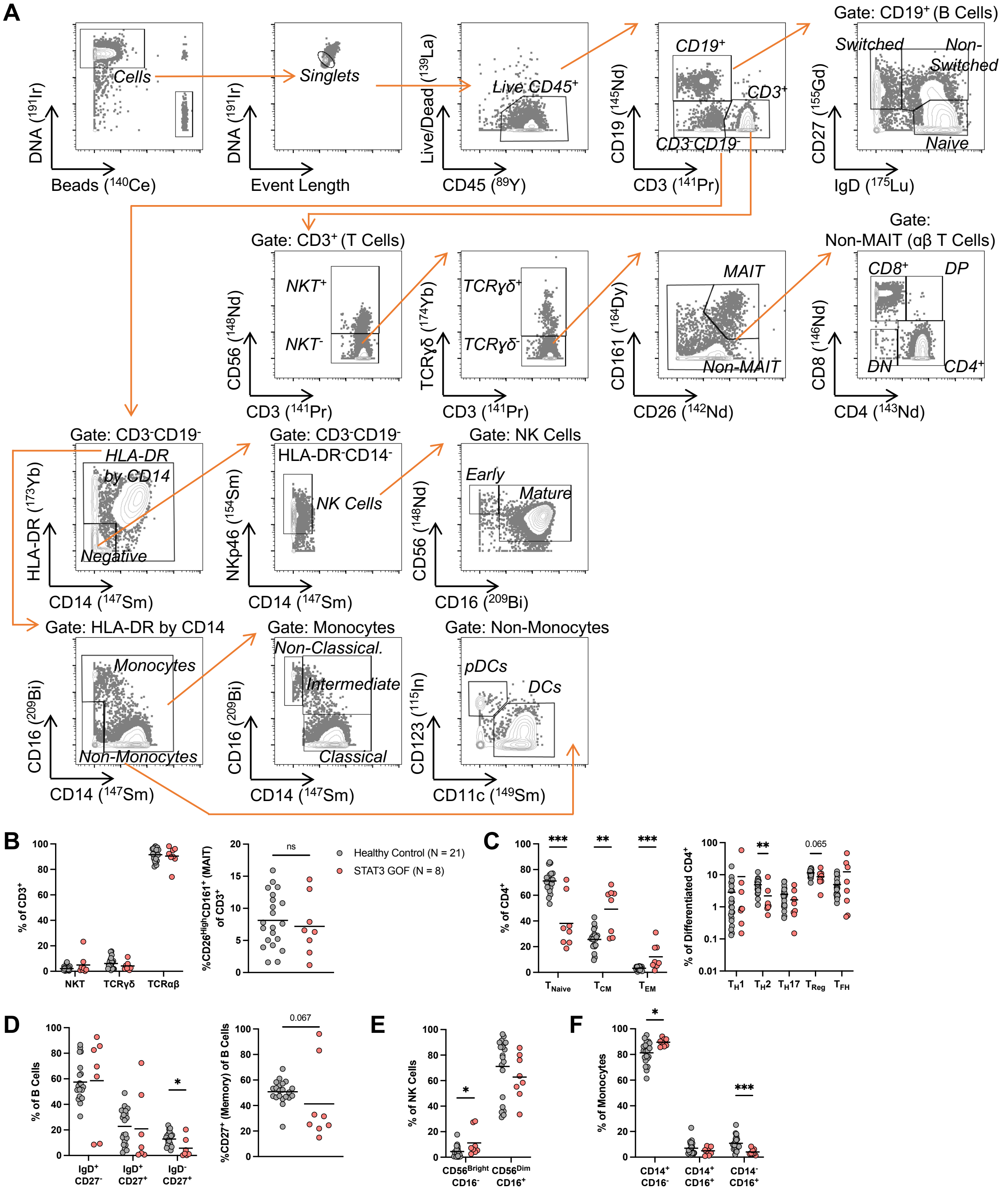

**Supplemental Figure 1: Gating strategies and additional characterization of immune cell perturbations in STAT3 GOF in CD4$^{+}$ T cells, NK cells, B cells, and Monocytes. (A)** Mass cytometry (CyTOF) gating strategy and frequencies of major immune cell populations in untreated STAT3 GOF patients (n = 8) and age-matched HC (n = 21). **(B)** Frequencies of NKT (CD3$^{+}$CD56$^{+}$), TCR$\gamma\delta$ (CD3$^{+}$CD56$^{-}$TCR$\gamma\delta^{+}$) and TCR$\alpha\beta$ (CD3$^{+}$CD56$^{-}$TCR$\gamma\delta^{-}$) and MAIT (CD3$^{+}$CD26$^{+}$CD161$^{+}$) cells. **(C)** Differentiation of CD4$^{+}$ T cells and frequencies of non-naïve CD4$^{+}$ T helper cell subsets as indicated.: $T_{H}1$: T-bet$^{+}$GranzymeB$^{+}$, $T_{H}2$: CD26$^{Int}$CRTH2$^{+}$, $T_{H}17$: CD26$^{High}$CRTH2$^{-}$, $T_{REG}$: CD127$^{Low}$FoxP3$^{+}$ and $T_{FH}$: PD-1$^{High}$CXCR5$^{+}$. **(D)** Frequencies of B cell populations: naïve: IgD$^{+}$CD27$^{-}$, non-switched memory: IgD$^{+}$CD27$^{+}$, switched memory: IgD$^{-}$CD27$^{+}$ and total memory (CD27$^{+}$) cells. **(E)** Frequencies of NK cell populations: immature: CD56$^{Bright}$CD16$^{-}$ and mature: CD56$^{Int}$CD16$^{+}$. **(F)** Frequencies of monocyte populations: non-classical: CD16$^{+}$CD14$^{-}$, intermediate: CD16$^{+}$CD14$^{+}$, and classical: CD16$^{-}$CD14$^{+}$. Data are pooled from 3 independent experiments. *$P \leq 0.05$, **$P \leq 0.01$, ***$P \leq 0.001$ by Mann-Whitney Test.

**Supp. Fig. 2**

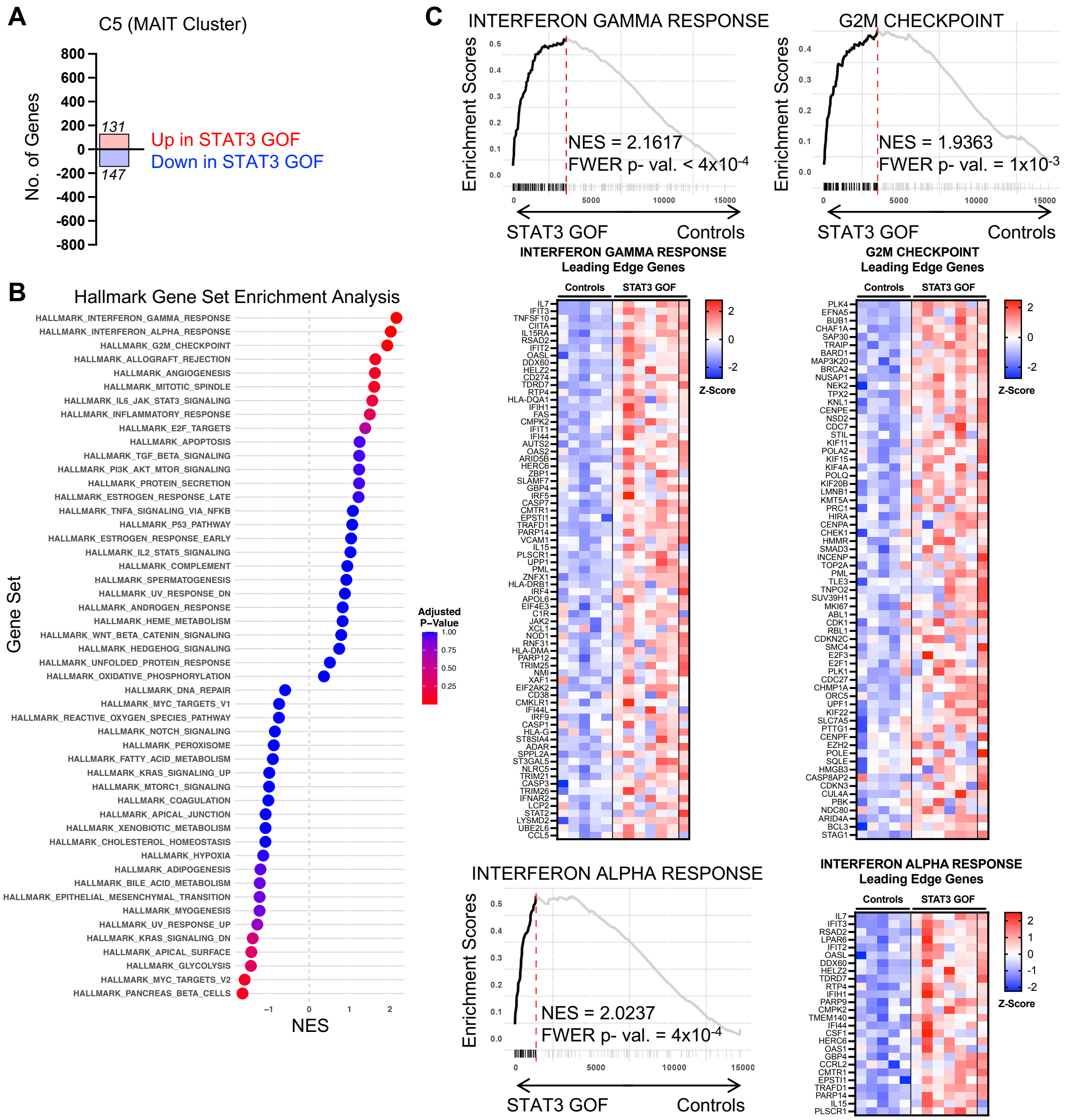


**Supplemental Figure 2: Hallmark Gene Set Enrichment Analysis (GSEA) leading-edge genes in STAT3 GOF. (A)** Bar plot showing number of differentially expressed genes upregulated or downregulated in C5 (MAIT cell cluster) from scRNA-seq analysis. **(B)** Summary results from MSigDB Hallmark GSEA. Positive NES indicates enrichment in STAT3 GOF. **(C)** Enrichment plots and heatmaps/annotation of leading-edge genes (ordered by Rank Metric Score) of statistically significant pathways enriched in STAT3 GOF. For GSEA analysis, correction was performed for multiple comparisons with adjusted p-values being indicated by color; statistical significance was deemed as adjusted p-value ≤ 0.05.

**Supp. Fig. 3**

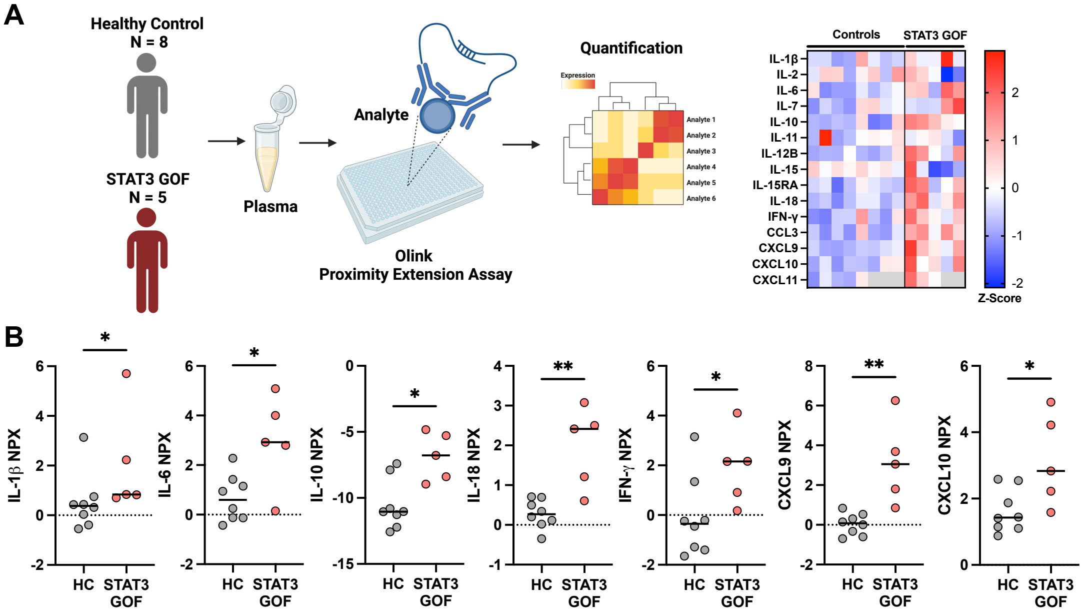


**Supplemental Figure 3: Olink proteomic analysis of healthy control and patients with STAT3 GOF with signs of systemic inflammation. (A)** Schematic of testing strategy follow by heatmap showing row normalized protein expression of select biomarkers from heparinized plasma from 8 HC (n = 8) and patients with STAT3 GOF (n = 5). Gray box indicates analyte not analyzed. **(B)** $Log_2$ normalized protein expression (NPX) quantification of select biomarkers. *$P \leq 0.05$ and **$P \leq 0.01$ by Mann-Whitney Test.

**Supp. Fig. 4**

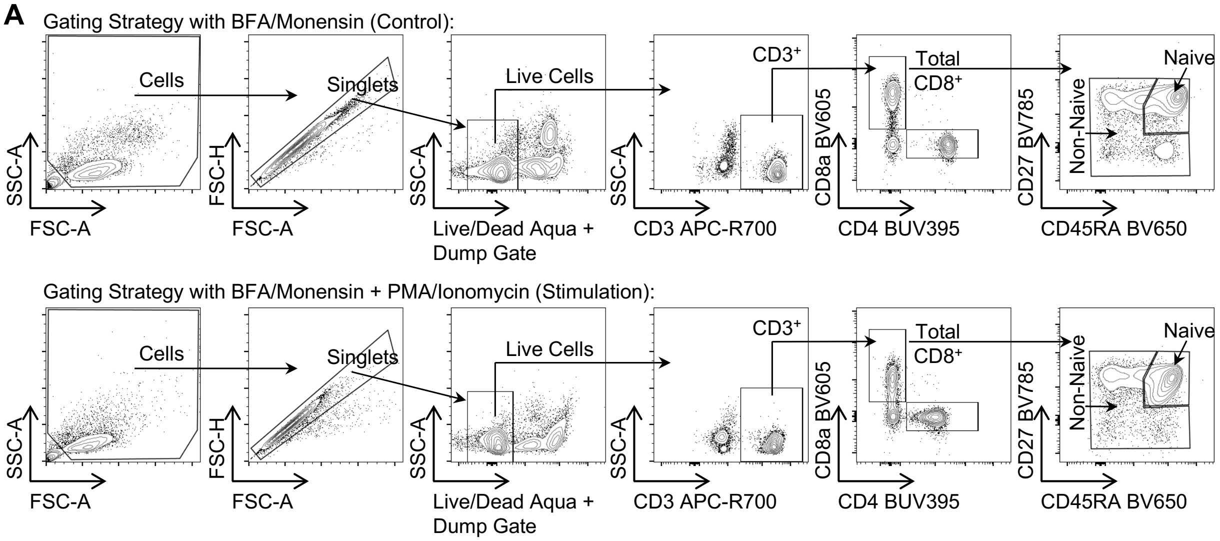


**Supplemental Figure 4: Gating strategy for intracellular cytokine detection in CD8$^{+}$ T cells**. **(A)** Gating strategy for spectral flow cytometry intracellular cytokine detection/quantification in the presence of BFA/Monensin or BFA/Monensin plus PMA/Ionomycin. Naïve CD8$^{+}$ T cells are defined as CD45RA$^{+}$CD27$^{+}$ while non-naive cells (antigen-experienced) are defined as non-CD45RA$^{+}$CD27$^{+}$ (L-shape encompassing CD45RA$^{-}$CD27$^{+}$, CD45RA$^{-}$CD27$^{-}$ and CD45RA$^{+}$CD27$^{-}$).

**Supp. Fig. 5**

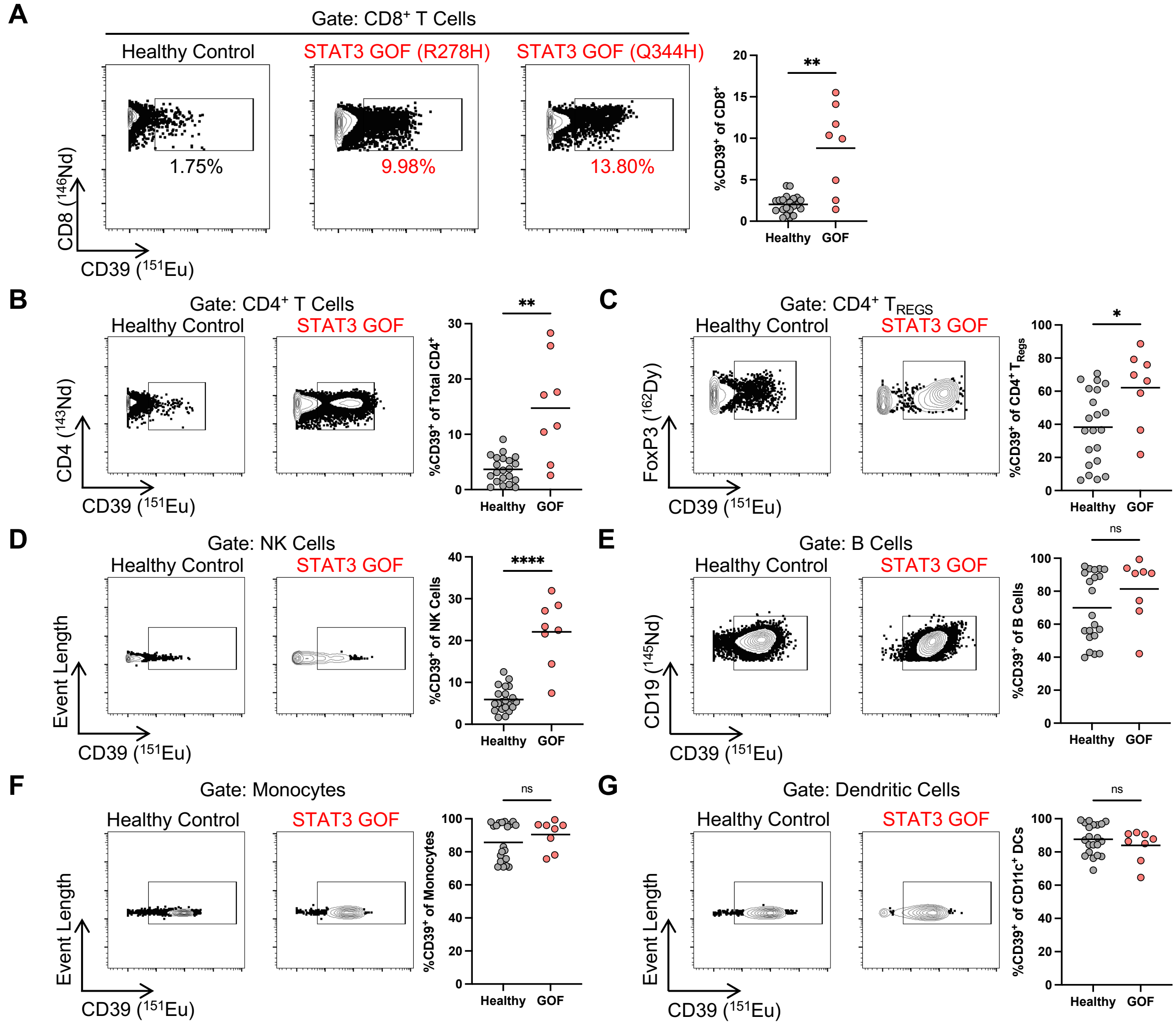


**Supplemental Figure 5: Mass cytometry quantification of CD39 expression across immune cell lineages.** Representative mass cytometry and quantification of CD39 frequency between HC (n = 21) and patients with STAT3 GOF (n = 8) in: **(A)** CD8$^{+}$ T cells, **(B)** CD4$^{+}$ T cells, **(C)** TREGS, **(D)** NK cells, **(E)** B cells, **(F)** monocytes and **(G)** CD11c$^{+}$ dendritic cells. Data are pooled from 3 independent experiments. *$P \leq 0.05$, **$P \leq 0.01$, ****$P \leq 0.0001$ by Mann-Whitney Test.

Supp. Fig. 6

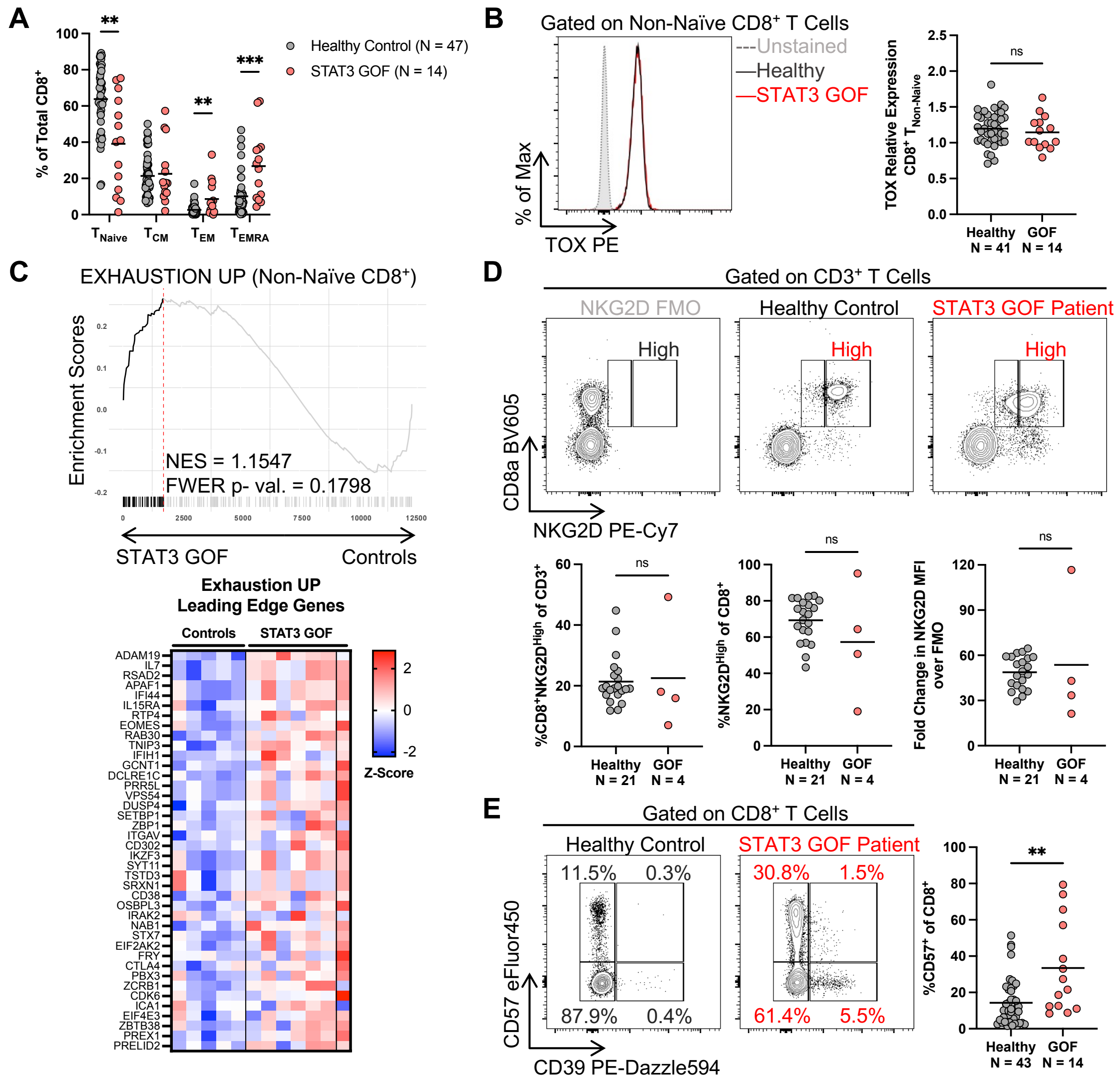


**Supplemental Figure 6: Characterization of TOX, NKG2D, and CD57 expression in CD8$^{+}$ T cells.** **(A)** Frequencies of CD8$^{+}$ T cell subsets from spectral flow cytometry assays. **(B)** Representative flow cytometry plot of TOX expression and quantification of relative expression among non-naïve CD8$^{+}$ T cells compared to the same well characterized HC in each experiment. **(C)** Enrichment plots and annotation of leading-edge genes (ordered by Rank Metric Score) of T cell exhaustion GSEA on non-naïve CD8$^{+}$ T cell clusters in STAT3 GOF compared to HC. **(D)** Representative flow cytometry plots and frequencies of NKG2D$^{High}$CD8$^{+}$ (within CD3$^{+}$ T cells), NKG2D$^{High}$ (within CD8$^{+}$ T cells), and fold change in NKG2D$^{High}$ MFI over FMO on CD8$^{+}$ T cells. **(E)** Representative CD57 and CD39 co-expression flow cytometry plots and CD57 frequency quantification. For **A-B, E**: data are pooled from >5 independent experiments. For **D**: data are pooled from 2 independent experiments. Statistical significance indicated above each plot and was calculated by Mann-Whitney Test.

**Supp. Fig. 7**

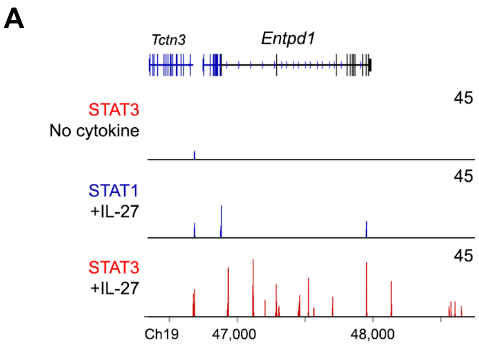


**Supplemental Figure 7: ChIP-Seq analysis of STAT3 binding at *Entpd1* locus in mouse CD4$^{+}$ T cells. (A)** Chromatin immunoprecipitation (ChIP)-Seq analysis of publicly available data demonstrating CD39 as a STAT3 target in mouse CD4$^{+}$ T cells stimulated with IL-27.

**Supp. Fig. 8**

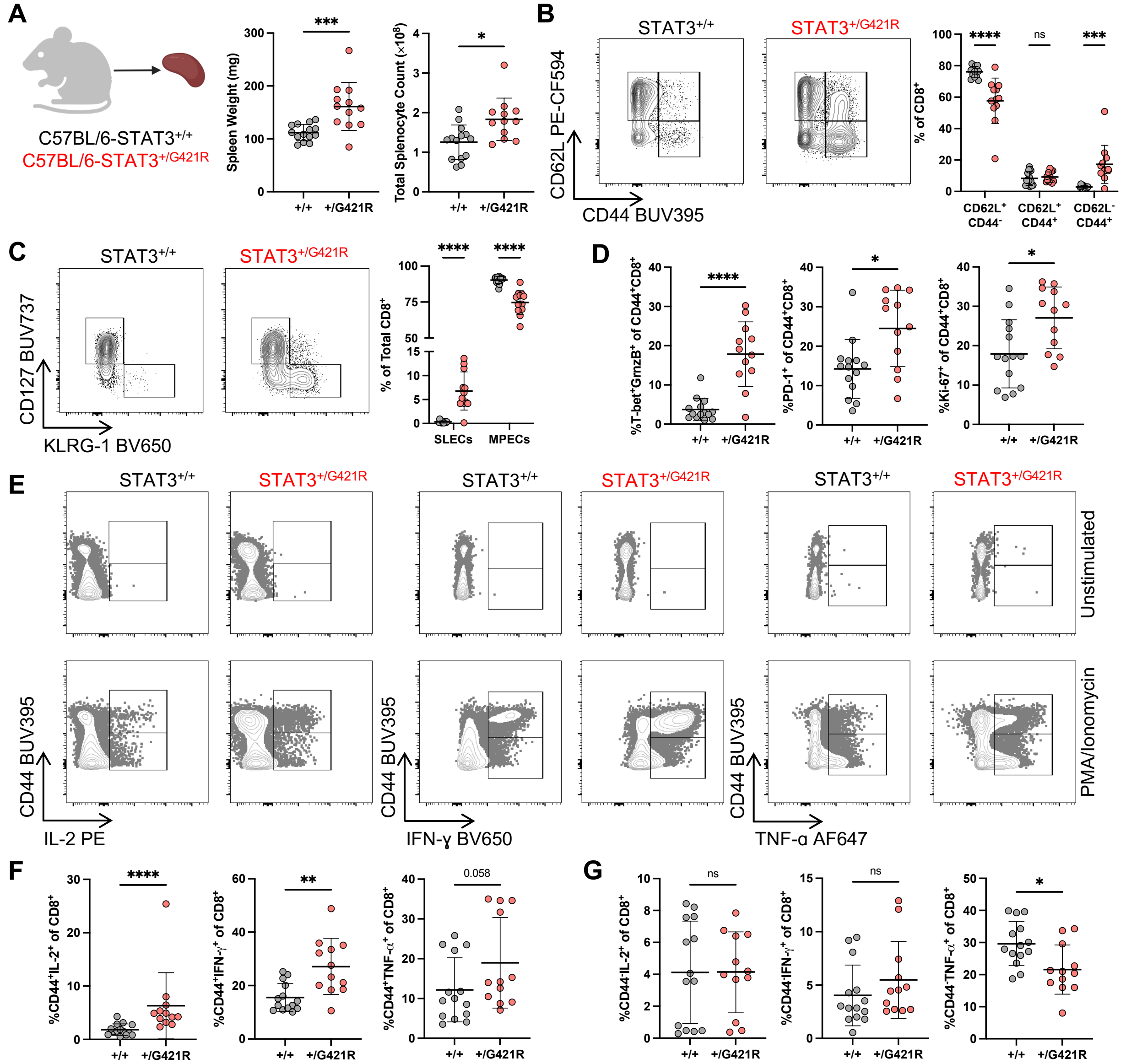


**Supplemental Figure 8: Additional characterization of $CD8^+$ T cell compartment in $STAT3^{+/G421R}$ model. (A)** Comparison of spleen weight and total cell numbers in the spleen of $STAT3^{+/+}$ (n = 14) and $STAT3^{+/G421R}$ (n = 12) mice. **(B)** Representative $CD8^+$ T cell subsets and quantification in $STAT3^{+/+}$ and $STAT3^{+/G421R}$ mice. **(C)** Representative MPEC ($CD127^+KLRG\text{-}1^-$) and SLEC ($CD127^-KLRG\text{-}1^+$) populations and quantification in $STAT3^{+/+}$ and $STAT3^{+/G421R}$ mice. **(D)** Quantification of activation markers on $CD44^+CD8^+$ T cells in $STAT3^{+/+}$ and $STAT3^{+/G421R}$ mice. **(E)** Representative flow cytometry plots of cytokine production by $CD8^+$ T cells after 5 hours with PMA/Ionomycin. **(F)** Percent $cytokine^+CD44^+CD8^+$ or **(G)** $cytokine^+CD44^-CD8^+$ T cells. Assay performed in total splenocytes and then gated on live $mCD45^+B220^-Gr\text{-}1^-CD3^+CD8^+$ cells. Data are pooled from 3 independent experiments. $*P \le 0.05$, $**P \le 0.01$, $***P \le 0.001$, $****P \le 0.0001$ by Mann-Whitney Test.

**Supp. Fig. 9**

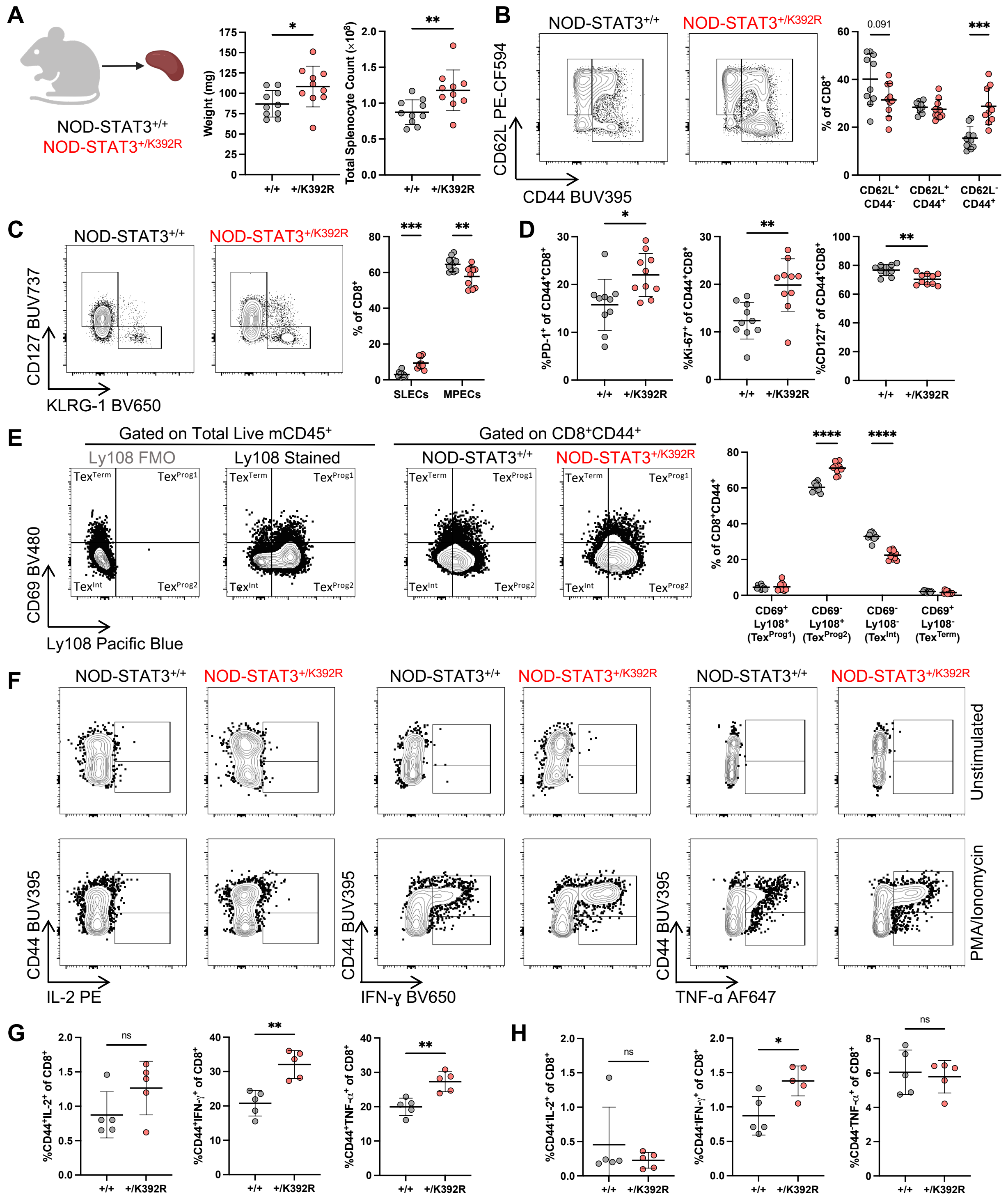
A
NOD-STAT3+/+
NOD-STAT3+/K392R
Weight (mg)
Total Splenocyte Count (×10⁸)
+/+
+/K392R
*
**
B
NOD-STAT3+/+
NOD-STAT3+/K392R
CD62L PE-CF594
CD44 BUV395
% of CD8+
0.091
***
CD62L+ CD44-
CD62L+ CD44+
CD62L- CD44+
C
NOD-STAT3+/+
NOD-STAT3+/K392R
CD127 BUV737
KLRG-1 BV650
% of CD8+
SLECs
MPECs
***
**
D
%PD-1+ of CD44+CD8+
%Ki-67+ of CD44+CD8+
%CD127+ of CD44+CD8+
+/+
+/K392R
E
Gated on Total Live mCD45+
Gated on CD8+CD44+
Ly108 FMO
Ly108 Stained
NOD-STAT3+/+
NOD-STAT3+/K392R
TexTerm
TexProg1
TexInt
TexProg2
CD69 BV480
Ly108 Pacific Blue
% of CD8+CD44+
****
CD69+ Ly108+ (TexProg1)
CD69- Ly108+ (TexProg2)
CD69- Ly108- (TexInt)
CD69+ Ly108- (TexTerm)
F
NOD-STAT3+/+
NOD-STAT3+/K392R
Unstimulated
PMA/Ionomycin
CD44 BUV395
IL-2 PE
IFN-γ BV650
TNF-α AF647
G
%CD44+IL-2+ of CD8+
%CD44+IFN-γ+ of CD8+
%CD44+TNF-α+ of CD8+
ns
H
%CD44-IL-2+ of CD8+
%CD44-IFN-γ+ of CD8+
%CD44-TNF-α+ of CD8+

**Supplemental Figure 9: Additional characterization of $CD8^+$ T cell compartment in NOD-$STAT3^{+/K392R}$ model. (A)** Comparison of spleen weight and total cell numbers in the spleen of NOD-$STAT3^{+/+}$ (n = 10) and NOD-$STAT3^{+/K392R}$ (n = 10) mice. **(B)** Representative $CD8^+$ T cell subsets and quantification in NOD-$STAT3^{+/+}$ and NOD-$STAT3^{+/K392R}$ mice. **(C)** Comparison of MPEC ($CD127^+KLRG\text{-}1^-$) and SLEC ($CD127^-KLRG\text{-}1^+$) populations in NOD-$STAT3^{+/+}$ and NOD-$STAT3^{+/K392R}$ mice. **(D)** Quantification of activation markers on $CD44^+CD8^+$ T cells in NOD-$STAT3^{+/+}$ and NOD-$STAT3^{+/K392R}$ mice. **(E)** Representative flow cytometry plots and quantification of $T_{Ex}$ progenitors on mouse $CD8^+$ T cells. Definitions: $CD69^+Ly108^+$ = $Tex^{Prog1}$, $CD69^-Ly108^+$ = $T_{Ex}{}^{Prog2}$, $CD69^-Ly108^-$ = $T_{Ex}{}^{Int}$, $CD69^+Ly108^-$ = $T_{Ex}{}^{Term}$. **(F)** Representative flow cytometry plots of cytokine production by $CD8^+$ T cells after 5 hours with PMA/Ionomycin. **(G)** Percent $cytokine^+CD44^+CD8^+$ or **(H)** $cytokine^+CD44^-CD8^+$ T cells (n = 5 each). Assay performed in total splenocytes and then gated on live $mCD45^+B220^-Gr\text{-}1^-CD3^+CD8^+$ cells. For **A-E**: data are pooled from 2 independent experiments. $^*P \leq 0.05$, $^{**}P \leq 0.01$, $^{***}P \leq 0.001$, $^{****}P \leq 0.0001$ by Mann-Whitney Test.

**Supp. Fig. 10**

A

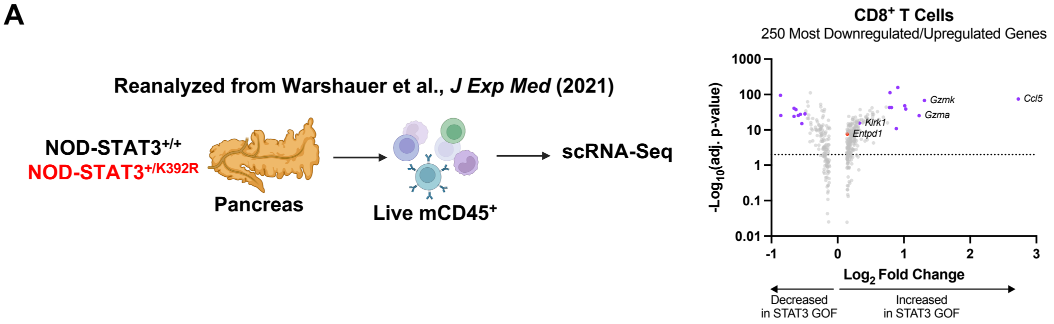


**Supplemental Figure 10: scRNA-Seq re-analysis of murine CD8$^{+}$ T cells for CD39 expression in NOD-STAT3$^{+/K392R}$ model. (A)** Schematic of testing strategy followed by volcano plot generated from scRNA-Seq re-analysis of *Entpd1* (CD39) expression on CD8$^{+}$ T cells within CD45$^{+}$ lymphocytes isolated from the islets of 8–10-week-old nondiabetic mice from NOD-STAT3$^{+/+}$ and NOD-STAT3$^{+/K392R}$ mice.

**Supp. Fig. 11**

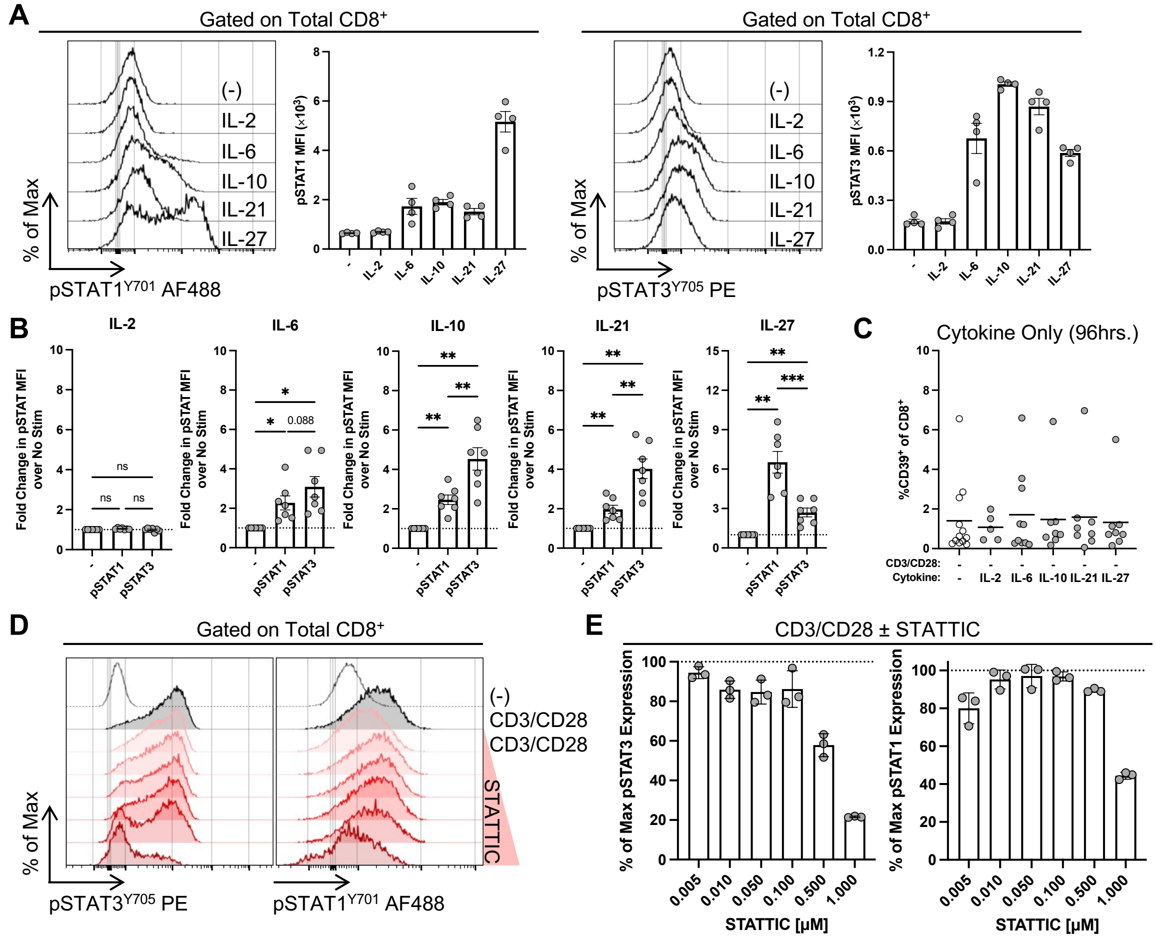


**Supplemental Figure 11: Characterization of STAT1 and STAT3 signaling in CD8$^+$ T cells by various cytokines. (A)** pSTAT1 and pSTAT3 induction in CD8$^+$ T cells after activation with indicated cytokines (IL-2 = 25IU/mL, IL-6 = 100ng/mL, and IL-10, IL-21 and IL-27 = 25ng/mL) for 20 minutes; data is representative of 1 of 2 experiments. **(B)** Summary of pSTAT3 and pSTAT1 fold change from 2 independent experiments (n = 7). **(C)** Frequencies of CD39$^+$CD8$^+$ T cells after 96-hour activation with indicated cytokines (n > 5). **(D)** Representative pSTAT3 and pSTAT1 levels in CD8$^+$ T cells incubated in the presence of αCD3/αCD28 ± STATTIC for 96hrs. **(E)** Quantification of percent of max pSTAT3 and pSTAT1 levels in αCD3/αCD28 + STATTIC treated cells relative to αCD3/αCD28 condition alone. For **D-E**: assays were performed in technical triplicates. *$P \leq 0.05$, **$P \leq 0.01$, ***$P \leq 0.001$ by RM one-way ANOVA with Tukey's multiple comparison test.

**Supp. Fig. 12**

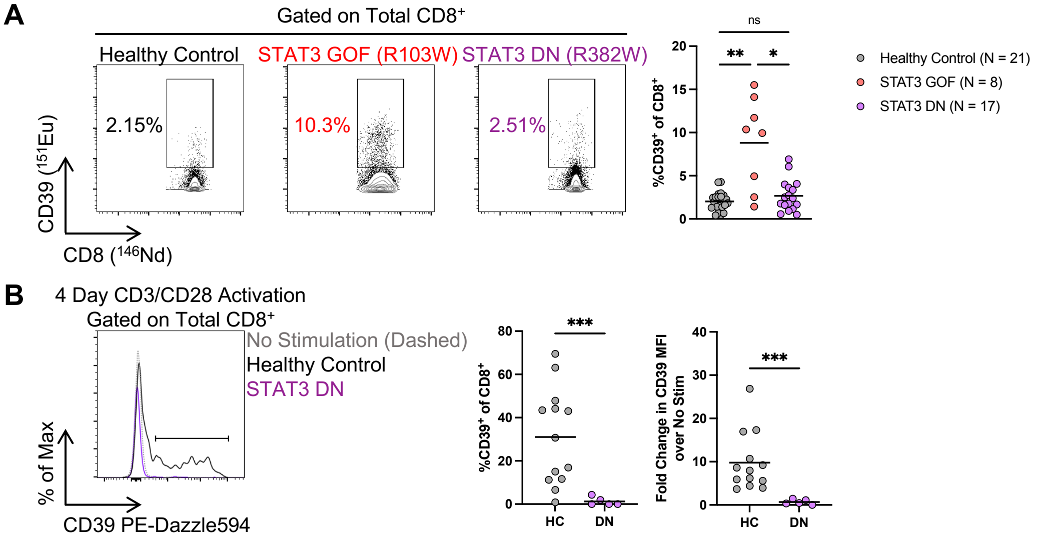


**Supplemental Figure 12: CD39 induction in patients bearing STAT3 DN variants. (A)** Representative mass cytometry CD39 frequency plots from a HC, patient with STAT3 GOF, and a patient with STAT3 DN, followed by quantification from multiple individuals. Healthy control and STAT3 GOF values are also used in **Supp. Fig. 5A**. **(B)** On the left, representative CD39 induction in a STAT3 DN patient compared to HC after 4 days of activation with αCD3/αCD28. On the right, quantification of CD39 frequency and fold change in expression from HCs (n = 13) and patients with STAT3 DN (n = 5). For **A**: data are pooled from 3 independent experiments. For **B**: data are from 1 experiment. *$P \leq 0.05$, **$P \leq 0.01$, ***$P \leq 0.001$ by Mann-Whitney Test or by Kruskal-Wallis test with Dunn's multiple comparison test.

**Supp. Fig. 13**

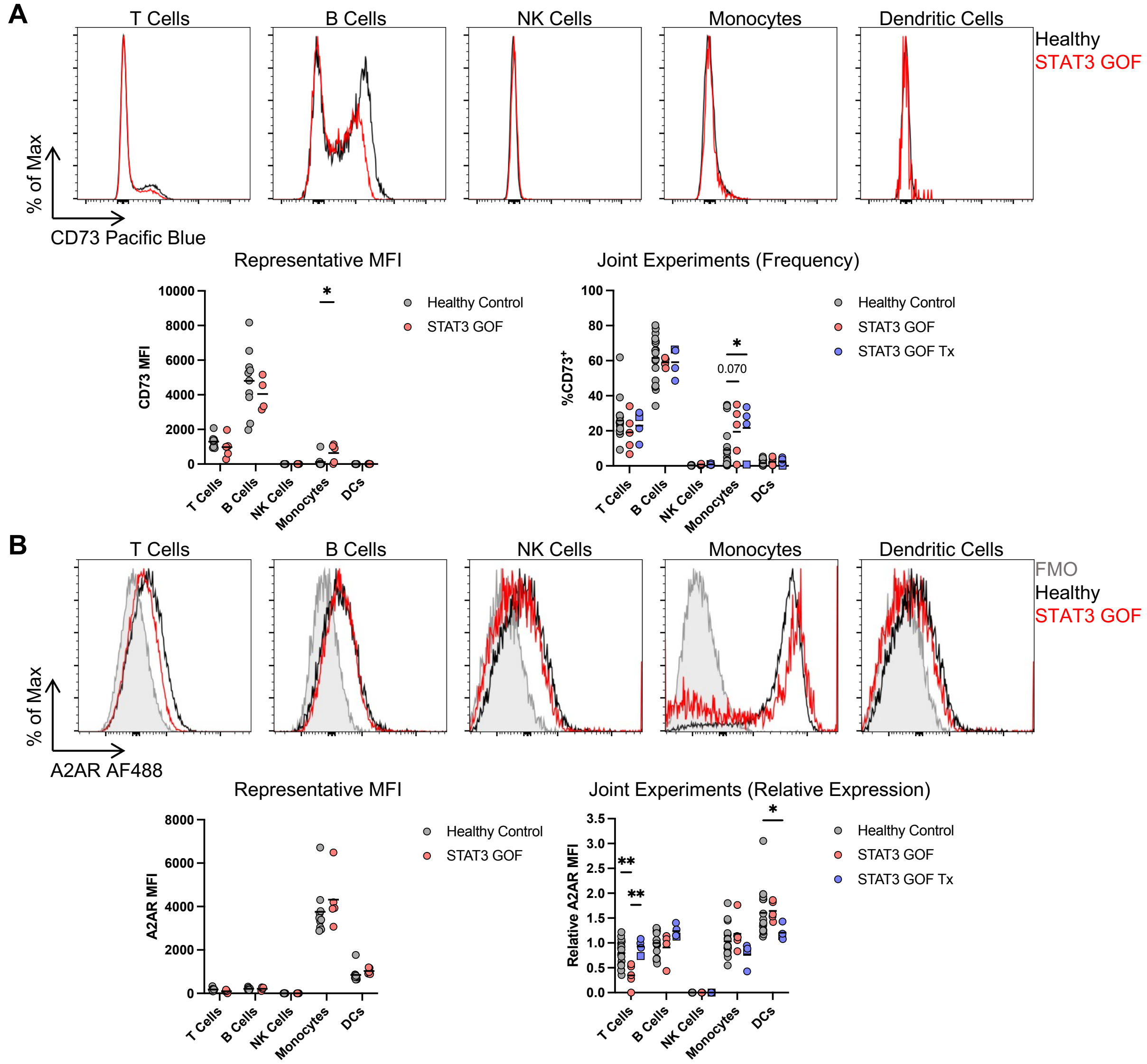


**Supplemental Figure 13: CD73 and $A_{2A}R$ across STAT3 GOF immune cell lineages by spectral flow cytometry**. **(A)** Representative CD73 histograms across immune cell lineages and quantification (below) of CD73 MFI and frequencies across cell types by disease state. A total of n = 18 HC (gray), n = 5 untreated (red), and n = 4 treated (blue) patients with STAT3 GOF were analyzed. Representative data from one experiment (left) and pooled (right) are shown. **(B)** Representative $A_{2A}R$ histograms across immune cell lineages and quantification (below) of $A_{2A}R$ expression across cell types by disease state after normalization. Representative data from one experiment (left) and pooled (right) are shown. Relative $A_{2A}R$ expression was calculated by comparing to the same well characterized donor used in **Supp. Fig. 6B** for a total of n =17 HC (gray), n = 5 untreated (red), and n = 4 treated (blue) patients with STAT3 GOF. Blue square represents patient treated with Rapamycin rather than JAKi (which is shown as blue circles). $*P \leq 0.05$, $**P \leq 0.01$ by Mann-Whitney test or two-way ANOVA with Tukey's multiple comparison test. Not listed or ns was not significant.

**Supp. Fig. 14**

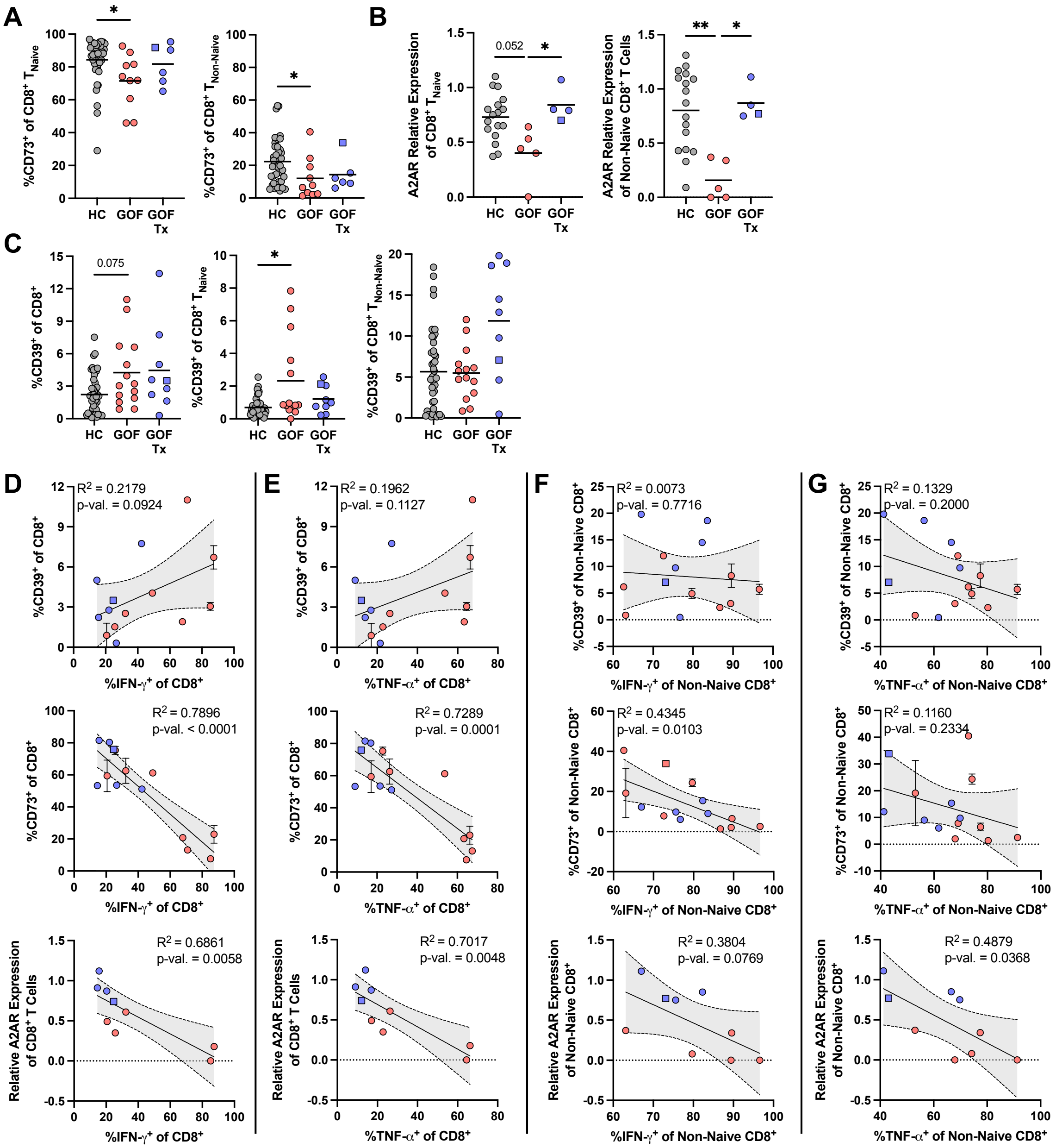
A
B
C
D
E
F
G
%CD73+ of CD8+ TNaive
%CD73+ of CD8+ TNon-Naive
A2AR Relative Expression of CD8+ TNaive
A2AR Relative Expression of Non-Naive CD8+ T Cells
%CD39+ of CD8+
%CD39+ of CD8+ TNaive
%CD39+ of CD8+ TNon-Naive
HC
GOF
GOF Tx
0.052
0.075
*
**
R2 = 0.2179
p-val. = 0.0924
R2 = 0.1962
p-val. = 0.1127
R2 = 0.0073
p-val. = 0.7716
R2 = 0.1329
p-val. = 0.2000
R2 = 0.7896
p-val. < 0.0001
R2 = 0.7289
p-val. = 0.0001
R2 = 0.4345
p-val. = 0.0103
R2 = 0.1160
p-val. = 0.2334
R2 = 0.6861
p-val. = 0.0058
R2 = 0.7017
p-val. = 0.0048
R2 = 0.3804
p-val. = 0.0769
R2 = 0.4879
p-val. = 0.0368
%IFN-γ+ of CD8+
%TNF-α+ of CD8+
%IFN-γ+ of Non-Naive CD8+
%TNF-α+ of Non-Naive CD8+
%CD39+ of Non-Naive CD8+
%CD73+ of CD8+
%CD73+ of Non-Naive CD8+
Relative A2AR Expression of CD8+ T Cells
Relative A2AR Expression of Non-Naive CD8+

**Supplemental Figure 14: Additional CD73 and $A_{2A}R$ expression and cytokine correlations by $CD8^+$ T cell subsets**. **(A)** CD73 frequencies within naive and non-naïve $CD8^+$ T cells in HC (gray, n = 42), untreated (red, n = 10), or treated (blue, n = 6) patients with STAT3 GOF. **(B)** Relative $A_{2A}R$ expression on naïve and non-naïve $CD8^+$ T cells in HC (n = 17), untreated (n = 5), or treated (n = 4) patients with STAT3 GOF. **(C)** Frequencies of CD39 positivity in total, naïve, and non-naïve $CD8^+$ T cells in HC (n = 47), untreated (n = 14), or treated (n = 9) patients with STAT3 GOF. $CD39^+$ values in total $CD8^+$ T cells from HC and STAT3 GOF were also used in **Fig. 3A**. Correlation between IFN-γ **(D)** and TNF-α **(E)** production and CD39, CD73 and $A_{2A}R$ levels in total $CD8^+$ T cells. Correlation between IFN-γ **(F)** and TNF-α **(G)** production and CD39, CD73 and $A_{2A}R$ levels in non-naïve $CD8^+$ T cells. Blue square represents patient treated with Rapamycin rather than JAKi (which is shown as blue circles). $*P \leq 0.05$, $**P \leq 0.01$ by Kruskal-Wallis test with Dunn's multiple comparison test or simple linear regression, as appropriate; shaded area indicates 95% confidence interval.

#### Supp. Fig. 15

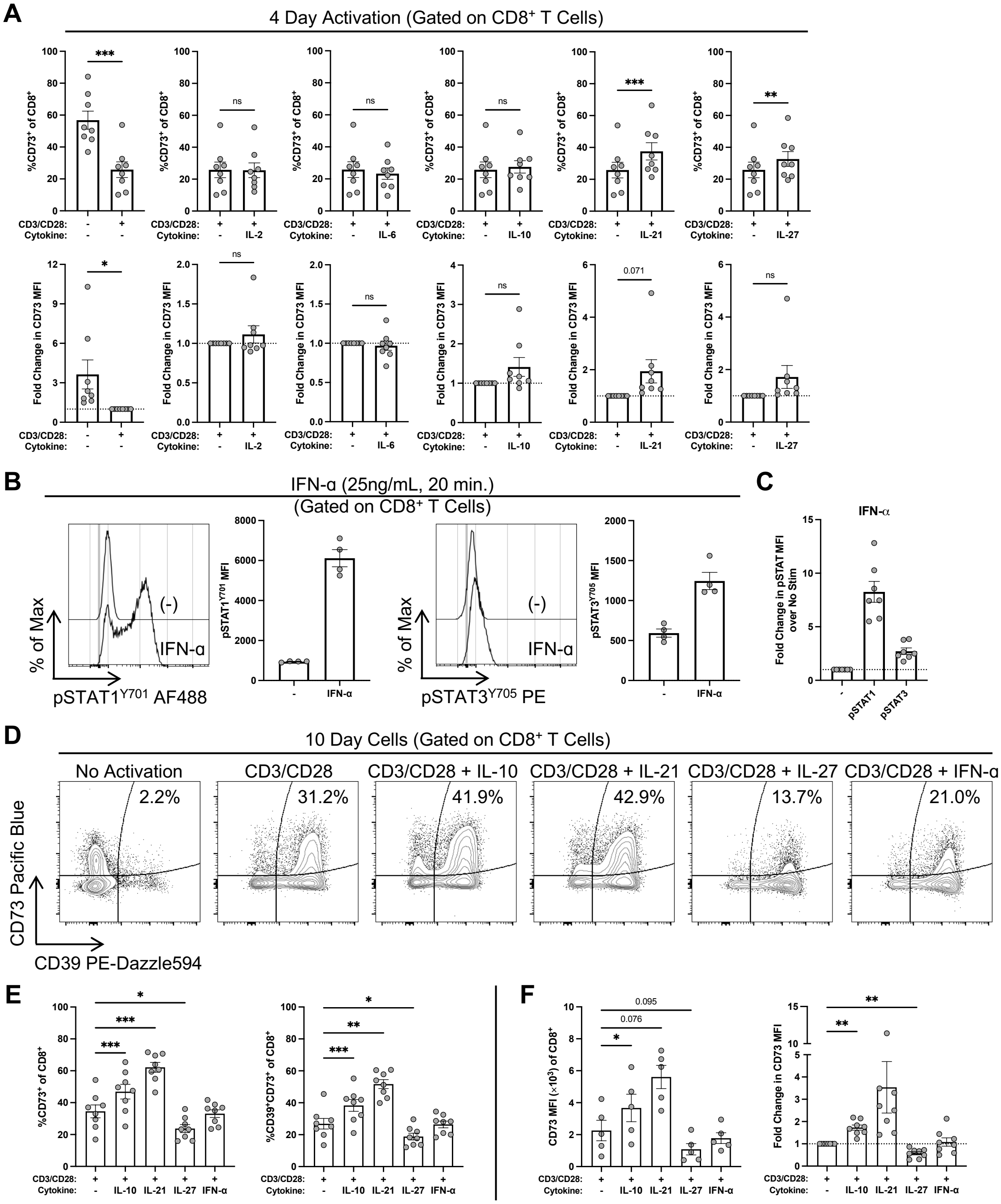
A
4 Day Activation (Gated on CD8⁺ T Cells)
%CD73⁺ of CD8⁺
Fold Change in CD73 MFI
CD3/CD28:
Cytokine:
IL-2
IL-6
IL-10
IL-21
IL-27
***
ns
**
*
0.071
B
IFN-α (25ng/mL, 20 min.)
(Gated on CD8⁺ T Cells)
% of Max
(-)
IFN-α
pSTAT1Y701 AF488
pSTAT1Y701 MFI
pSTAT3Y705 PE
pSTAT3Y705 MFI
C
IFN-α
Fold Change in pSTAT MFI over No Stim
pSTAT1
pSTAT3
D
10 Day Cells (Gated on CD8⁺ T Cells)
No Activation
CD3/CD28
CD3/CD28 + IL-10
CD3/CD28 + IL-21
CD3/CD28 + IL-27
CD3/CD28 + IFN-α
2.2%
31.2%
41.9%
42.9%
13.7%
21.0%
CD73 Pacific Blue
CD39 PE-Dazzle594
E
%CD73⁺ of CD8⁺
%CD39⁺CD73⁺ of CD8⁺
F
CD73 MFI (×10³) of CD8⁺
Fold Change in CD73 MFI
0.095
0.076

**Supplemental Figure 15: Role of STAT3 in regulating CD73 expression in $CD8^+$ T cells. (A)** Frequency of $CD73^+CD8^+$ T cells (top row) and fold change in CD73 MFI (bottom row, relative to αCD3/αCD28 condition) in HC PBMCs left unactivated or activated with αCD3/αCD28 ± indicated cytokine for 96 hours; data are pooled from 2 independent experiments (n = 8). **(B)** Induction of pSTAT1 (left) and pSTAT3 (right) in $CD8^+$ T cells after 20 minutes of activation with 25ng/mL IFN-α from 1 of 2 independent experiments. **(C)** Comparison of fold change in pSTAT1 and pSTAT3 levels pooled from 2 independent experiments (n = 7). **(D)** Representative bi-variate flow cytometry plots of CD39 and CD73 expression on $CD8^+$ T cells after 10 days of activation, with media exchange at Day 4. **(E)** Quantification of frequencies of $CD73^+$ and $CD39^+CD73^+$ $CD8^+$ T cells after 10 days of activation in the indicated conditions. **(F)** CD73 MFI (left, representative of one experiment) and fold change in CD73 MFI (pooled from 2 experiments) relative to αCD3/αCD28 condition after 10 days of activation. For **D-F**: n = 8. $^*P \leq 0.05$, $^{**}P \leq 0.01$, $^{***}P \leq 0.001$ by paired t-test or RM one-way ANOVA with Dunnett's multiple comparison test; adjusted p-values are listed.

**Supp. Fig. 16**

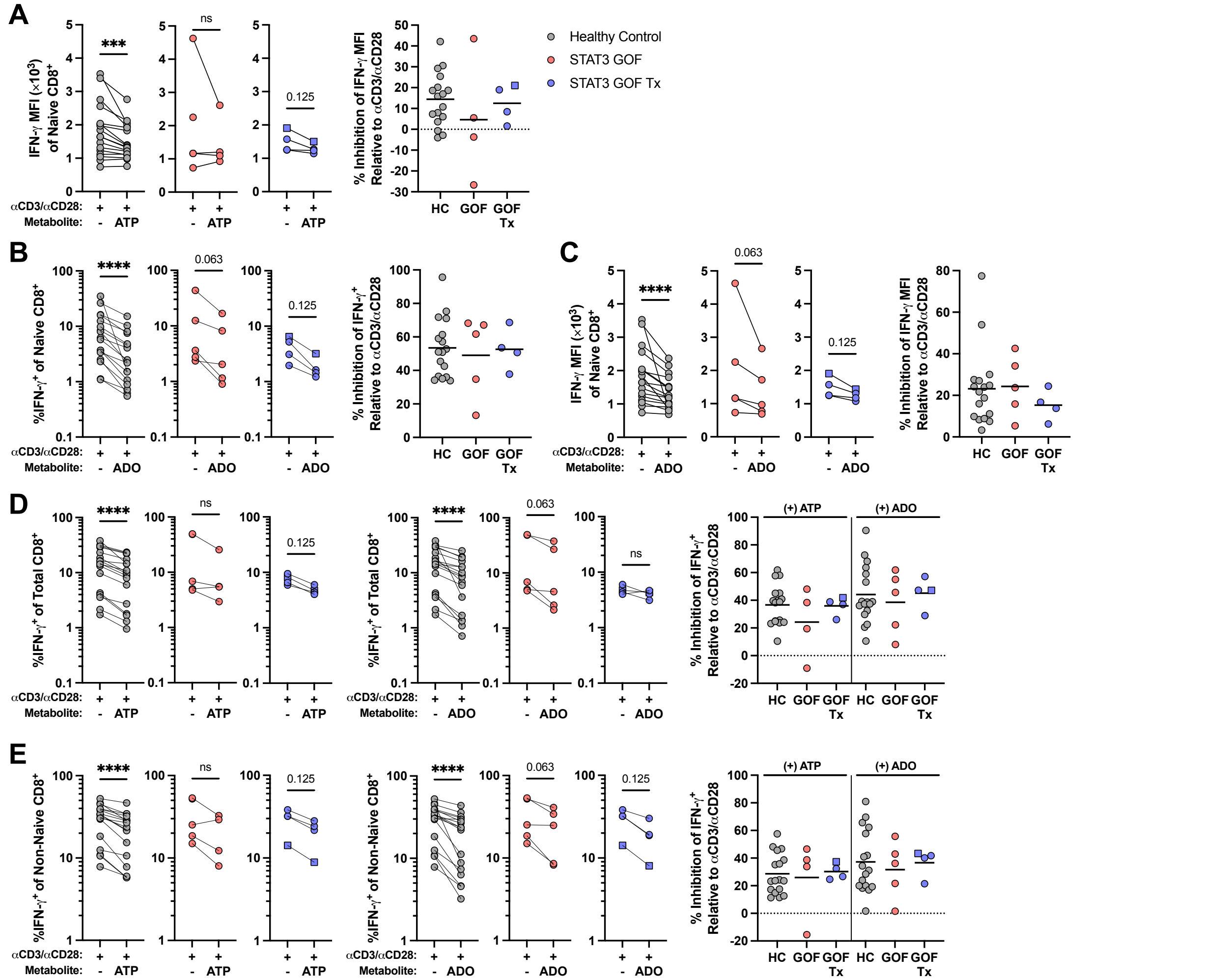


**Supplemental Figure 16: Additional quantification of cytokine production inhibition in total, naïve, and non-naïve CD8$^{+}$ T cells after activation in the presence of ATP or adenosine.** **(A)** Paired analysis of naïve CD8$^{+}$ T cell IFN-γ MFI after *in vitro* activation with αCD3/αCD28 in the presence or absence of ATP, followed by quantification of percent inhibition of IFN-γ MFI. Paired analysis of naïve CD8$^{+}$ T cell IFN-γ$^{+}$ frequency **(B)** and MFI **(C)** after activation with αCD3/αCD28 in the presence or absence of ADO, followed quantification of percent inhibition of IFN-γ frequency and MFI. Paired analysis of frequency of IFN-γ$^{+}$ cells in total **(D)** and non-naïve **(E)** CD8$^{+}$ T cells after activation with αCD3/αCD28 in the presence of ATP or ADO, followed quantification of percent inhibition of IFN-γ production. Data are pooled from 2 independent experiments with 3 repeat HC and 1 patient with STAT3 GOF run in each experiment; repeated samples across the two experiments were averaged into one data point per participant for a total of n = 17 HC (gray), n = 4 or 5 untreated (red), and n = 4 treated (blue) patients with STAT3 GOF. Blue square represents patient treated with Rapamycin rather than JAKi (which is shown as blue circles). ***$P \leq 0.001$, ****$P \leq 0.0001$ by Wilcoxon matched-pairs signed rank test.

**Table S1. Healthy control, STAT3 GOF, and STAT3 DN participant metadata.**

Table S1 is available as an attached supplemental spreadsheet (Excel) due to its size.

**Table S2. List of DEGs in conventional $CD8^+$ T cells from human scRNA-seq analysis.**

Table S2 is available as an attached supplemental spreadsheet (Excel) due to its size.

#### Table S3. GSEA result summary in conventional CD8$^{+}$ T cells.

| NAME | SIZE | ES | NES | NOM p-val | FDR q-val | FWER p-val | RANK AT MAX |
|---|---|---|---|---|---|---|---|
| **Upregulated in STAT3 GOF** | | | | | | | |
| HALLMARK_INTERFERON_GAMMA_RESPONSE | 173 | 0.51208454 | 2.1617355 | 0 | 0 | 0 | 3711 |
| HALLMARK_INTERFERON_ALPHA_RESPONSE | 93 | 0.5218289 | 2.0236654 | 0 | 0.000173 | 0.0004 | 1391 |
| HALLMARK_G2M_CHECKPOINT | 182 | 0.45364296 | 1.9362842 | 0 | 0.000311 | 0.001 | 3499 |
| HALLMARK_ALLOGRAFT_REJECTION | 160 | 0.39055002 | 1.6409074 | 0.0003196 | 0.017706 | 0.0762 | 1929 |
| HALLMARK_ANGIOGENESIS | 13 | 0.6443134 | 1.6300333 | 0.0146421 | 0.015815 | 0.0849 | 799 |
| HALLMARK_MITOTIC_SPINDLE | 183 | 0.37711966 | 1.6124222 | 0 | 0.016007 | 0.1024 | 5350 |
| HALLMARK_IL6_JAK_STAT3_SIGNALING | 54 | 0.44277263 | 1.5635253 | 0.0117019 | 0.023905 | 0.1711 | 1069 |
| HALLMARK_INFLAMMATORY_RESPONSE | 125 | 0.37017506 | 1.5037724 | 0.0046624 | 0.037382 | 0.2862 | 2545 |
| HALLMARK_E2F_TARGETS | 185 | 0.3257695 | 1.3908293 | 0.0125846 | 0.092994 | 0.6131 | 3499 |
| HALLMARK_APOPTOSIS | 134 | 0.30513975 | 1.2494717 | 0.0780119 | 0.266093 | 0.9527 | 3090 |
| HALLMARK_TGF_BETA_SIGNALING | 48 | 0.36092505 | 1.2395236 | 0.1402816 | 0.260194 | 0.9613 | 3825 |
| HALLMARK_PI3K_AKT_MTOR_SIGNALING | 93 | 0.31946695 | 1.2382655 | 0.1093492 | 0.240469 | 0.9625 | 6664 |
| HALLMARK_PROTEIN_SECRETION | 90 | 0.32169664 | 1.2374315 | 0.1171448 | 0.22347 | 0.9635 | 5597 |
| HALLMARK_ESTROGEN_RESPONSE_LATE | 119 | 0.3056772 | 1.2232299 | 0.1065343 | 0.23038 | 0.975 | 2167 |
| HALLMARK_TNFA_SIGNALING_VIA_NFKB | 164 | 0.2576583 | 1.0837178 | 0.2756308 | 0.530647 | 1 | 2204 |
| HALLMARK_P53_PATHWAY | 166 | 0.25319943 | 1.0669346 | 0.3049363 | 0.545587 | 1 | 2539 |
| HALLMARK_ESTROGEN_RESPONSE_EARLY | 127 | 0.2555726 | 1.0390085 | 0.3664861 | 0.595937 | 1 | 2167 |
| HALLMARK_IL2_STAT5_SIGNALING | 161 | 0.24369316 | 1.0230992 | 0.3948685 | 0.610352 | 1 | 2791 |
| HALLMARK_COMPLEMENT | 143 | 0.22850192 | 0.9450612 | 0.5934933 | 0.820208 | 1 | 2943 |
| HALLMARK_SPERMATOGENESIS | 72 | 0.24838637 | 0.9205552 | 0.6161081 | 0.856555 | 1 | 2950 |
| HALLMARK_UV_RESPONSE_DN | 99 | 0.22630708 | 0.88524216 | 0.7169439 | 0.919609 | 1 | 5028 |
| HALLMARK_ANDROGEN_RESPONSE | 81 | 0.2192046 | 0.8308656 | 0.8089907 | 1 | 1 | 1698 |
| HALLMARK_HEME_METABOLISM | 148 | 0.19880524 | 0.82817465 | 0.8713687 | 0.977027 | 1 | 4780 |
| HALLMARK_WNT_BETA_CATENIN_SIGNALING | 35 | 0.24611123 | 0.79375875 | 0.8039423 | 0.997934 | 1 | 1812 |
| HALLMARK_HEDGEHOG_SIGNALING | 20 | 0.2634278 | 0.74397695 | 0.8307227 | 1 | 1 | 4265 |
| HALLMARK_UNFOLDED_PROTEIN_RESPONSE | 106 | 0.12970278 | 0.5126856 | 1 | 1 | 1 | 4340 |
| HALLMARK_OXIDATIVE_PHOSPHORYLATION | 198 | 0.08511553 | 0.36575946 | 1 | 0.999987 | 1 | 13541 |
| **Downregulated in STAT3 GOF** | | | | | | | |
| **NAME** | **SIZE** | **ES** | **NES** | **NOM p-val** | **FDR q-val** | **FWER p-val** | **RANK AT MAX** |
| HALLMARK_PANCREAS_BETA_CELLS | 13 | -0.63277864 | -1.6534991 | 0.0159539 | 0.074123 | 0.061 | 683 |
| HALLMARK_MYC_TARGETS_V2 | 57 | -0.43097544 | -1.6006259 | 0.0066178 | 0.058798 | 0.0949 | 3674 |
| HALLMARK_GLYCOLYSIS | 146 | -0.33231044 | -1.4458131 | 0.0072633 | 0.153016 | 0.3263 | 3307 |
| HALLMARK_APICAL_SURFACE | 31 | -0.44156492 | -1.4351146 | 0.0510133 | 0.124624 | 0.3482 | 2240 |
| HALLMARK_KRAS_SIGNALING_DN | 81 | -0.3524503 | -1.4005957 | 0.0301445 | 0.131989 | 0.4349 | 1738 |
| HALLMARK_UV_RESPONSE_UP | 126 | -0.30240548 | -1.2866288 | 0.0525353 | 0.270103 | 0.7606 | 3896 |
| HALLMARK_MYOGENESIS | 112 | -0.29367712 | -1.2319508 | 0.0993727 | 0.347931 | 0.8834 | 2806 |

| | | | | | | | |
|---|---|---|---|---|---|---|---|
| HALLMARK_EPITHELIAL_MESENCHYMAL_TRANSITION | 91 | -0.30347574 | -1.2303358 | 0.100785 | 0.307982 | 0.8858 | 1830 |
| HALLMARK_BILE_ACID_METABOLISM | 74 | -0.31498793 | -1.2269938 | 0.1236094 | 0.280457 | 0.8916 | 2935 |
| HALLMARK_ADIPOGENESIS | 163 | -0.2730537 | -1.2092015 | 0.0870483 | 0.286252 | 0.9206 | 2417 |
| HALLMARK_HYPOXIA | 142 | -0.26434535 | -1.1402669 | 0.1739694 | 0.414726 | 0.9846 | 2963 |
| HALLMARK_CHOLESTEROL_HOMEOSTASIS | 63 | -0.28694054 | -1.0858455 | 0.3015051 | 0.532358 | 0.9983 | 1369 |
| HALLMARK_XENOBIOTIC_METABOLISM | 121 | -0.25642326 | -1.0829833 | 0.2683756 | 0.499942 | 0.9985 | 2798 |
| HALLMARK_APICAL_JUNCTION | 125 | -0.25395516 | -1.0806516 | 0.2744593 | 0.470623 | 0.9986 | 2240 |
| HALLMARK_COAGULATION | 63 | -0.26736543 | -1.0126895 | 0.4242351 | 0.64403 | 1 | 2085 |
| HALLMARK_MTORC1_SIGNALING | 188 | -0.22053681 | -0.995604 | 0.4602917 | 0.661111 | 1 | 813 |
| HALLMARK_KRAS_SIGNALING_UP | 117 | -0.23425283 | -0.9905183 | 0.4742991 | 0.638213 | 1 | 1612 |
| HALLMARK_FATTY_ACID_METABOLISM | 123 | -0.21259327 | -0.9017264 | 0.7069455 | 0.883654 | 1 | 3506 |
| HALLMARK_PEROXISOME | 84 | -0.22022115 | -0.87832516 | 0.7291102 | 0.903865 | 1 | 2404 |
| HALLMARK_NOTCH_SIGNALING | 22 | -0.2841464 | -0.85108507 | 0.6867968 | 0.925591 | 1 | 606 |
| HALLMARK_REACTIVE_OXYGEN_SPECIES_PATHWAY | 46 | -0.21190755 | -0.7510673 | 0.888783 | 1 | 1 | 3535 |
| HALLMARK_MYC_TARGETS_V1 | 195 | -0.1649641 | -0.74635416 | 0.9906542 | 0.989005 | 1 | 11367 |
| HALLMARK_DNA_REPAIR | 144 | -0.13774304 | -0.5970895 | 1 | 0.993725 | 1 | 4394 |

#### Table S4. Mass cytometry antibodies.

| Channel/ Metal | Marker | Clone | Dilution | Surface / ICS | Manufacturer | Catalog # | Purchased/ Conjugated | Conjugation Kit Manufacturer | Conjugation Kit Catalog # |
|---|---|---|---|---|---|---|---|---|---|
| 89Y | CD45 | HI30 | 300 | Surface | Fluidigm | 3089003B | Purchased | | |
| 113 In | CD45RO | UCHL1 | 100 | Surface | BD Biosciences | 555491 | Conjugated | Metal from Penn CyTOF Core | |
| 115 In | CD123 | 6H6 | 50 | Surface | BioLegend | 306002 | Conjugated | Metal from Penn CyTOF Core | |
| 139 La | Live/Dead | N/A | 300 | N/A | Macrocyclics | B-272-50 | Conjugated | Sigma Aldrich | 203521-25g |
| 141 Pr | CD3 | UCHT1 | 800 | ICS | BioLegend | 300443 | Conjugated | Fluidigm | 201141A |
| 142 Nd | CD26 | BA 5b | 200 | Surface | BioLegend | 302702 | Conjugated | Fluidigm | 201142A |
| 143 Nd | CD4 | RPA-T4 | 800 | ICS | BioLegend | 300502 | Conjugated | Fluidigm | 201143A |
| 144 Nd | CD11b | ICRF44 | 150 | Surface | Fluidigm | 3144001B | Purchased | | |
| 145 Nd | CD19 | HIB19 | 100 | Surface | BioLegend | 302202 | Conjugated | Fluidigm | 201145A |
| 146 Nd | CD8 | RPA-T8 | 100 | ICS | Fluidigm | 3146001B | Purchased | | |
| 147 Sm | CD14 | M5E2 | 200 | Surface | BioLegend | 301843 | Conjugated | Fluidigm | 201147A |
| 148 Nd | CD56 | HCD56 | 150 | Surface | BioLegend | 318345 | Conjugated | Fluidigm | 201148A |
| 149 Sm | CD11c | 3.9 | 200 | Surface | CyTEK | 70-0116-U100 | Conjugated | Fluidigm | 201149A |
| 150 Nd | FceRI | AER-37 (CRA-1) | 150 | Surface | Fluidigm | 3150027B | Purchased | | |
| 151 Eu | CD39 | A1 | 400 | Surface | BioLegend | 328202 | Conjugated | Fluidigm | 201151A |
| 152 Sm | Granzyme B | CLB-GB11 | 800 | ICS | Invitrogen | MA110338 | Conjugated | Fluidigm | 201152A |
| 153 Eu | CD45RA | HI100 | 150 | Surface | Fluidigm | 3153001B | Purchased | | |
| 154 Sm | NKp46 | 9E2 | 100 | Surface | BioLegend | 331947 | Conjugated | Fluidigm | 201154A |
| 155 Gd | CD27 | L128 | 200 | Surface | Fluidigm | 3155001B | Purchased | | |
| 156 Gd | Helios | 22F6 | 100 | ICS | BioLegend | 137202 | Conjugated | Fluidigm | 201156A |
| 158 Gd | PD-1 | EH12.2H7 | 400 | Surface | BioLegend | 329902 | Conjugated | Fluidigm | 201158A |
| 159 Tb | CCR7 | G043H7 | 200 | Surface | Fluidigm | 3159003A | Purchased | | |
| 160 Gd | Tbet | 4B10 | 200 | ICS | Fluidigm | 3160010B | Purchased | | |
| 161 Dy | CTLA-4 | BNI3 | 100 | ICS | Fluidigm | 3161004B | Purchased | | |
| 162 Dy | Foxp3 | PCH101 | 100 | ICS | Fluidigm | 3162011A | Purchased | | |
| 163 Dy | CRTH2 | BM16 | 75 | Surface | Fluidigm | 3163003B | Purchased | | |
| 164 Dy | CD161 | HP-3G10 | 200 | Surface | Fluidigm | 3164009B | Purchased | | |
| 165 Ho | Eomes | WD1928 | 300 | ICS | Invitrogen | 14-4877-82 | Conjugated | Fluidigm | 201165A |
| 166 Er | TCF-1 | 7F11A10 | 50 | ICS | BioLegend | 655202 | Conjugated | Fluidigm | 201166A |
| 167 Er | CD38 | HIT2 | 200 | Surface | Fluidigm | 3167001B | Purchased | | |
| 168 Er | CD138 | DL-101 | 100 | Surface | Fluidigm | 3168009B | Purchased | | |
| 169 Tm | TIGIT | MBSA43 | 100 | Surface | Invitrogen | 16-9500-82 | Purchased | | |
| 170 Er | CXCR5 | RF8B2 | 50 | Surface | BD Biosciences | 552032 | Conjugated | Fluidigm | 201170A |
| 171 Yb | ST2 | B4E6 | 50 | Surface | MDBioSciences | 101002 | Conjugated | Fluidigm | 201171A |
| 172 Yb | Ki67 | B56 | 400 | ICS | Fluidigm | 3172024B | Purchased | | |

| | | | | | | | | | |
|---|---|---|---|---|---|---|---|---|---|
| 173 Yb | HLA-DR | L243 | 300 | Surface | Fluidigm | 3173005B | Purchased | | |
| 174 Yb | TCRgd | B1 | 150 | Surface | BioLegend | 331202 | Conjugated | Fluidigm | 201174A |
| 175 Lu | IgD | IA6-2 | 800 | Surface | BioLegend | 348235 | Conjugated | Fluidigm | 201175A |
| 176 Yb | CD127/ IL-7Ra | A019D5 | 200 | Surface | Fluidigm | 3176004B | Purchased | | |
| 191+193 | Iridium | N/A | 4000 | N/A | Fluidigm | 201192B | Purchased | | |
| 209 Bi | CD16 | 3G8 | 200 | Surface | Fluidigm | 3209002B | Purchased | | |

#### Table S5. Flow cytometry antibodies.

| PMA/Ionomycin Cytokine Panel (Human PBMCs) | | | | | | |
|---|---|---|---|---|---|---|
| **Marker** | **Fluorophore** | **Source** | **Clone** | **Catalog No.** | **Dilution** | **Stain** |
| Fc Block | N/A | BD Biosciences | N/A | 564220 | 200 | Surface |
| CD4 | BUV395 | BD Biosciences | RPA-T4 | 564724 | 400/400 | Surface/Intracellular |
| CD45RO | BUV805 | BD Biosciences | UCHL1 | 748367 | 200 | Surface |
| IL-2 | BV421 | BD Biosciences | 5344.111 | 562914 | 100 | Intracellular |
| CD14 | V500 | BD Biosciences | M5E2 | 561391 | 300 | Surface |
| CD16 | V500 | BD Biosciences | 3G8 | 561394 | 300 | Surface |
| CD19 | V500 | BD Biosciences | HIB19 | 561121 | 300 | Surface |
| Live/Dead | Aqua | Invitrogen/ThermoFisher | N/A | L34996A | 300 | Surface |
| CD8a | BV605 | BioLegend | RPA-T8 | 301040 | 100 | Surface |
| CD45RA | BV650 | BD Biosciences | HI100 | 563963 | 200 | Surface |
| CD27 | BV785 | BioLegend | O323 | 302832 | 200 | Surface |
| IL-17A | AF488 | BioLegend | BL168 | 512308 | 100 | Intracellular |
| TNF-α | PerCP-Cy5.5 | BioLegend | MAb11 | 502926 | 100 | Intracellular |
| IL-13 | PE | BioLegend | JES10-5A2 | 501903 | 100 | Intracellular |
| CD39 | PE-Dazzle 594 | BioLegend | A1 | 328224 | 200 | Surface |
| IFN-γ | PE-Cy7 | BioLegend | B27 | 506518 | 400 | Intracellular |
| IL-21 | AF647 | BioLegend | 3A3-N2 | 513006 | 50 | Intracellular |
| CD3 | APC-R700 | BD Biosciences | UCHT1 | 565119 | 400/400 | Surface/Intracellular |

| Purinergic Molecules Panel | | | | | | |
|---|---|---|---|---|---|---|
| **Marker** | **Fluorophore** | **Vendor** | **Clone** | **Catalog No.** | **Dilution** | **Stain** |
| Fc Block | N/A | BD Biosciences | N/A | 564220 | 200 | Surface |
| CD45RA | BUV395 | BD Biosciences | H100 | 740298 | 200 | Surface |
| Live/Dead | Blue | Invitrogen/ThermoFisher | N/A | L34962A | 1000 | Surface |
| CD16 | BUV496 | BD Biosciences | 3G8 | 612944 | 200 | Surface |
| CD14 | BUV563 | BD Biosciences | M5E2 | 741360 | 200 | Surface |
| CD45 | BUV615 | BD Biosciences | HI30 | 751472 | 300 | Surface |
| CD11c | BUV661 | BD Biosciences | B-ly6 | 612967 | 100 | Surface |
| CD56 | BUV737 | BD Biosciences | NCAM16.2 | 612766 | 200 | Surface |
| CD26 | BUV805 | BD Biosciences | L272 | 749179 | 100 | Surface |
| CD19 | BV421 | BD Biosciences | HIB19 | 562440 | 300 | Surface |
| CD73 | Pacific Blue | BioLegend | AD2 | 344012 | 100 | Surface |
| CD3 | BV570 | BioLegend | UCHT1 | 300436 | 400 | Surface |
| CD8a | BV785 | BioLegend | RPA-T8 | 301046 | 200 | Surface |
| A2AR | AF488 | R&D Systems | 599717 | FAB94971G | 50 | Surface |

| | | | | | | |
|---|---|---|---|---|---|---|
| CD20 | PerCP | BioLegend | 2H7 | 302324 | 50 | Surface |
| TCRgd | PerCP-eFluor 710 | Invitrogen/ThermoFisher | B1.1 | 46-9959-42 | 200 | Surface |
| CD4 | cFluor YG584 | CyTEK | SK3 | R7-20041 | 400 | Surface |
| CD25 | PE-Cy5 | BD Biosciences | M-A251 | 555433 | 200 | Surface |
| CD39 | PE-Fire 810 | BioLegend | A1 | 328245 | 200 | Surface |
| CD27 | APC | BioLegend | O323 | 302810 | 100 | Surface |
| CD127 | APC-R700 | BD Biosciences | HIL-7R-M21 | 565185 | 100 | Surface |
| CD38 | APC-eFluor 780 | Invitrogen/ThermoFisher | HIT2 | 47-0389-42 | 100 | Surface |
| HLA-DR | APC-Fire 810 | BioLegend | L243 | 307674 | 200 | Surface |

Control

| | | | | | | |
|---|---|---|---|---|---|---|
| IgG2A Isotype | AF488 | R&D Systems | 20102 | IC003G | 50 | Surface |

**T Cell Activation-Exhaustion Panel (Small)**

| **Marker** | **Fluorophore** | **Vendor** | **Clone** | **Catalog No.** | **Dilution** | **Stain** |
|---|---|---|---|---|---|---|
| Fc Block | N/A | BD Biosciences | N/A | 564220 | 200 | Surface |
| TIM-3 | BV421 | BioLegend | F38-2E2 | 345008 | 100 | Surface |
| CD69 | Pacific Blue | BioLegend | FN50 | 310920 | 200 | Surface |
| CD14 | V500 | BD Biosciences | M5E2 | 561391 | 300 | Surface |
| CD16 | V500 | BD Biosciences | 3G8 | 561394 | 300 | Surface |
| CD19 | V500 | BD Biosciences | HIB19 | 561121 | 300 | Surface |
| Live/Dead | Aqua | Invitrogen/ThermoFisher | N/A | L34996A | 300 | Surface |
| CD8a | BV605 | BioLegend | RPA-T8 | 301040 | 200 | Surface |
| CD45RA | BV650 | BD Biosciences | H100 | 563963 | 200 | Surface |
| CD27 | BV785 | BioLegend | O323 | 302832 | 200 | Surface |
| CD4 | AF488 | BD Biosciences | RPA-T4 | 557695 | 200 | Surface |
| TOX | PE | Miltenyi Biotec | REA473 | 130-120-716 | 200 | Intracellular |
| CD39 | PE-Dazzle 594 | BioLegend | A1 | 328224 | 200 | Surface |
| CD25 | PE-Cy5 | BD Biosciences | M-A251 | 555433 | 200 | Surface |
| LAG-3 | PE-Cy7 | BioLegend | 11C3C65 | 369310 | 200 | Surface |
| TCF1/TCF7 | AF647 | Cell Signaling Technologies | C63D9 | 6709S | 200 | Intracellular |
| CD3 | APC-R700 | BD Biosciences | UCHT1 | 565119 | 200 | Surface |
| PD-1 | APC-Fire 750 | BioLegend | EH12.2H7 | 329954 | 100 | Surface |

Additional Markers and Alternatives for Small T Cell Activation-Exhaustion Panel

| | | | | | | |
|---|---|---|---|---|---|---|
| CD4 | BUV395 | BD Biosciences | RPA-T4 | 564724 | 200 | Surface |
| CD45RO | BUV805 | BD Biosciences | UCHL1 | 748367 | 200 | Surface |
| CD57 | eFluor 450 | Invitrogen/ThermoFisher | TB01 | 48-0577-42 | 200 | Surface |
| CD73 | Pacific Blue | BioLegend | AD2 | 344012 | 100 | Surface |
| CD25 | FITC | BioLegend | M-A251 | 356106 | 200 | Surface |
| CD73 | FITC | BioLegend | AD2 | 344016 | 100 | Surface |

| | | | | | | |
|---|---|---|---|---|---|---|
| CTLA-4 | PE-Cy5 | BD Biosciences | BNI3 | 555854 | 50 | Intracellular |

| T Cell Activation-Exhaustion Panel (Large) | | | | | | |
|---|---|---|---|---|---|---|
| **Marker** | **Fluorophore** | **Vendor** | **Clone** | **Catalog No.** | **Dilution** | **Stain** |
| Fc Block | N/A | BD Biosciences | N/A | 564220 | 200 | Surface |
| Ki67 | BUV395 | BD Biosciences | B56 | 564071 | 300 | Intracellular |
| Live/Dead | Live/Dead Blue | Invitrogen/ThermoFisher | N/A | L34962A | 1000 | Surface |
| CD4 | BUV496 | BD Biosciences | RPA-T4 | 741134 | 400/400 | Surface/Intracellular |
| CD127 | BUV563 | BD Biosciences | HIL-7R-M21 | 748489 | 100 | Surface |
| CD27 | BUV737 | BD Biosciences | O323 | 751681 | 400 | Surface |
| CD49D | BUV805 | BD Biosciences | 9F10 | 749454 | 300 | Surface |
| PD-1 | BV421 | BioLegend | EH12.2H7 | 329920 | 100 | Surface |
| CD57 | eFluor 450 | Invitrogen/ThermoFisher | TB01 | 48-0577-42 | 200 | Surface |
| CD69 | BV480 | BD Biosciences | FN50 | 747519 | 200 | Surface |
| Granzyme B | BV510 | BD Biosciences | GB11 | 563388 | 300 | Intracellular |
| HLA-DR | BV570 | BioLegend | L243 | 307638 | 150 | Surface |
| CD8a | BV605 | BioLegend | RPA-T8 | 301040 | 100 | Surface |
| CD45RA | BV650 | BD Biosciences | H100 | 563963 | 200 | Surface |
| LAG-3 | BV711 | BioLegend | 11C3C65 | 369320 | 100 | Surface |
| CCR7 | BV750 | BioLegend | G043H7 | 353254 | 100 | Surface |
| Tbet | BV785 | BioLegend | 4B10 | 644835 | 300 | Intracellular |
| CD73 | FITC | BioLegend | AD2 | 344016 | 100 | Surface |
| KLRG1 | PerCP-Cy5.5 | BioLegend | SA231A2 | 367708 | 100 | Surface |
| CD38 | PerCP-eFluor 710 | Invitrogen/ThermoFisher | HIT2 | 46-0389-42 | 150 | Surface |
| TOX | PE | Miltenyi Biotec | REA473 | 130-120-716 | 200 | Intracellular |
| CD39 | PE-Dazzle 594 | BioLegend | A1 | 328224 | 200 | Surface |
| CTLA-4 | PE-Cy5 | BD Biosciences | BNI3 | 555854 | 200 | Intracellular |
| NKG2D | PE-Cy7 | BioLegend | 1D11 | 320812 | 200 | Surface |
| TCF1/TCF7 | AF647 | Cell Signaling Technologies | C63D9 | 6709S | 200 | Intracellular |
| CD3 | APC-R700 | BD Biosciences | UCHT1 | 565119 | 200 | Surface |
| EOMES | APC-eFluor 780 | Invitrogen/ThermoFisher | WD1928 | 47-4877-42 | 100 | Intracellular |

Additional Markers and Alternatives for T Cell Activation-Exhaustion Panel (Large)

| | | | | | | |
|---|---|---|---|---|---|---|
| Helios | PE-Cy7 | Invitrogen/ThermoFisher | 22F6 | 25-9883-42 | 200 | Intracellular |

| Single Cell RNA-Sequencing Panel for Sorting | | | | | | |
|---|---|---|---|---|---|---|
| **Marker** | **Fluorophore** | **Vendor** | **Clone** | **Catalog No.** | **Dilution** | **Stain** |
| CD14 | V500 | BD Biosciences | M5E2 | 561391 | 300 | Surface |
| CD16 | V500 | BD Biosciences | 3G8 | 561394 | 300 | Surface |
| CD19 | V500 | BD Biosciences | HIB19 | 561121 | 300 | Surface |
| Live/Dead | Aqua | Invitrogen/ThermoFisher | N/A | L34996A | 300 | Surface |

| | | | | | | |
|---|---|---|---|---|---|---|
| CD8a | BV605 | BioLegend | RPA-T8 | 301040 | 200 | Surface |
| CD4 | AF488 | BD Biosciences | RPA-T4 | 557695 | 200 | Surface |
| CD3 | APC-R700 | BD Biosciences | UCHT1 | 565119 | 100 | Surface |

| Phospho-Flow Panel Antibodies | | | | | | |
|---|---|---|---|---|---|---|
| **Marker** | **Fluorophore** | **Vendor** | **Clone** | **Catalog No.** | **Dilution** | **Stain** |
| Fc Block | N/A | BD Biosciences | N/A | 564220 | 200 | Surface |
| CD4 | BUV395 | BD Biosciences | RPA-T4 | 564724 | 200 | Surface |
| CD45RO | BUV805 | BD Biosciences | UCHL1 | 748367 | 200 | Surface |
| pSTAT6 (Y641) | V450 | BD Biosciences | 18/P-Stat6 | 561203 | 50 | Intracellular |
| CD14 | V500 | BD Biosciences | M5E2 | 561391 | 300 | Surface |
| CD16 | V500 | BD Biosciences | 3G8 | 561394 | 300 | Surface |
| CD19 | V500 | BD Biosciences | HIB19 | 561121 | 300 | Surface |
| Live/Dead | Aqua | Invitrogen/ThermoFisher | N/A | L34996A | 300 | Surface |
| CD8a | BV605 | BioLegend | RPA-T8 | 301040 | 200 | Surface |
| CD45RA | BV650 | BD Biosciences | H100 | 563963 | 200 | Surface |
| CD27 | BV785 | BioLegend | O323 | 302832 | 200 | Surface |
| pSTAT1 (Y701) | AF488 | BD Biosciences | 4a | 612596 | 50 | Intracellular |
| pSTAT5 (Y694) | RB705 | BD Biosciences | 47/Stat5(pY694) | 570286 | 100 | Intracellular |
| pSTAT3 (Y705) | PE | BD Biosciences | 4/P-STAT3 | 612569 | 50 | Intracellular |
| CD39 | PE-Cy7 | BioLegend | A1 | 328212 | 200 | Surface |
| Total STAT3 | APC | BD Biosciences | M59-50 | 560392 | 50 | Intracellular |
| CD3 | APC-R700 | BD Biosciences | UCHT1 | 565119 | 200 | Surface |

Additional Markers and Alternatives

| | | | | | | |
|---|---|---|---|---|---|---|
| pSTAT1 (Y701) | BV421 | BD Biosciences | 4a | 562985 | 50 | Intracellular |
| CD73 | FITC | BioLegend | AD2 | 344016 | 100 | Surface |

| Other Commonly Used Human Antibodies | | | | | | |
|---|---|---|---|---|---|---|
| **Marker** | **Fluorophore** | **Vendor** | **Clone** | **Catalog No.** | **Dilution** | **Stain** |
| CD126 (IL-6Ra) | BV421 | BD Biosciences | M5 | 564163 | 50 | Surface |
| CD130 (gp130) | PE | BD Biosciences | AM64 | 555757 | 50 | Surface |
| pSTAT5 (Y694) | PE-Cy7 | BD Biosciences | 47/Stat5(pY694) | 560117 | 50 | Intracellular |

| Mouse T Cell Activation/Exhaustion Panel | | | | | | |
|---|---|---|---|---|---|---|
| **Marker** | **Fluorophore** | **Vendor** | **Clone** | **Catalog No.** | **Dilution** | **Stain** |
| CD16/CD32 (FcBlock) | N/A | BD Biosciences | 2.4G2 | 553142 | 300 | Surface |
| CD44 | BUV395 | BD Biosciences | IM7 | 740215 | 400 | Surface |
| B220/CD45R | BUV496 | BD Biosciences | RA3-6B2 | 612950 | 150 | Surface |
| CD4 | BUV563 | BD Biosciences | GK1.5 | 741083 | 800 | Surface |

| | | | | | | |
|---|---|---|---|---|---|---|
| CD127 | BUV737 | BD Biosciences | SB/199 | 612841 | 100 | Surface |
| CD8a | BUV805 | BD Biosciences | 53-6.7 | 612898 | 1800 | Surface |
| Granzyme B | BV421 | BD Biosciences | GB11 | 563389 | 100 | Intracellular |
| Ly108 | Pacific Blue | BioLegend | 330-AJ | 134608 | 100 | Surface |
| CD69 | BV480 | BD Biosciences | H1.2F3 | 746813 | 200 | Surface |
| CD3 | BV510 | BD Biosciences | 17A2 | 740147 | 100 | Surface |
| CD45 | BV570 | BioLegend | 30-F11 | 103136 | 200 | Surface |
| TIM-3 | BV605 | BioLegend | RMT3-23 | 119721 | 100 | Surface |
| KLRG-1 | BV650 | BD Biosciences | 2F1 | 740553 | 100 | Surface |
| CD39 | SB702 | Invitrogen | 24DMS1 | 67-0391-82 | 100 | Surface |
| T-bet | BV786 | BD Biosciences | O4-46 | 56141 | 25 | Intracellular |
| CD73 | FITC | BioLegend | Ty/11.8 | 127220 | 100 | Surface |
| CXCR5 (CD185) | BB700 | BD Biosciences | 2G8 | 566493 | 50 | Surface |
| TCF-7/TCF-1 | PE | BD Biosciences | S33-966 | 564217 | 100 | Intracellular |
| CD62L | PE-CF594 | BD Biosciences | MEL-14 | 562404 | 800 | Surface |
| Ly6C | PE-Cy5.5 | Invitrogen | RB6-8C5 | 35-5931-80 | 400 | Surface |
| Eomes | PE-Cy7 | Invitrogen | Dan11mag | 25-4875-82 | 200 | Intracellular |
| TOX | APC | Miltenyi Biotec | REA473 | 130-118-335 | 100 | Intracellular |
| Ki-67 | AF700 | BD Biosciences | B56 | 561277 | 3000 | Intracellular |
| Zombie NIR | Viability | BioLegend | N/A | 423106 | 3000 | Surface |
| PD-1 | APC-Fire 750 | BioLegend | 29F.1A12 | 135239 | 600 | Surface |

| **Mouse Intracellular Cytokine Detection** | | | | | | |
|---|---|---|---|---|---|---|
| **Marker** | **Fluorophore** | **Vendor** | **Clone** | **Catalog No.** | **Dilution** | **Stain** |
| CD16/CD32 (FcBlock) | N/A | BD Biosciences | 2.4G2 | 553142 | 300 | Surface |
| CD44 | BUV395 | BD Biosciences | IM7 | 740215 | 400 | Surface |
| CD4 | BUV563 | BD Biosciences | GK1.5 | 741083 | 1600/1600 | Surface/Intracellular |
| IL-17A | BUV737 | Invitrogen | eBio17B7 | 367-7177-82 | 50 | Intracellular |
| CD8a | BUV805 | BD Biosciences | 53-6.7 | 612898 | 3600/3600 | Surface/Intracellular |
| Granzyme B | BV421 | BD Biosciences | GB11 | 563389 | 100 | Intracellular |
| RORgt | BV480 | BD Biosciences | Q31-378 | 567176 | 100 | Intracellular |
| CD45 | BV570 | BioLegend | 30-F11 | 103136 | 200 | Surface |
| IFN-γ | BV650 | BioLegend | XMG1.2 | 505832 | 100 | Intracellular |
| CD39 | SB702 | Invitrogen | 24DMS1 | 67-0391-82 | 100 | Surface |
| T-bet | BV786 | BD Biosciences | O4-46 | 56141 | 25 | Intracellular |
| CD62L | BB515 | BD Biosciences | MEL-14 | 565261 | 200 | Surface |
| CD19 | Biotinylated | BioLegend | 6D5 | 115503 | 600 | Surface |
| CD11b | Biotinylated | BioLegend | M1/70 | 101203 | 600 | Surface |
| CD11c | Biotinylated | BioLegend | N418 | 117303 | 600 | Surface |
| NK-1.1 | Biotinylated | BioLegend | PK136 | 108703 | 600 | Surface |

| | | | | | | |
|---|---|---|---|---|---|---|
| Gr-1 (Ly-6G/Ly-6C) | Biotinylated | BioLegend | RB6-8C5 | 108403 | 600 | Surface |
| Ep-Cam | Biotinylated | BioLegend | G8.8 | 118203 | 600 | Surface |
| Streptavidin | PerCP | BioLegend | NA | 405213 | 900 | Surface |
| IL-2 | PE | BioLegend | JES6-5H4 | 503808 | 200 | Intracellular |
| IL-13 | PE-eFluor 610 | Invitrogen | eBio13A | 61-7133-82 | 250 | Intracellular |
| CD3 | PE-Cy7 | BioLegend | 17A2 | 100220 | 400/400 | Surface/Intracellular |
| TNF-α | AF647 | BioLegend | MP6-XT22 | 506314 | 100 | Intracellular |
| Zombie NIR | Viability | BioLegend | N/A | 423106 | 3000 | Surface |
| CXCR3 (CD183) | APC-Fire 750 | BioLegend | CXCR3-173 | 126539 | 150 | Surface |

2474 **Table S6. Additional key reagents/resources.**

| Buffers/Media | Source | Catalog No. |
|---|---|---|
| DPBS (1X) without Ca++ & Mg++ | Penn Cell Center | MT21-031-CM |
| Corning® 500 mL RPMI 1640 | Corning | 10-040-CV |
| GemCell™ U.S. Origin Fetal Bovine Serum | GeminiBio | 100-500 |
| PENICILLIN STREPTOMYCIN SOL | Penn Cell Center | 15140122 |
| L-Glutamine - 200mM (100X) | Penn Cell Center | MT25-005-CI |
| HEPES Buffer, pH 7.4, 1M, TC tested | Penn Cell Center | 1138 |
| Gibco™ MEM Non-Essential Amino Acids Solution (100X) | Fisher Scientific | 11-140-050 |
| Sodium Pyruvate - 100X 11g/l 100mM | Penn Cell Center | 1175 |
| 2-Mercaptoethanol | Sigma-Aldrich | M6250-10ML |
| UltraPure™ 0.5M EDTA, pH 8.0 | ThermoFisher Scientific | 15575020 |
| eBioscience™ Foxp3 / Transcription Factor Staining Buffer Set | ThermoFisher Scientific | 00-5523-00 |
| BD Horizon™ Brilliant Stain Buffer | BD Biosciences | 566349 |
| ACK Lysing Buffer | ThermoFisher Scientific | A1049201 |

| Biologics | Source | Catalog No. |
|---|---|---|
| BD GolgiStop™ Protein Transport Inhibitor (Containing Monensin) | BD Biosciences | 554724 |
| BD GolgiPlug™ Protein Transport Inhibitor (Containing Brefeldin A) | BD Biosciences | 555029 |
| Ultra-LEAF™ Purified anti-human CD3 Antibody (Clone = OKT3; 1mg) | Biolegend | 317326 |
| Ultra-LEAF™ Purified anti-human CD28 Antibody (Clone = CD28.2; 1mg) | Biolegend | 302934 |
| Human IL-2 IS, premium grade | Miltenyi Biotec | 130-097-746 |
| Human IL-6 Recombinant Protein, PeproTech® | ThermoFisher Scientific | 200-06-100UG |
| Recombinant Human IL-10 Protein | R&D Systems | 217-IL-010 |
| Human IL-21 Recombinant Protein, PeproTech® | ThermoFisher Scientific | 200-21-10UG |
| Recombinant Human IL-27 Protein | R&D Systems | 2526-IL-010 |
| Human IFN-α2a, research grade | Miltenyi Biotec | 130-093-873 |
| Ultra-LEAF™ Purified anti-mouse CD3ε (Clone = 145-2C11; 1mg) | Biolegend | 100340 |
| Ultra-LEAF™ Purified anti-mouse CD28 Antibody (Clone = 37.51; 1mg) | Biolegend | 102116 |
| InVivoMAb anti-mouse CD3ε (Clone = 145-2C11) | BioXcell | BE0001-1 |
| nVivoMAb anti-mouse CD28 (Clone = 37.51) | BioXcell | BE0015-1 |
| Recombinant Mouse IL-27 (NS0-expressed) Protein | R&D Systems | 2799-ML-010 |
| InVivoMAb anti-mouse IL-4 (Clone = BE0045) | BioXcell | BE0045 |
| InVivoMAb anti-mouse IFNγ (Clone = XMG1.2) | BioXcell | BE0055 |
| Phorbol 12-myristate 13-acetate | Sigma-Aldrich | P1585-1MG |
| Ionomycin calcium salt | EMD Millipore | I0634-1MG |
| Adenosine 5′-triphosphate disodium salt hydrate | Sigma-Aldrich | A6419-1G |
| 2-Chloroadenosine | Sigma-Aldrich | C5134-10MG |
| STAT3 Inhibitor V, Stattic | EMD Millipore | 573099-25MG |

| Additional Flow Cytometry Reagents | | |
|---|---|---|
| SpectroFlo® QC Beads | Cytek | SKU B7-10001 |
| UltraComp eBeads | ThermoFisher Scientific | 01-2222-42 |

| Plasmid Reagent | Source |
|---|---|
| MigR1K | Krummel Lab |
| Stat3-203_MigR1K | Genscript |
| STAT3-203_Q344H_MigR1K | Genscript |
| pCL_Eco | Abbas Lab |

| Prokaryotic Cells | Source | Catalog No. |
|---|---|---|
| 5-alpha Competent E. coli (High Efficiency) DH5a | New England Biolabs | C2987H |
| SOC Outgrowth Medium | New England Biolabs | B9020S |

| Eukaryotic Cell Lines | Source | Catalog No. |
|---|---|---|
| 293T (CRL-3216) | ATCC | CRL-3216 |

| Other Reagents | Source | Catalog No. |
|---|---|---|
| Miller Luria Broth (LB):<br>2.5% w/v, autoclaved and supplemented with ampicillin (100ug/ml) | ThermoFisher Scientific | 12795027 |
| Lipofectamine 2000 | ThermoFisher Scientific | 11668027 |
| Dreamtaq Green PCR Master Mix (2x) | ThermoFisher Scientific | K1081 |

| PCR Primers | |
|---|---|
| STAT3 flx (Forward) | 5'- GAA GGC AGG TCT CTC TGG TG |
| STAT3 flx (reverse) | 5'- AGG CTG CCA ACA GCC ACT GCC |
| Cre Primer 1 (Forward) | 5'- GCT AAG GAT GAC TCT GGT CA |
| Cre Primer 2 (Reverse) | 5'- CTA ATC GCC ATC TTC CAG CA |
